# Netrin G1 promotes pancreatic tumorigenesis through cancer associated fibroblast driven nutritional support and immunosuppression

**DOI:** 10.1101/330209

**Authors:** Ralph Francescone, Débora Barbosa Vendramini-Costa, Janusz Franco-Barraza, Jessica Wagner, Alexander Muir, Allison N. Lau, Linara Gabitova, Tatiana Pazina, Sapna Gupta, Tiffany Luong, Neelima Shah, Dustin Rollins, Ruchi Malik, Roshan Thapa, Diana Restifo, Yan Zhou, Kathy Q. Cai, Harvey H. Hensley, Yinfei Tan, Warren D. Kruger, Karthik Devarajan, Siddharth Balachandran, Andres J. Klein-Szanto, Huamin Wang, Wafik S. El-Deiry, Matthew G. Vander Heiden, Suraj Peri, Kerry S. Campbell, Igor Astsaturov, Edna Cukierman

## Abstract

Pancreatic ductal adenocarcinoma (PDAC) has a poor 5-year survival rate and lacks effective therapeutics. Therefore, it is of paramount importance to identify new targets. Using multi-plex data from patient tissue, three-dimensional co-culturing *in vitro* assays, and orthotopic murine models, we identified Netrin G1 (NetG1) as a promoter of PDAC tumorigenesis. NetG1^+^ cancer-associated fibroblasts (CAFs) supported PDAC survival, through a NetG1 mediated effect on glutamate/glutamine metabolism. NetG1^+^ CAFs were intrinsically immunosuppressive and inhibited NK cell mediated killing of tumor cells. These pro-tumor functions were controlled by a signaling circuit downstream to NetG1, which was comprised of AKT/4E-BP1, p38/FRA1, vesicular glutamate transporter 1, and glutamine synthetase. Finally blocking NetG1 with a neutralizing antibody stunted *in vivo* tumorigenesis, suggesting NetG1 as potential target in PDAC.

**Significance:** PDAC is a devastating disease lacking effective therapies. A major hallmark of PDAC is desmoplasia, characterized by the expansion of CAFs and their extracellular matrix, creating a unique microenvironment that limits blood-supplied nutrition and is highly immunosuppressive. A better understanding of the role of CAFs in PDAC may lead to the identification of new targets for therapeutic intervention. Here, we uncovered roles for NetG1 in CAFs to promote tumorigenesis. NetG1 was important for two major CAF functions: the metabolic support of PDAC cells and the intrinsic immunosuppressive capacity of CAFs. Our results helped clarify the role that CAFs play in PDAC, by defining CAF phenotypes through NetG1 expression. Moreover, we established a link between CAF driven metabolism and their intrinsic immunosuppressive capacity, and identified a signaling circuit that governs NetG1 functions. Finally, we demonstrated the therapeutic potential of inhibiting NetG1 *in vivo* by limiting tumorigenesis in mice with a neutralizing antibody, illustrating that targeting stromal NetG1 could be an attractive therapeutic approach.

## Introduction

Pancreatic cancer is projected to become the 2^nd^ leading cause of cancer related deaths by 2030 [1], due to its abysmal 5 year survival rate [2]. The most common form of pancreatic cancer is pancreatic ductal adenocarcinoma (PDAC), to which treatments are often refractory, and where diagnosis is often made at advanced stages of the disease [3], making few patients eligible for surgical interventions [4]. Therefore, there is an urgent need to develop new diagnostic and therapeutic approaches.

PDAC has a unique microenvironment that consists of a fibrous expansion known as desmoplasia, characterized by the deposition of an abundant extracellular matrix (ECM) by cancer associated fibroblasts (CAFs) [5]. Studies have demonstrated that CAFs limit the efficacy of chemotherapies [6–8], promote PDAC progression [9, 10], and correlate with poor prognosis [11]. While there is a clear role for CAFs in facilitating PDAC progression, there is also evidence that CAFs can be tumor restrictive, as complete ablation of fibroblasts from the tumor microenvironment accelerated PDAC progression [12, 13] and provided no added patient benefit [14]. Thus, the role of CAFs in PDAC development and progression is incompletely understood.

An important consequence of desmoplasia is the generation of substantial interstitial pressure that contributes to the collapse of blood vessels [15], resulting in a nutrient deprived [16] and hypoxic microenvironment [17]. Further, *Kras* driven cancers like PDAC, where ∼90% of tumors have mutant *Kras*, have been suggested to become “addicted” to exogenous sources of glutamine as a main carbon supply [18]. As such, PDAC cells make use of metabolic pathways to promote tumor growth and survival [18–20]. By taking advantage of CAF driven metabolic support [21], CAFs supply PDAC cells with key nutrients [22]. Therefore, further exploration of these mechanisms, aiming to disrupt tumor-stromal crosstalk, is warranted.

In addition to the metabolic roles, CAFs are also at the center of the immunosuppressive PDAC microenvironment [23–26]. CAFs exert immunosuppressive effects through direct modulation of immune cell function via cytokine secretion [27], exclusion of anti-tumor immune cells from the tumor [28], and/or recruitment of immunosuppressive immune cells to the tumor [29]. Intriguingly, recent studies have reported a functional link between metabolism and immune cell function, which has suggested that anti-tumor immune cells rely on a constant supply of nutrients to perform their functions [30, 31]. In the cancer context, key metabolites—glucose, arginine, glutamine, and tryptophan—are depleted, with a concomitant increase of immunosuppressive waste products, such as lactate, inhibiting anti-tumor immune cell function [30, 32, 33]. In PDAC, the immunosuppressive environment created by CAFs, combined with the nutrient poor milieu, build a hostile environment that inhibits anti-tumor immune cell function. Thus, means to simultaneously revert these linked features of the PDAC microenvironment would be therapeutically beneficial.

In this study we sought to clarify the roles of CAFs in PDAC tumorigenesis. We uncovered up-regulation of the glutamatergic pre-synaptic protein, Netrin G1 (NetG1) [34, 35], in CAFs. Of note, we also found expression of Netrin G1 Ligand (NGL-1), the sole known post-synaptic binding partner of NetG1 [36], in PDAC cells. In fact, we saw that NetG1 was particularly overexpressed in human PDAC tissue compared to samples from normal human pancreas and that fibroblastic NetG1 expression correlated inversely with patient survival. We demonstrated that NetG1 in CAFs and NGL-1 in tumor cells enhanced tumorigenesis, as ablation of either protein reduced tumor burden in mice. Functionally, NetG1 was responsible for maintaining high levels of glutamate, glutamine, and cytokine release by CAFs which allowed PDAC cells to survive in low nutrient conditions and reduced death induced by NK cells. Mechanistically, NetG1 controlled a signaling circuit comprised of glutamine synthetase and vesicular glutamate transporter 1, p38/FRA-1 and AKT/4E-BP1, and systematic inhibition of each component of the pathway revealed their roles in the pro-tumor metabolic and immunosuppressive functions of CAFs. Finally, the therapeutic applicability of targeting NetG1 *in vivo* was demonstrated, as an anti NetG1 neutralizing monoclonal antibody inhibited tumorigenesis in a murine model of PDAC. Thus, we demonstrate the importance of considering the stroma as a viable therapeutic option in PDAC.

## Results

### NetG1 is upregulated in CAFs compared to patient matched tumor adjacent fibroblasts

To gain insight into the mechanism by which naive fibroblasts are activated into cancer associated fibroblasts (CAFs), and knowing that our cell-derived extracellular matrix (ECM) based culturing system assures maintenance of *in vivo*-like fibroblastic phenotypes [37, 38], we performed a transcriptomic analysis comparing two sets of patient matched tumor adjacent fibroblasts (TA) and CAFs **(Fig 1A, Sup Table 1)**. We identified the most differentially regulated genes between patient matched TA and CAFs (117 genes, p<0.01, log fold change; **Sup Table 1**). To characterize the underlying molecular pathways and biological processes, we performed Gene Set Enrichment Analysis (GSEA) and found significant enrichment (Benjamini-Hochberg False Discovery Rate < 0.25) of several pathways previously found to be associated with the transition from normal fibroblasts to CAFs, including upregulation of integrin signaling, focal adhesions, actin cytoskeleton, and extracellular matrix (ECM) organization (**Fig 1B-C, Sup Table 1**). Interestingly, results also revealed an upregulation of genes involved with axonal guidance and neurite outgrowth in CAFs compared to TA. The second most upregulated gene overall and top hit among the neural genes was *NTNG1*, encoding NetG1 [34], a synapse stabilizing protein known for its involvement in axonal communication. We also noticed in CAFs a negative enrichment of pathways associated with antigen presentation (KEGG Systemic Lupus Erythematosus; FDR < 0.25, Normalized Enrichment Score −2.66) and interferon signaling (Reactome Interferon Signaling; FDR < 0.25; NES = −2.66) (**Fig 1D, Sup Table 1**). Overall, this transcriptomic data confirmed that our ECM cultured CAFs indeed upregulate known pathways previously linked to fibroblast activation in cancer, as well as unveil a novel association with neural gene expression.

**Figure 1.**
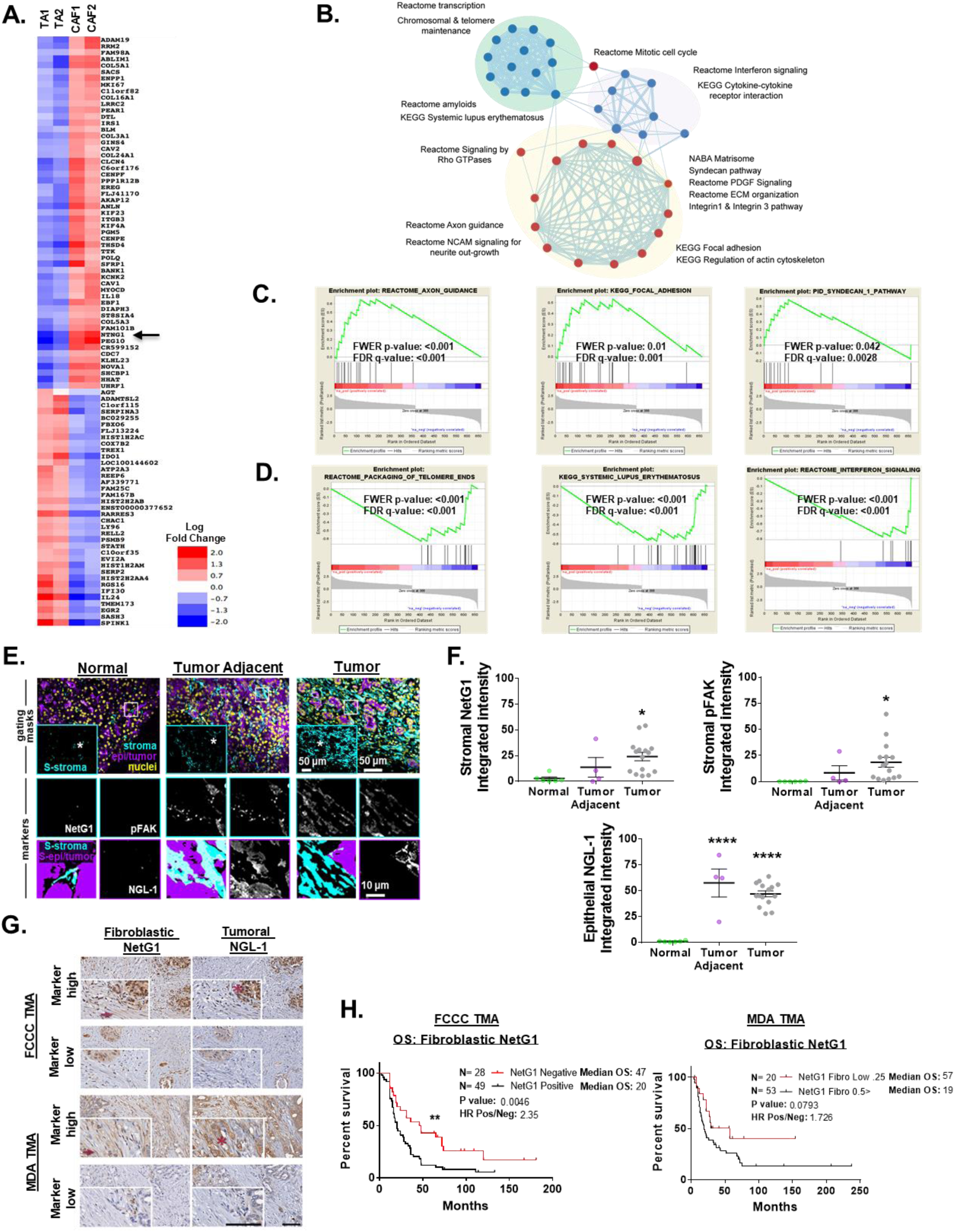
High NetG1 expression in CAFs inversely correlates with patient survival. **A.** Heatmap representing increased (red) or decreased (blue) gene expression levels (log fold change) between patient derived tumor adjacent fibroblasts (TA) and CAFs. Position of NetG1 (*NTNG1*) in the heatmap is highlighted with a black arrow. Top 117 differentially expressed genes were chosen by p-value (<0.01). **B.** Pathways that are significantly enriched between these two cell types are represented as network map. Each node (circle) represent a pathway while the edges (lines) connecting nodes show shared genes between pathways with thickness of edge corresponds to degree of sharing. Color of node indicate positive (red) or negative (blue) enrichment in CAF cells. Enrichment of select pathways are shown as GSEA enrichment plots with positive (**C**) and negative (**D**) enrichment in CAFs. **E.** Representative images of simultaneous multi-channel immunofluorescent (SMI) approach, performed on formalin fixed paraffin embedded tissue samples corresponding either to normal pancreatic tissue (normal human pancreas), normal tissue adjacent to PDAC tumor (tumor adjacent), or PDAC tumor tissue (tumor) from patient/donor surgical samples. Upper row panels show merged and pseudo-colored images corresponding to the three locations used as “gating masks.” These masks correspond to pan-cytokeratins positive areas in purple demarking epithelial/tumoral compartment, vimentin regions in cyan and DRAQ5 for nuclei in yellow. The SMIA-CUKIE algorithm (described in Material and Methods) was instructed to render an intersection “mask” image corresponding to vimentin positive epithelial/tumoral negative areas (S-stroma in upper panels inserts), whereas epi/tumor positive masks omitted all vimentin positive pixels. The two above-mentioned masks were used as areas to generate the images shown in the medium and lower rows, which are magnified images corresponding to the regions of interest highlighted in upper row images within white squares and asterisks. Medium row show “markers” in stromal areas corresponding to NetG1 and pFAK, while lower row shows NGL-1 at tumor areas. Lower left corner panels indicate the algorithm generated area “masks” used (S-epi/tumor; S/stroma). Representative scale bars are provided for each magnification **F.** Graphs depict integrated intensities of stromal NetG1, stromal pFAK, or epithelial NGL-1 staining in normal (N= 6), tumor adjacent (N= 4), or tumor (N= 15) pancreatic tissue. *Compared to normal tissue. One-Way ANOVA, Dunnett’s multiple comparison test. *p<0.05, ****p<0.0001. **G.** Representative images of patient-matched pancreatic cancer tissue evaluated in TMAs obtained from FCCC and MDA. Top row panels show high stromal NG1 and tumoral NGL-1 expression in patients with poor survival, while bottom row images illustrate tissue examples with low levels of mentioned markers, corresponding with patients with extended survival. Inserts in each image show magnified regions with examples of stromal cells pointed by dark red arrows; tumor areas are highlighted by dark red asterisks. Scale bars represent 100 μm. **H.** Kaplan-Meier plots depicting overall survival (OS) of PDAC patients from two independent TMAs (**LEFT:** FCCC; N= 80) and (**RIGHT:** MDA; N=143), stratified by immunohistological scores of fibroblastic NetG1. Log rank test was used to determine statistical significance. **p<0.01.

### NetG1 in CAFs, and its binding partner NGL-1 in tumor cells, are overexpressed in PDAC

NetG1 is present on pre-synaptic neurons, and it is known to stabilize excitatory glutamatergic synapses by interacting with its only known receptor on post-synaptic cells, Netrin G1 Ligand (NGL-1) [35, 36]. To assess the clinical relevance of NetG1, we performed Simultaneous Multichannel Immunofluorescence (SMI) [38] on 6 normal, 4 tumor adjacent (TA) and 15 tumor pancreatic tissue samples to discern NetG1 and NGL-1 expression patterns in distinct subsets of cells. NetG1 was upregulated in the stromal compartment (pan-cytokeratin^-^, vimentin^+^ areas) in tumor tissue compared to normal pancreatic tissue from individuals without cancer (**Fig 1E-F**). Moreover, levels of tyrosine 397 phosphorylated focal adhesion kinase (pFAK) followed a similar pattern of increased expression in tumor tissue compared to normal tissue, in line with our established pro-tumor CAF signature [38] (**Fig 1E-F**). A strong correlation between stromal NetG1 expression and stromal pFAK expression was observed across all tissues (R^2^ = 0.8046, p<0.0001) (**S1A**). Further, NGL-1 was upregulated in the epithelial compartment (pan-cytokeratin^+^ areas) of both tumor adjacent and tumor tissue (**Fig 1E-F**), with a weak correlation between stromal NetG1 and epithelial NGL-1 expressions across all tissues (R^2^ = 0.2359, p<0.0001) (**S1A**), suggesting that ectopic expression of NetG1 and NGL-1 may be associated with PDAC pathology. Comparing 4 TA and 5 CAFs isolated from patient tissue, we confirmed that CAFs upregulated NetG1 protein expression (∼2.5 fold increase) (**S1B**). Additionally, we observed the expression of NGL-1 in 5 patient derived cell lines from PDX models (**S1C**), as well as upregulated expression in tumor cell lines derived from an isogenic cell line model, hTERT-HPNE E6/E7 (referred to as E6/E7) and hTERT-HPNE E6/E7/K-RasG12D (referred to as PDACc), compared to their immortalized epithelial cell counterpart (hTERT-HPNE; referred to as hTERT) [39] (**S1D**). Collectively, NetG1 and its binding partner NGL-1 were upregulated in cancer compared to normal tissue in the pancreas, and segregated into distinct cellular compartments (stromal vs. epithelial).

### Fibroblastic NetG1 expression correlates with worse overall survival in PDAC patients

Using the cBioportal database [40, 41], we queried both NetG1 (*NTNG1)* and NGL-1 (*LRRC4C*) mRNA expression across 31 cancer types and found that PDAC was among the highest expressors of both genes (**S2A, arrows)**. Additionally, we stratified *NTNG1* and *LRRC4C* by clinical subtype according to three independent datasets [42, 43]; (TCGA, [44]) (**S2B-E**). We found that in two out of three datasets there was a trend towards elevated *NTNG1* and *LRRC4C* in the basal classification, which presents with the worst prognosis of all PDAC subtypes [43].

Moreover, we used the datasets from TCGA and Puleo et al. and stratified patients into 4 groups based on *NTNG1* and *LRRC4C* expression: *LRRC4C* high/*NTNG1* high (UP-UP), (*LRRC4C* high/*NTNG1* low (UP-DN), *LRRC4C* low/*NTNG1* high (DN-UP), and *LRRC4C* low/*NTNG1* low (DN-DN) and examined if these groups correlated with overall survival (OS) (**S2F**). Although we did not see a significant association between expression and OS, we found a marginal association between relapse free survival (RFS) and co-expression of *LRRC4C* and *NTNG1* (hazard ratio (HR) of 9.45 (1.27-70.27; P = 0.0074; FDR > 0.5) (**S2G**). These inconclusive results are likely due to the intermingling of stromal and epithelial RNA that is inherent in cancers with a high degree of stromal expansion, such as PDAC [44], highlighting the importance of carefully dissecting the contributions of the stromal and epithelial compartments of the tumor microenvironment.

Thus, to further address the clinical significance of NetG1 and NGL-1, we performed immunohistochemical staining of NetG1 and NGL-1 on two independent tissue microarrays (TMA; **Fig 1G, Sup Tables 2-3**). Protein expression of fibroblastic NetG1 and tumoral NGL-1 were scored blindly by a pathologist (0-4 scale; **Sup Tables 4-5**). Strikingly, fibroblastic NetG1 expression correlated inversely with patient overall survival (p<0.01; median OS NetG1^+^ vs NetG1^-^: 20 vs 47 months; hazard ratio: 2.35) in the Fox Chase Cancer Center (FCCC) TMA, while there was a similar trend in the MD Anderson (MDA) TMA (p=0.0793; median OS NetG1^High^ vs NetG1^Low^: 19 vs 57 months; hazard ratio: 1.73) (**Fig 1H**). Tumoral NGL-1 expression was not predictive of survival (**S1E**). This was likely due to the fact that nearly every patient stained positive for tumoral expression of NGL-1 (72/77 FCCC; 129/135 MDA), and the acquisition of NGL-1 expression may be an early event in tumorigenesis. Overall, fibroblastic NetG1 expression resulted in worse overall survival for PDAC patients, suggesting that NetG1 could be a useful prognostic marker in PDAC.

### RNAseq analysis reveals that ablation of NetG1 in CAFs results in a normalized stromal gene expression signature

CAFs are known to function in an ECM dependent manner and regulate tumorigenesis in a number of ways [5]. Hence, we relied on our well-established 3D cell-derived matrix culturing system, referred to as “3D” for simplicity, which simulates the physiological as well as pathologic tumor microenvironment [37, 38, 45], to dissect the functions of NetG1 in more detail *in vitro*. To this end, we first generated NetG1 knockout (KO) CAFs through CRISPR/Cas9 (**S3A**), from one of our patient derived CAF lines that displayed high levels of NetG1 mRNA and protein expression compared to tumor adjacent fibroblasts (TA) (**S3B, S1B**). To gain insight into global changes in gene expression, we performed RNAseq analysis comparing control (CON) CAFs and NetG1 KO CAFs (**S3C, Sup Table 6).** Pathway and gene enrichment analysis identified downregulation of genes associated with many critical pathways, known as being upregulated in CAFs, in our microarray analysis (**Fig 1**), including: axon guidance, focal adhesion, ECM and TGF-β signaling, and inflammatory responses (**S3D-E, Sup Table 6**). Thus, taken together, these results suggest a reversion of gene expression profiles linked with known pro-tumor CAF functions upon loss of NetG1 expression.

### Ablation of NetG1 does not influence myofibroblastic features of CAFs, but reverses pro-tumorigenic CAF traits

Next, we followed NetG1 expression during ECM production *in vitro*, as this is a major function of CAFs in the *in vivo* tumor microenvironment [5]. Importantly, NetG1 protein expression in CAFs was significantly elevated in 3D compared to 2D culturing conditions (**S4A**). NetG1 expression was maintained throughout ECM production (**S4B**). We then assessed the desmoplastic phenotype of CON or NetG1 KO CAFs during ECM production, according to our previous study [38]. Interestingly, NetG1 KO CAFs did not have altered canonical myofibroblastic features, as determined by ECM fiber (fibronectin) alignment and levels/localization of alpha smooth muscle actin (α-SMA) (**S4C-D**). However, NetG1 KO CAFs displayed reduced levels of active α_5_β_1_-integrin and pFAK, indicative of a CAF phenotype that was associated with delayed recurrence following surgery in PDAC patients (**S4E-F**) [38], consistent with the correlation seen in human tissue samples (**S1A**). Thus, deletion of NetG1 does not alter canonical myofibroblastic features of CAFs, but attenuates the expression of known pro-tumor CAF markers. Thus, we hypothesized that deletion of NetG1 would revert pro-tumorigenic features of CAFs and potentially stunt tumorigenesis *in vivo*.

### Ablation of NetG1 in CAFs and NGL-1 in PDAC cells significantly stunts tumorigenesis in orthotopic murine models of PDAC

To test the impact of NetG1 and NGL-1 in pancreatic cancer *in vivo*, we performed multiple murine models of PDAC, in which we orthotopically injected CAFs and PDAC cells, and observed tumor progression and burden. In the first model, we co-injected human CON or NetG1 KO CAFs with RFP^+^ CON or NGL-1 KO PDACc cells (**S4G**), at a CAF:PDAC ratio of 3:1, into the pancreas of severely compromised immunodeficient (SCID) mice and followed tumorigenesis over 1 month. Animals co-injected with CON CAFs and CON PDACc cells had a substantially worse tumor burden, with more obvious macroscopic RFP^+^ coverage of the pancreas (**Fig 2A**), as well as greater tumor weight (∼2-3 fold) **Fig 2B**), pancreas weight (∼2-3 fold) (**Fig 2C**), and tumor area (**Fig 2D**), than all other conditions. Importantly, mice co-injected with NetG1 KO CAFs and CON or NGL-1 KO PDACc cells developed tumors to a similar extent as mice injected with CON or NGL-1 KO tumor cells alone, suggesting that CAFs lacking NetG1 do not confer any benefit for tumorigenesis. As a control, mice were injected with the CON or NetG1 KO CAF lines alone, and those animals did not develop tumors. We also observed that mice injected with NGL-1 KO PDACc cells alone displayed a tendency for less tumor burden than CON PDACc cells, thus we proceeded to investigate this in more detail.

**Figure 2.**
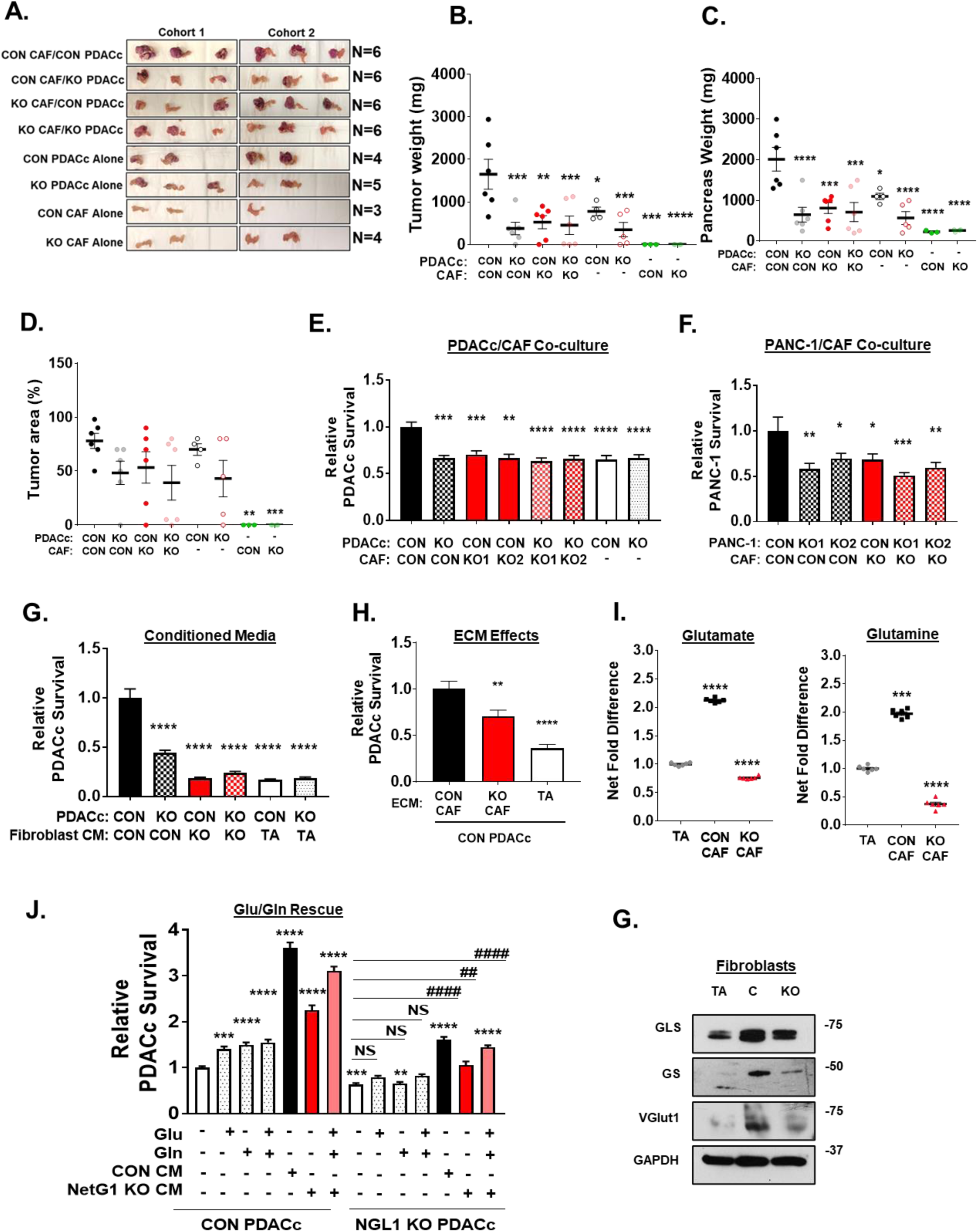
Ablation of NetG1 in CAFs stunts tumorigenesis in vivo and regulates Glu/Gln driven support of nutrient deprived PDAC cells. 7.5×10^5^ CON or NetG1 KO CAFs were orthotopically injected into the pancreas of SCID mice with RFP^+^ CON or NGL-1 KO PDACc cells (2.5×10^5^), at a 3:1 ratio. CAFs or PDACc cells injected alone served as controls. Mice were sacrificed 1 month after injection and tumorigenesis was assessed. **A.** Images of the isolated pancreata. B. Graph depicting relative tumor weight from each experimental group. **C.** Quantification of pancreas weights from each experimental group. **D.** Graph displaying the quantification of the % area of the pancreas that was classified as a tumor. **E**. RFP^+^ CON or NGL-1 KO PDACc (2×10^4^) were co-cultured in 3D with GFP^+^ CON or NetG1 KO CAFs (2×10^4^) (2 clones, KO1 and KO2) or alone in the absence of serum and Gln for 4 days followed by cell survival assessments. * compared to CON PDACc/CON CAF. **B.** Same assay as in (**A**), but with RFP^+^ CON or NGL-1 KO PANC-1 cells. * compared to CON PANC-1/CON CAF. **C.** RFP^+^ CON or NGL-1 KO PDACc (2×10^4^) were co-cultured in 3D with CM from CON, NetG1 KO CAFs, or tumor adjacent normal fibroblasts (TA) in SF/Gln free media, and PDACc survival was measured after 4 days. * compared to CON PDACc/CON CAF. **D.** RFP^+^ CON PDACc (2×10^4^) were grown in 3D in the indicated fibroblast derived ECMs alone in the absence of serum and Gln for 4 days and PDAC cell survival was measured. * compared to CON PDACc/CON CAF. **E.** Relative glutamate and glutamine levels in the CM of TA, CON or NetG1 KO CAFs. N= 6 biological replicates, all groups were compared to TA condition. **F.** RFP^+^ CON or NGL-1 KO PDACc (2×10^4^) were cultured alone in 3D under serum and Gln deprivation (-). Graph depicts relative PDACc survival after exposure to media alone, Glu, Gln, or CAF CM. Treatment groups consisted of Glu (150 µM) and Gln (25 µM) addbacks to determine if those amino acids alone could rescue PDAC cell survival in the absence of CAFs. CM from CON CAFs was used as a positive control for rescue. Note how amino acids alone partially rescue CON PDACc but not NGL-1 KO PDACc, which also benefit from CM media to a lesser extent than their CON counterparts. * compared to CON PDACc alone; # compared to KO PDACc alone. All Graphs: One-Way ANOVA, Dunnett’s multiple comparison test. *p<0.05, **p<0.01, ***p<0.001; ****p<0.0001**. G.** Representative western blots of glutaminase (GLS), glutamine synthetase (GS) and vesicular glutamate transporter 1 (VGlut1) in TA, CON CAFs (C), and NetG1 KO CAFs (KO). GAPDH was used as a loading control.

We first explored NetG1 expression in the stroma of LSL-Kras^G12D/+^; Pdx-1-Cre (KC) mice, which readily develop precancerous lesions known as PanINs (Pancreatic Intraepithelial Neoplasia) and eventually full blown invasive PDAC [46]. Using SMI, we observed that as KC mice age and develop PanINs, stromal NetG1 expression was detected in pancreatic tissue as early as 12 and 16 weeks, while undetectable in non-tumor bearing mice (**S5A**). We generated two murine PDAC cell lines, isolated and cloned from the more aggressive LSL-KRAS^G12D^;TP53^-/-^;PDX1-Cre (KPC) model [47], one with low NGL-1 expression that resulted in an average survival time of 7.5 weeks after orthotopic injection into C57BL/6 mice (KPC4B), and one with high expression of NGL-1 that resulted in an average survival time of 3.5 weeks after orthotopic injection (KPC3) (**S5B**). These data indicated that, similarly to human PDAC, NetG1/NGL-1 are present in PDAC in mice. Using CRISPR/Cas9, we generated two NGL-1 KO murine PDAC cell lines and one control line from the aggressive KPC3 cells (**S5C**) and injected them orthotopically into C57BL/6 mice (CON, KO1, KO2). Mice injected with CON KPC3 cells (CON group) had a tumor incidence of 100% (7/7), while the groups injected with NGL-1 KO1 and KO2 KPC3 cells (NGL-1 KO group) had an incidence of 22.2% (2/9) and 40% (4/10) with KO tumors displaying significantly less tumor weight (**S5D**), pancreas weight (**S5E**) and % tumor area (**S5F-G**) compared with CON group. In addition, representative MRI images revealed the expansion of the pancreas in CON group over 3 weeks, while the pancreas from the NGL-1 KO group seemingly remained at normal size over that time span (**S5H**). Moreover, tumors from the KO group had a trend towards being more differentiated than those from the CON group. An average of 79% of the cells from CON tumors were classified as poorly differentiated, compared to 48% in KO tumors (**S5I**), indicative of less aggressive tumors overall. Consequently, NGL-1 KO tumors also proliferated slower, as measured by Ki67 staining, and had higher rates of apoptosis, as measured by TUNEL staining (**S5J**). These results were in line with less Ki67 expression in human NGL-1 KO PDACc compared to CON PDACc cultured in 3D *in vitro* (**S4H**).

In order to evaluate the impact of the immune system in this model, orthotopic injections of CON or NGL-1 KO KPC3 cells into SCID (lacks T and B cells) and Nod SCID Gamma (NSG; lacks T, B, and NK cells) mice were performed and compared to immunocompetent mice (B6). All animals injected with CON cells presented more tumors than those injected with NGL-1 KO cells, independently of the immune backgrounds (**S5L**). This suggests that tumorigenesis, in relation to tumoral NGL-1, is not significantly affected by the immune system. Overall, the results demonstrated that both NetG1 in CAFs, and NGL-1 in tumor cells, promote PDAC tumorigenesis *in vivo*.

### NetG1 increase heterotypic cell-cell interactions between CAFs and PDAC

Because of NetG1’s known role in cell-cell adhesion in the brain, namely through its sole known receptor NGL-1, we tested the potential function of NetG1/NGL-1 in CAF-PDAC cell-cell interactions. Ablation of NetG1 in CAFs reduced cell engagement with control (CON) or NGL-1 KO PDACc by 72% and 79%, respectively (identified by the yellow regions resulted from the interaction between GFP+ CAFs and RFP+ PDACc) (**S6A-C**). Accordingly, CON or NGL-1 KO PDACc had increased motility (81% and 74%, respectively) when co-cultured with NetG1 KO CAFs in 3D (**S6D-E**). Unexpectedly, deletion of NGL-1 from PDACc was not sufficient to alter PDACc-CAF cell engagement or PDACc motility, signifying that there may be a redundant mechanism controlling PDAC-CAF engagement in the absence of NGL-1 in tumor cells and that NetG1 in CAFs is the driver of PDAC-CAF heterotypic interactions.

Because NetG1 in CAFs mediated PDAC-CAF interactions, we questioned if there was a further functional consequence of the increased heterotypic cell-cell interactions observed during co-culture. Since PDAC cells have been suggested to use numerous extracellular macromolecules as a source of energy [16, 48], and receive nutrients from CAFs [49, 50], we hypothesized that CAFs may provide resources to PDAC cells in a NetG1 dependent manner. To test this, we followed the exchange of GFP from GFP^+^ CAFs and RFP from RFP^+^ PDACc. Indeed, after 36 hours of co-culture, GFP was detected intracellularly in PDACc (**S6F**), demonstrating that PDACc received CAF derived material. Knockout of NetG1 in CAFs, or NGL-1 in PDACc, resulted in a 49%-61% reduction in GFP transfer to PDACc, compared to CON PDACc/CON CAF group (**S6G**). Therefore, while NGL-1 was dispensable for CAF-PDAC engagement, both NGL-1 and NetG1 were necessary for the ability of PDACc to receive material from CAFs, and for CAFs to provide material to PDACc.

### NetG1/NGL-1 axis protects PDAC cells from death under nutrient restriction

Since the PDAC microenvironment contains limited nutrients and CAFs can transfer material in a NetG1/NGL-1 dependent manner (**S6F-G**), we hypothesized that CAFs serve as a critical nutrient conduit and could rescue PDAC cells from nutrient deprivation. To this end, we co-cultured GFP^+^ CAFs with RFP^+^ PDACc in 3D under serum free/glutamine (Gln) free conditions and measured the survival of RFP^+^ PDACc over a 4 day time span. We observed that disruption of NetG1/NGL-1, by deletion of either NetG1 in CAFs or NGL-1 in PDAC cells, resulted in 30-37% decrease in PDACc survival, compared to CON PDACc/CON CAF group, similar to PDACc cultured alone (**Fig 2E**). These results were replicated using the pancreatic cancer cell line PANC-1 (CON or NGL-1 KO), with a 40-59% reduction in PANC-1 survival compared to CON/CON group (**Fig 2F, S4I**). To determine if CAF secreted factors alone were important for tumor cell survival, PDACc were cultured in 3D with conditioned media (CM) obtained from CON and NetG1 KO CAFs. We found that NetG1 KO CAF CM was 76-82% less efficient at supporting PDACc survival when compared to CON CAF CM, similarly to the CM from TA fibroblastic cells (81-83% of CON CM) (**Fig 2G**). To further assess the effect of NetG1 on the microenvironment generated by CAFs, we performed survival assays using ECMs produced by CON and NetG1 KO CAFs as substrates for cancer cells, and compared them to the ECM produced by TAs, which express less NetG1 than CAFs (**Fig 2H**). Accordingly, the microenvironment generated by NetG1 KO CAFs was less supportive compared to the CON ECM, with a 30% decrease in PDACc survival. The TA generated ECM was even less supportive, resulting in a decrease in PDACc survival of 65%. For experimental rigor, we also generated NetG1 KD CAFs, using CRISPRi, in two CAF lines (CAF and CAF2) and replicated the survival assay results (**S7A-C**). Collectively, these results suggest that both direct physical contact with CAFs, and the factors secreted by CAFs, including CM and ECM, support PDAC survival under nutritional deprivation, in a NetG1/NGL-1 dependent fashion.

### Alterations in Glutamate/Glutamine generation in NetG1 KO CAFs reduce PDAC cell survival

In an effort to gain insight into key factors that could be supporting PDAC survival under nutritional deprivation, we performed an amino acid screen comparing TA, CON CAFs, and NetG1 KO CAFs, during ECM production (**Sup Tables 4-6**). The greatest net change in amino acid secretion between TAs and CAFs was found to be in glutamate (Glu) and Gln, as CAFs secreted 2-fold more of these amino acids (**Fig 2I**). These results are in line with recent studies that suggest that the catabolism of Glu plays a role in PDAC and other cancers [18, 19, 51, 52]. Strikingly, we found that NetG1 KO CAF CM contained 64% less Glu and 81% less Gln, compared to CON CAFs (**Fig 2I**), and this was comparable in CAF2 (**S7D**). In order to determine the relative contributions of Glu and Gln to PDAC survival conferred by CON CAFs CM, we performed a Glu/Gln rescue assay, where the amounts of Glu/Gln produced by CON CAFs (150 µM and 25 µM respectively) were added to the depleted media and compared to the rescue provided by CON CAFs CM. Interestingly, while Glu or Gln alone or in combination promoted 40%, 49% or 55% more survival compared to deprived media alone, CON CAF CM increased CON PDACc survival by 3.5 fold, indicating that there are additional factors in the CM, besides Glu and Gln, contributing to PDAC survival (**Fig 2J**). Importantly, Glu/Gln were not able to rescue NGL-1 KO PDACc survival, suggesting a potential defect in Glu/Gln utilization in PDAC cells lacking NGL-1. However, supplementing back Glu/Gln at CON CAF CM levels to NetG1 KO CM was able to significantly rescue PDACc survival, although to a lesser degree in NGL-1 KO cells. These results suggest that adding Glu/Gln to factors secreted by NetG1 KO CAFs suffices to mimic CON CAF CM effects in rescuing PDACc survival under nutrient deprived conditions.

Guided by the clear differences in Glu/Gln levels upon KO of NetG1 in CAFs, we explored if glutaminase (GLS) and glutamine synthetase (GS) protein levels were altered in these cells. We detected expression of GLS and GS in TAs and CAFs (**Fig 2G**). While GLS levels were generally higher in CON and NetG1 KO CAFs than TAs, GS was significantly reduced in NetG1 KO CAFs, suggesting that NetG1 KO CAFs are compromised in their ability to generate Gln from Glu. Interestingly, NetG1 KO CAFs had reduced levels of vesicular glutamate transporter 1 (VGlut1), a major protein responsible for loading presynaptic vesicles with Glu, assuring synaptic Glu secretion, during neuronal communication [53] (**Fig 2G**). Thus, while CON and NetG1 KO CAFs expressed similar levels of GLS, NetG1 KO CAFs displayed decreased levels of VGlut1 and GS (also confirmed at the mRNA level, **S7E**), which could account for the reduced amounts of Gln produced and Glu released by NetG1 KO CAFs. These results were also observed in the NetG1 KD CAFs (**S7A**) and CAF2 (**S7F**). As expected from KRAS mutant cells, PDACc expressed lower levels of GS compared to CON CAFs, while maintaining GLS expression (**S4K**). Interestingly, knockout of NGL-1 in PDACc resulted in a reduction in GLS protein levels, suggesting an even greater metabolic dependence on CAFs or extracellular nutrients in the KO cells compared to CON PDACc. Additionally, NGL-1 expression in PDAC cells appears to be critical for the utilization of some extracellular factors, such as Gln and Glu, perhaps due to downregulation of glutamate receptor binding pathways, as suggested by TCGA NGL-1 stratification (**S4J**). Collectively, these results suggest that Gln and Glu derived from CAFs help support PDAC survival, under nutrient deprived conditions, and that this support depends on NetG1 expression in CAFs.

### Knockout of NetG1 in CAFs abrogates their immunosuppressive phenotype and permits NK cell anti-tumor function, partially through IL-15

Thus far, we noticed that NetG1 expression in CAFs was critical for their pro-tumor functions, including modulation of the ability of CAFs to provide tumor cells with critical nutrients for their survival (**Fig 2**). Moreover, the RNAseq analysis suggested that KO of NetG1 led to a normalization of CAFs, through the downregulation of pro-tumor fibroblastic pathways, including inflammatory signaling (**S3**). This led us to question if NetG1 is also a functional mediator of immunosuppression, another key CAF function [5]. Hence, we performed both traditional and multiplex (U-Plex) enzyme-linked immunosorbent assays (ELISAs) with TAs and CAFs cultured in 3D, to identify a cytokine profile that could define the immunosuppressive milieu that CAFs generate. We found that CON CAFs produce increased levels of GM-CSF, IL-1β, IL-8, CCL20/MIP3α and TGF-β compared to TAs (**Fig 3A**). Strikingly, ablation of NetG1 in CAFs significantly decreased protein levels of GM-CSF (40.6 fold), IL-6 (1.4 fold), IL-8 (13.3 fold), IL-1β (5.2 fold), CCL20/MIP3α (80 fold), and TGF-β (1.5 fold) compared to CON CAFs (**Fig 3A**), with a similar downregulation in NetG1 KD CAFs (**S8A**) and CAF2 (**S8B**). At the mRNA level, GM-CSF and TGF-β were correspondingly downregulated in NetG1 KO CAFs, with increases in IL-6 and IL-8 mRNA expression (**S8C**), suggesting post-translational regulation of IL-6 and IL-8. On the other hand, expression of IL-2, IL-12 p70, IL-10, IFN-γ, TNF-α, and IFN-β were all below the limit of detection in all three fibroblastic populations tested (data not shown). Interestingly, we observed that IL-15 protein levels, one of the most potent known NK cell activators, were up-regulated in both CON and NetG1 KO CAFs compared to TAs (4 and 2.9 fold, respectively; **Fig 3A**). Also, no differences were noted in mRNA expression of IL-15 when comparing CON CAFs to NetG1 KO CAFs (**S8C**). This suggested that NetG1 KO CAFs may generate a less immunosuppressive microenvironment than the one from CON CAFs, possibly allowing NK cells to more effectively kill PDAC cells, which could represent an important anti-tumor mechanism. Indeed, in our two TMA cohorts, the majority of patients presented with low NK cell infiltrates (represented by IHC of NKp46, **S9A**) (FCCC: 59/79 or 75%; MDA: 80/137 or 58%). Importantly, similarly to the median OS times observed in stromal NetG1 high patients (**Fig 1H**), patients with low NK cell infiltrate numbers had a median OS of 20.5 months and 21 months, in the FCCC and MDA TMAs respectively (**S9B, Sup Tables 4-5**). However, in patients that had more NK cell infiltration, there was a trend for a noticeable increase in survival, 46 and 33 months in the FCCC and MDA TMAs, respectively (p-values: 0.1213, 0.0546; HR: 0.65, 0.68).

**Figure 3.**
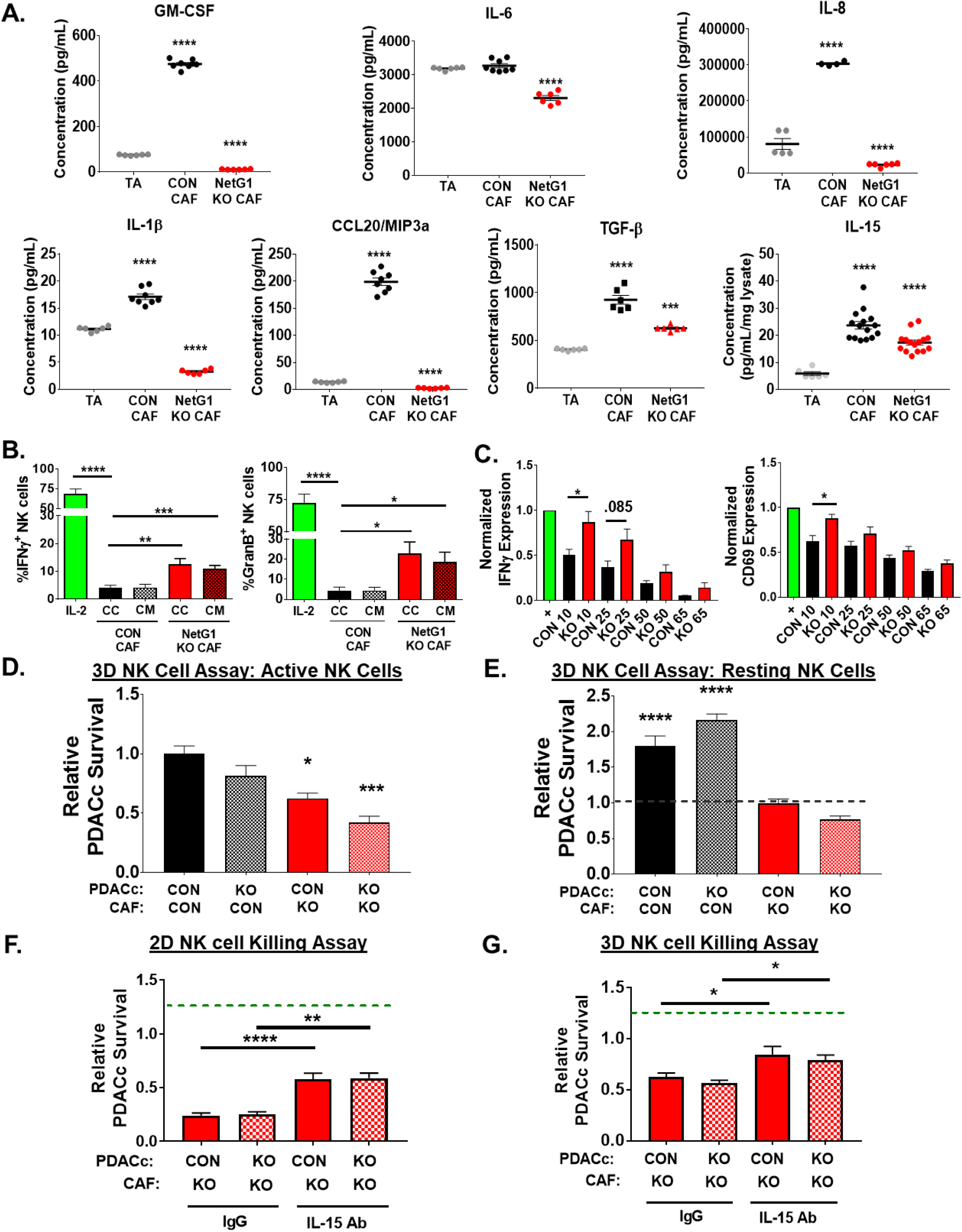
NetG1^+^ CAFs create an immunosuppressive microenvironment that protects PDAC cells from NK cell induced death. **A.** Quantification of U-Plex (multiplex ELISA; GM-CSF, IL-1β, CCL20, IL-6, IL-8) and ELISAs (IL-15, TGF-β) of assorted cytokines with immunomodulatory or immunoattractive potentials, detected in the CM of TA, CON CAFs, or NetG1 KO CAFs, growing in 3D. N= 6 biological replicates. * compared to TA. **B.** Quantification of the % of NK-92 cells positive for markers of activation (IFNγ and Granzyme B) determined by flow cytometry after IL-2 pre-activated NK-92 cells (8×10^4^) were in direct co-culture (CC) with CON or NetG1 KO CAFs (2×10^4^) or treated with their conditioned media (CM) for 16 hours. * compared to CON CAF CC. **C.** Primary NK cells (10^5^) were isolated from healthy human donors, pre-activated with IL-2/IL-12, incubated with CM from CON or NetG1 KO CAFs for 16 hours, and their activation status was determined by flow cytometry, using IFNγ and CD69 as markers. Expression of markers was normalized to the positive control (IL-2 alone = 1.0). * comparison between the CON and KO at each % of CM. **D-E.** RFP^+^ CON or NGL-1 KO PDACc (2×10^4^) were co-cultured in 3D with GFP^+^ CON or NetG1 KO CAFs (2×10^4^) and with active **(D)** or resting **(E)** NK-92 cells (8×10^4^) for 48 hours and PDAC survival was quantified. Groups were normalized to CON PDACc/CON CAF with active NK cells. Dotted line in the resting graph **(RIGHT)** denotes PDACc survival with CON PDACc/CON CAF with active NK cells. * compared to CON PDACc/CON CAF with active NK cells. One-Way ANOVA, Dunnett’s multiple comparison test **(A, B, D)** or T-test **(C).** *p<0.05, **p<0.01, ***p<0.001; ****p<0.0001. **F.** NK cell killing assay in 2D, where RFP^+^ CON or NGL-1 KO PDACc (2×10^4^) were co-cultured with CON or NetG1 KO CAFs (2×10^4^) and 8×10^4^ active NK-92 cells (IL-2 preactivated) for 6 hours in the presence of isotype control IgG or IL-15 neutralizing antibody. Graphs depict PDACc survival, relative to the CON PDACc/CON CAF condition treated with IgG (dotted green line). * compared to IgG treated CON PDACc/CON CAF with NK cells. **G.** Same assay as in (**F**), but the co-culture is performed in 3D for 48 hours. * compared to IgG treated CON PDACc/CON CAF with NK cells.

To test the influence of CAF NetG1 in NK cells activation and function, we first assayed the activation of an NK cell line, NK-92, in our *in vitro* 3D culturing system. After IL-2 treatment alone, 68.5% and 72.3% of the NK cells became positive for IFN-γ and Granzyme B (i.e. activated), respectively (**Fig 3B**). Conversely, direct co-culture of activated NK cells with CON CAFs or with CM from CON CAFs abolished NK cell activation marker expression, illustrating the potent immunosuppressive potential of CON CAFs. NetG1 KO CAFs, however, were less immunosuppressive, as 12-23% of NK cells maintained IFN-γ or Granzyme B expression. This result is consistent with the decreased levels of immunosuppressive cytokines generated by the NetG1 KO CAFs (**Fig 3A**). A similar experiment was performed using primary NK cells isolated from healthy human donor blood, with IFN-γ and CD69 as markers of activation. As with NK-92 cells, we observed that while CM from CON CAFs inactivates primary NK cells in a concentration dependent manner (i.e. 10% CM prevented NK activation by ∼50%), NetG1 KO CAF CM was significantly less immunosuppressive at the same dilutions (**Fig 3C**). We next tested if the activation status of NK-92 cells correlated with their functional ability to kill PDAC cells. To this end, we performed an experiment co-culturing resting and active NK cells with CON or NGL-1 KO RFP^+^ PDACc, using complete media (to avoid stress due to nutrient deprivation) in 3D. Accordingly, active NK cells were twice as effective in killing both CON and NGL-1 KO PDACc than resting NK cells (**S10A-B**), suggesting that the ability of CAFs to control anti-tumor NK function is independent of PDACc NGL-1 status. To probe how CAFs could directly affect the ability of NK cells to kill PDACc, we co-cultured CON or NetG1 KO CAFs with CON or NGL-1 KO RFP^+^ PDACc in the presence of resting or active NK cells, again using complete media in 3D. In agreement with the observed immunosuppressive profile of CON CAFs (**Fig 3A-B**), active NK-92 cells were prevented from killing PDACc in the presence of CON CAFs, and this protection was decreased by 38-39% when cultured with NetG1 KO CAFs (**Fig 3D, S10A BOTTOM**). These results were effectively replicated when using CAF2 (**S8D**). Intriguingly, even when resting NK cells were cultured in the presence of NetG1 KO CAFs, they were 45-65% more effective at eliminating PDACc compared to co-culture with CON CAFs, suggesting that NetG1 KO CAFs could partially activate NK-92 cells (**Fig 3E, S10A BOTTOM**).

The NK-92 killing assay was replicated in 2D culturing conditions. The KO of NetG1 in CAFs resulted in a significant decrease in CON PDACc or CON PANC-1 survival against NK cell induced death compared to CON CAF conditions (84 and 66% less, respectively) (**S10C-D**). Also, the partial activation of NK cells by NetG1 KO CAFs was reproduced (**S10E**). Interestingly, in 2D conditions, NGL-1 KO cells (KO1 and KO2) were more sensitive to NK cell killing even when co-cultured with CON CAFs (88% and 78% less survival compared to CON PDACc, respectively), suggesting that NetG1/NGL-1 heterotypic cell-cell contacts were important for protection against NK cell induced death in the absence of ECM (**S10C**). Importantly, neither resting nor active NK cells affected CON or NetG1 KO CAFs survival (**S10F**). To question whether tumor to stromal cell ratios are important for the observed effect, we co-cultured various ratios of PDACc to CAFs (5:1, 3:1, 1:3) and again performed the NK cell killing assay. As the number of CAFs increased, PDACc survival increased as well, indicating a direct effect of CAF numbers on NK cell killing function (**S10G**).

Overall, NetG1 expression in CAFs creates an immunosuppressive microenvironment that inactivates NK cells and protects PDAC cells from NK cell mediated death. Loss of NetG1 expression in CAFs partially reverts the immunosuppressive capacity of CAFs, allowing NK cells to maintain their activity and eliminate PDAC cells. Moreover, the microenvironment generated by CAFs (i.e. ECM) also plays an important role in the support of PDAC survival, highlighting the differential effects of 2D culture versus 3D culture.

While deletion of NetG1 in CAFs led to a decrease in the expression of immunosuppressive cytokines, levels of IL-15, a key activator of NK cells, remained significantly higher than in TAs (**Fig 3A**). Thus, we hypothesized that the downregulation of immunosuppressive cytokines, coupled with the maintenance of IL-15 expression, was partially responsible for the observed anti-tumor phenotype of the NetG1 KO CAFs. Thus, we repeated the NK cell killing assay in 2D and 3D, co-culturing CON or NetG1 KO CAFs with CON or NGL-1 KO PDACc, in the presence of a neutralizing IL-15 antibody or IgG isotype control. Indeed, neutralization of IL-15 resulted in a ∼35% and ∼25% (2D and 3D, respectively) increase in PDACc survival compared to IgG treated conditions, independent of PDAC NGL-1 status, suggesting that CAF expressed IL-15 was partially responsible for the anti-tumor microenvironment created by NetG1 KO CAFs (**Fig 3F-G**). Thus, collectively, the *in vitro* NK assays and patient data suggest CAFs lacking NetG1 improve NK cell function, which is better for overall survival in PDAC patients, in a mechanism partially dependent on IL-15 production from CAFs, accompanied by limiting immunosuppressive cytokine secretion.

### p38 and AKT pathways are downregulated in NetG1 KO CAFs and mediate pro-tumor CAF functions

Having identified key functions regulated by NetG1, we sought to uncover the downstream mediators of NetG1 that were responsible for pro-tumor functions of CAFs. We identified a downregulation of p-AKT and p-p38 in NetG1 KO CAFs (**Fig 4A**), also confirmed in CAF2 (**S7F**), and postulated that these two pathways were largely mediating the effects of NetG1. Therefore, we employed the use of pharmacological inhibitors of p38 (p38i) and AKT (AKTi) on CAFs in our 3D system and assessed PDAC survival against metabolic stress and NK induced death, as well as Glu/Gln and cytokine production. Interestingly, we observed a greater decrease on PDACc survival under serum/Gln free conditions when CAFs were treated with AKTi (∼70%) vs. p38i (∼30%) compared to DMSO treated CAFs (**Fig 4B**). Accordingly, AKTi reduced both Glu/Gln production in CAFs (∼30%), while p38i only inhibited Gln production (**Fig 4C**). Together, these findings suggest that AKT regulated the metabolic parameters of CAFs to a greater degree than p38. On the other hand, p38i CAFs displayed a significant decrease in the secretion of more cytokines than AKTi, with a reduction in GM-CSF, IL-6, and TGF-β levels, and a maintenance of IL-15 (akin to NetG1 deficient CAFs), compared to DMSO treated CAFs (**Fig 4D**). In contrast, AKTi CAFs had increased GM-CSF production compared to DMSO treated CAFs, with decreases in IL-8, TGF-β, and IL-15 production. Based on these profiles we hypothesized that p38i CAFs would be less immunosuppressive than AKTi CAFs. To test this, we performed the NK killing assay and saw that indeed p38i CAFs lost their immunosuppressive capacity compared to DMSO CAFs, while AKTi CAFs did not (**Fig 4E**), thus confirming our hypothesis.

**Figure 4.**
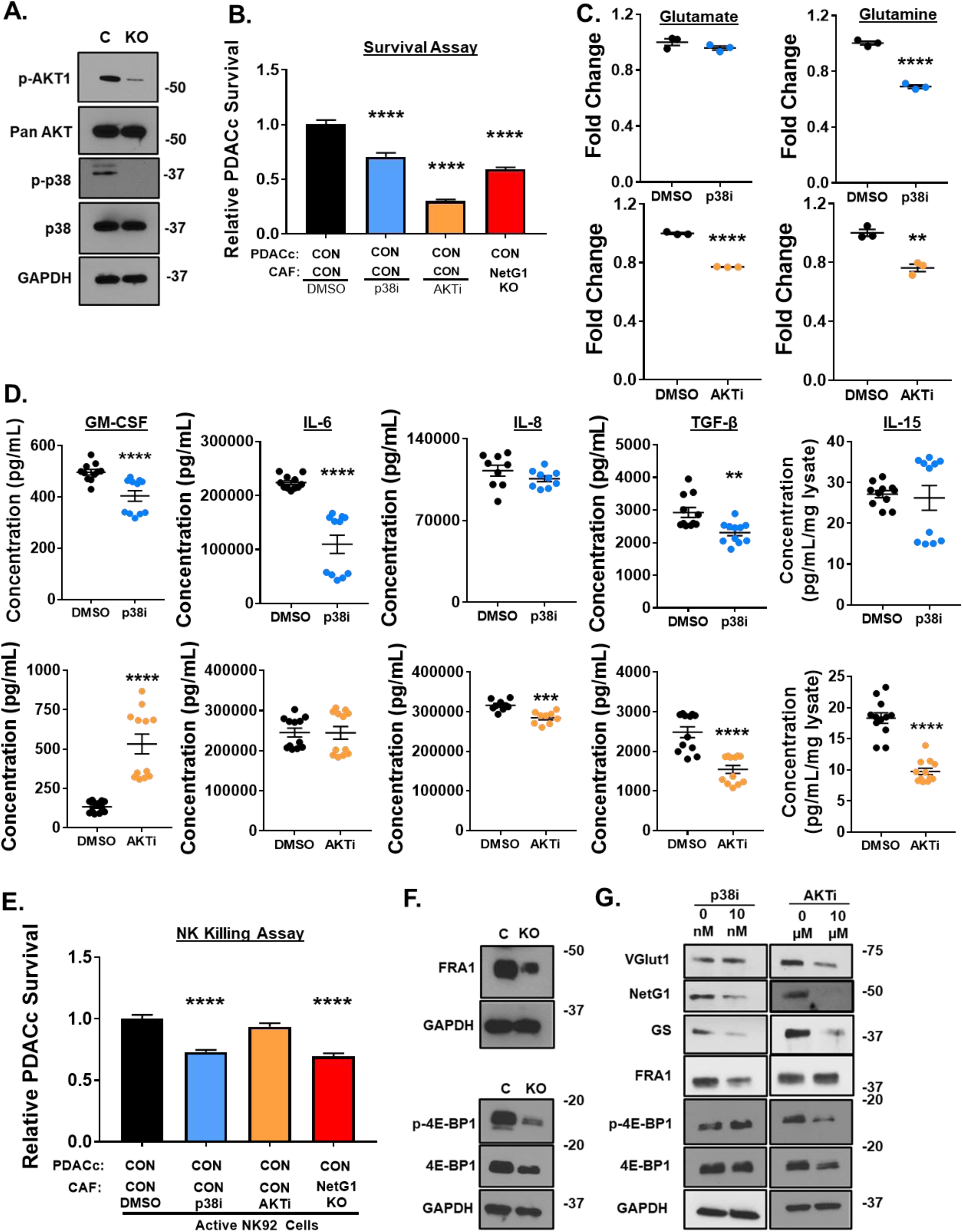
NetG1 functions are mediated through AKT and p38 pathways. **A.** Representative western blots demonstrating downregulation of p-AKT and p-p38 in NetG1 KO CAFs compared to CON. GAPDH was used as a loading control. **B.** RFP^+^ PDACc (2×10^4^) were co-cultured with CON CAFs pre-treated for 24 hours with DMSO, 10 nM p38 inhibitor (p38i), or 10 µM AKT inhibitor (AKTi) and PDACc survival was assessed 96 hours later. * compared to DMSO. **C.** Graphs depicting relative Glu and Gln levels measured from CAFs treated with p38i or AKTi in serum/Gln free media for 48 hours. DMSO was used as a treatment control. * Compared to DMSO. **D.** ELISAs for GM-CSF, IL-6, IL-8, TGF-β, and IL-15 were performed on CM or lysates of CAFs treated for 48 hours with DMSO or p38i (**TOP panel)** and DMSO or AKTi (**BOTTOM panel**). * compared to DMSO. Student’s T-test: *p<0.05, **p<0.01, ***p<0.001; ****p<0.0001. **E.** Quantification of PDACc survival after co-culture with CON CAFs pre-treated for 24 hours with DMSO, p38i, or AKTi in the presence of active NK cells (8*10^4^). **F.** Representative western blots demonstrating a reduction in FRA-1, p-4E-BP1, and 4E-BP1 protein expression in NetG1 KO CAFs compared with CON CAFs. GAPDH was used as a loading control. **G.** Representative western blots of VGlut1, NetG1, GS, FRA1, p-4E-BP1, and 4E-BP1 from the lysates of DMSO or p38i CAFs (**LEFT**) and DMSO or AKTi CAFs 48 hour treated CAFs (**RIGHT**).

To further dissect the p38 and AKT signaling pathways in CON and NetG1 KO CAFs, we tested the expression of numerous known downstream factors of these two pathways. We found substantial modulations of both pathways in NetG1 KO CAFs compared to CON CAFs, with increased levels of FosB and cFos and reduced levels of c-Jun, JunB and FRA-1 downstream to p38, and reduced protein levels of p-FOXO1, p-FOXO3, p-GSK3β, and p-4E-BP1 downstream to AKT (**S8E-F, Fig 4F**). To complement our loss of function approaches, we decided to further explore the modulation of the two most downregulated proteins observed in each pathway, FRA-1 and 4E-BP1, due to their previous association with inflammation or fibrosis [54, 55]. The downregulation of FRA1 and 4E-BP1 was confirmed in CAF2 NetG1 KDs as well (**S7F**). Next, we measured the expression of VGlut1, NetG1, GS, as well as FRA1 and 4E-BP1 in p38i and AKTi CAFs to observe how the developing signaling circuit changed in response to the kinase inhibitors (**Fig 4G**). In accordance with Glu/Glu production, p38i CAFs had reduced protein levels of NetG1 and GS, but not VGlut1, while AKTi CAFs exhibited decreases in all three proteins. In addition, FRA1 and 4EBP-1 were only downregulated in response to the corresponding pathway inhibitor (p38 or AKT, respectively). Overall, the data pointed towards a greater role for p38 in the immunosuppressive functions of CAFs, while AKT regulated metabolic parameters to a greater extent.

### FRA1 and 4E-BP1 partially regulate metabolic and immunosuppressive characteristics of CAFs

Because there was a large downregulation of FRA1 and 4E-BP1 in NetG1 KO CAFs, we decided to explore the functional roles of these proteins by knocking them down in CAFs, and observe how they fit into the NetG1 signaling circuit. First, we obtained one effective KD of FRA1 and 2 effective KDs for 4E-BP1, confirmed by western blotting (**Fig 5A**). Intriguingly, the effects of FRA1 and 4E-BP1 KDs in CAFs on the signaling circuit were quite different. FRA1 KD in CAFs resulted in a loss in expression of VGlut1, NetG1, and GS, while 4E-BP1 KD CAFs up-regulated these three proteins. In terms of the upstream kinases, p-AKT remained unchanged in both KDs, while p-p38 increased in FRA1 KD CAFs and remained largely unchanged in 4E-BP1 KD CAFs. Strikingly, while loss of 4E-BP1 in CAFs had no effect on FRA1 expression levels, knockdown of FRA1 resulted in a downregulation of p-4EBP1 in CAFs. These results illustrated a crosstalk between the p38/FRA1 and AKT/4E-BP1 pathways, as well as a potential compensatory feedback loop that upregulated factors upstream in the circuit. Based on these observations and the findings with p38i and AKTi CAFs, we predicted that FRA1 KD CAFs would functionally phenocopy p38i CAFs and 4E-BP1 KD CAFs would mirror the AKTi CAFs. However, both FRA1 and 4E-BP1 KD CAFs did not support PDACc and PANC-1 survival under metabolic stress compared to CON CAFs (**Fig 5B**). Consequently, Gln levels were lower in the CM of both FRA1 and 4EBP1 KD CAFs compared to CON CAFs, and Glu was lower in 4E-BP1 KD CAFs, which could account for the decreased survival of the PDAC cell lines (**Fig 5C**). On the other hand, when measuring cytokine secretion, FRA1 KD CAFs displayed markedly reduced cytokine levels compared to CON CAFs, and 4E-BP1 KDs had increased cytokine levels, which were in line with the results from p38i and AKTi CAFs (**Fig 5D**). Therefore, we hypothesized that the FRA1 KD CAFs would allow NK cell mediated killing while 4E-BP1 KD CAFs would retain their immunosuppressive capacity. As expected, in the NK killing assay, PDACc and PANC-1 cells co-cultured with FRA1 KD CAFs had significantly reduced survival compared to CON CAFs, while more PDACc and PANC-1 cells survived when co-cultured with 4E-BP1 KDs, similarly CON CAFs, when compared to FRA1 KD CAFs (**Fig 5E**). Collectively, these results suggest that FRA1 and 4E-BP1 contribute to the metabolic support of PDAC cells by CAFs, while FRA1 mediates their immunosuppressive effects. Additionally, FRA1 can modulate the 4E-BP1 side of the signaling circuit, illustrating a crosstalk between the downstream parts of the p38 and AKT pathways.

**Figure 5.**
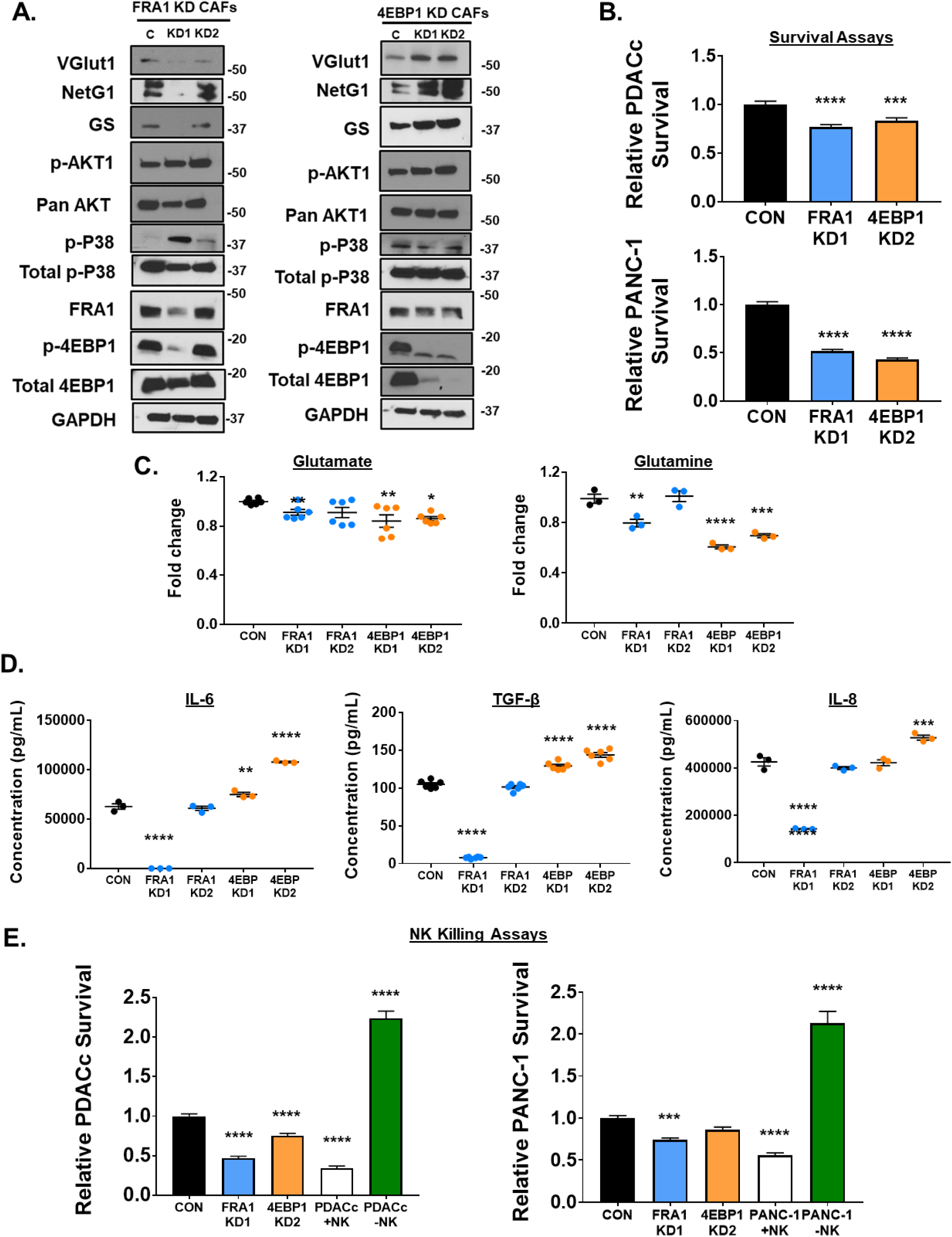
Downstream mediators of p38 and AKT signaling, FRA1 and 4E-BP1, regulate metabolic and immunosuppressive properties of CAFs. A. Representative blots of VGlut1, NetG1, GS, p-AKT, AKT, p-p38, p38, FRA1, p-4E-BP1, and 4E-BP1 in (**LEFT**) CON (C) or FRA1 KD1/KD2 (KD) CAFs and (**RIGHT**) in CON or 4E-BP1 KD CAFs. GAPDH was used as a loading control. **B.** Graphs depicting PDAC survival after assays were performed with PDACc cells (**TOP**) or PANC-1 cells (**BOTTOM**) co-cultured in 3D, under serum/Gln free conditions, with the most effective KD CAFs (FRA1 KD1 and 4E-BP1 KD2) from (**A**), *compared with co-culture with CON CAFs. **C.** Relative Glu and Gln levels detected in the CM of CON, FRA1 KD, or 4E-BP1 KD CAFs after culture for 48 hours in serum/Gln free media in 3D. **D.** Measurement of indicated cytokine production in the CM of CON, FRA1 or 4E-BP1 KD CAFs after culture for 48 hours in serum/Gln free media in 3D, as determined by ELISA. **E.** Quantification of PDAC cell survival after co-culture with CON, FRA1 KD1, or 4E-BP1 KD2 KD CAFs in the presence of active NK cells (8*10^4^). PDAC cells cultured alone with or without active NK cells served as controls. **LEFT:** PDACc; **RIGHT:** PANC-1. *Compared to CON. One-Way ANOVA, Dunnett’s multiple comparison test. *p<0.05, **p<0.01, ***p<0.001; ****p<0.0001.

### VGlut1 and GS inhibition in CAFs modulates metabolic and immunosuppressive functions in CAFs, in a manner similar to NetG1

Because of the consistent joint modulation of VGlut1, GS, and NetG1 across nearly all of the genetically and pharmacologically inhibited CAFs, we evaluated if the absence of VGlut1 or GS in CAFs could phenocopy the effects of NetG1 KO in CAFs. First, we produced knockdowns for VGlut1 (VGlut1 KD) using CRISPRi technology (**Fig 6A**). Interestingly, VGlut1 KD CAFs lost the expression of NetG1 and GS (**Fig 6B**), suggesting a co-regulatory mechanism for these proteins. Functionally, VGlut1 KD CAFs had a similar effect on PDACc survival as NetG1 KD CAFs, with PDACc survival reduced by ∼50% compared to CON CAFs (**Fig 6C**). Moreover, VGlut1 KD CAFs produced similar amounts of Glu and Gln, as was observed for NetG1 KD CAFs, both producing significantly less Glu/Gln than CON CAFs (**Fig 6D, Sup Table 10**). In terms of their cytokine profile, VGlut1 KD CAFs produced significantly less GM-CSF, IL-6, and TGF-β than CON CAFs, while maintaining IL-15 levels (**Fig 6E**), indicative of less immunosuppressive capacity. Indeed, in the NK killing assay, PDACc cells had ∼30-40% decreased survival when co-cultured with VGlut1 KD CAFs compared to CON CAFs (**Fig 6F**). Related to downstream signaling, VGlut1 KD CAFs displayed reduced levels of p-AKT, p-p38, FRA1, and p-4E-BP1, phenocopying NetG1 KO CAFs (**Fig 6G**). Lastly, we stained PDAC patient tissue for VGlut1 expression and detected high fibroblastic expression in 4/14 cases, and low expression in 10/14 cases (**Fig 6H**). In sum, VGlut1 is expressed in human CAFs, and regulates expression of proteins and functions in CAFs involved in supporting PDAC cell survival, phenocopying NetG1 functions.

**Figure 6.**
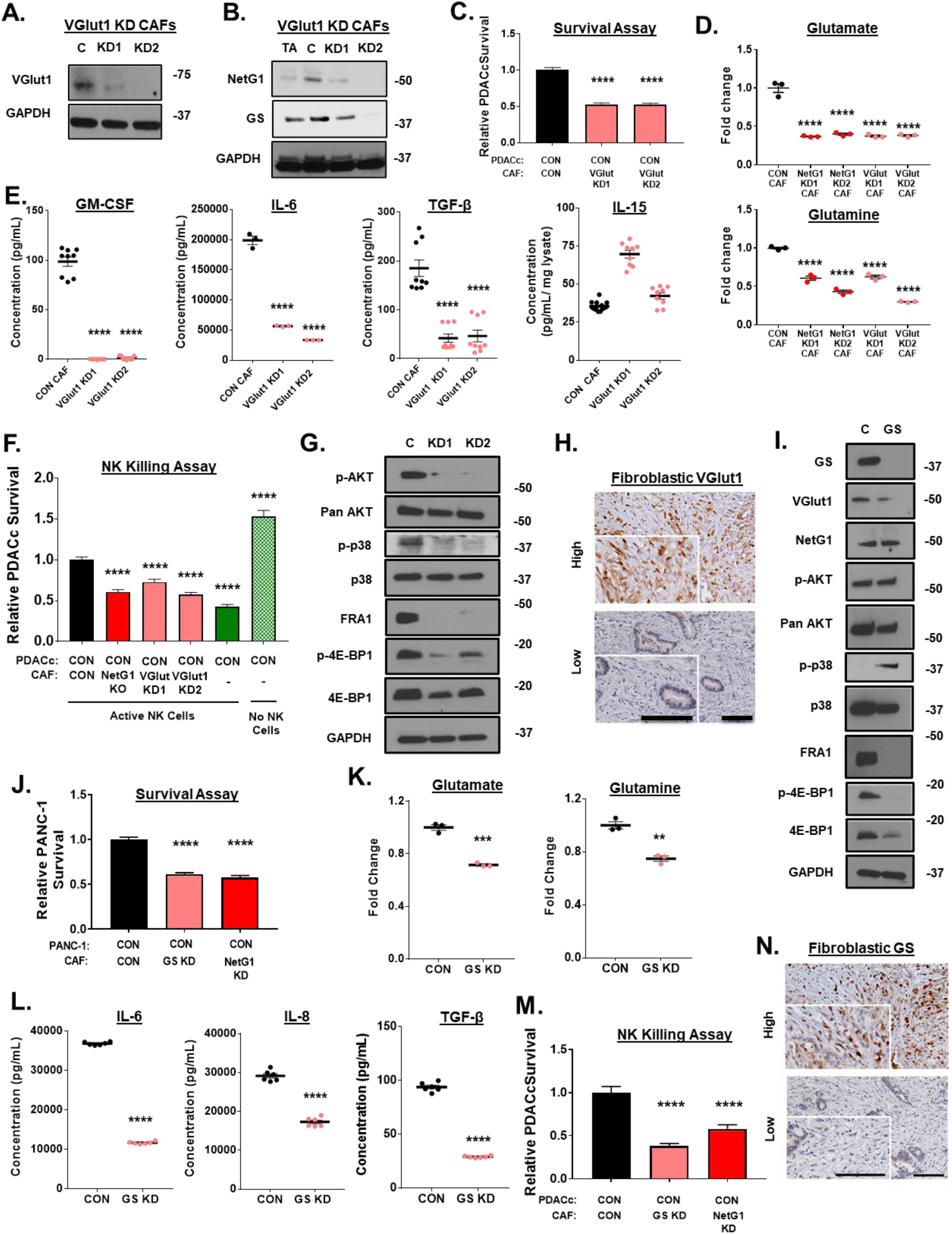
VGlut1 and GS regulate pro-tumor CAFs functions, in a manner similar to NetG1. **A.** Representative western blots of VGlut1 in CAFs, demonstrating effective knockdown (KD) by CRISPRi. **B.** Representative western blot of NetG1 and GS in TA, CON, or VGlut1 KD CAFs. GAPDH was used as a loading control. **C.** RFP^+^ CON PDACc cells (2×10^4^) were cultured in 3D with CON or VGlut1 KD CAFs (2×10^4^) for 96 hours in SF/Gln Free media and PDACc cell survival was assessed. **D.** Relative Glu and Gln levels in the CM of NetG1 and VGlut1 KD CAFs compared to CON CAF. Note that both NetG1 KD and VGlut1 KD CAFs have a reduction in Glu/Gln levels. * Compared to CON CAF. **E.** Measurement of indicated cytokine production in the CM of CON or VGlut1 KD CAFs after culture for 48 hours in serum/Gln free media in 3D, as determined by ELISA. * Compared to CON CAF. **F.** Quantification of PDACc survival after co-culture with CON or VGlut1 KD CAFs in the presence of active NK cells (8*10^4^). One-Way ANOVA, Dunnett’s multiple comparison test. *p<0.05, **p<0.01, ***p<0.001; ****p<0.0001**. G.** Representative western blots of p-AKT1, AKT, p-p38, p38, FRA1, p-4E-BP1, and 4E-BP1 comparing protein expression in CON and VGlut1 KD CAFs. GAPDH was used as a loading control. **H.** Representative images of the immunohistochemical staining of VGlut1 in patient tissue, segregated into high or low fibroblastic VGlut1 expression. Inserts correspond to magnified regions, in which is stromal cells are more evident. Scale bars represent 100 μm. **I.** Representative blots of GS, VGlut1, NetG1, p-AKT, AKT, p-p38, p38, FRA1, p-4E-BP1, and 4E-BP1 in CON (C) or GS KD (GS) CAFs. GAPDH was used as a loading control. **J.** RFP^+^ CON or NGL-1 KO PDACc (2×10^4^) were cultured in 3D with CON or GS KD CAFs (2×10^4^) for 96 hours in SF/Gln Free media and PDACc cell survival was assessed. **K.** Relative Glu and Gln levels in the CM of GS KD CAFs compared to CON CAF. **L.** Measurement of indicated cytokine production in the CM of CON or GS KD CAFs after culture for 48 hours in serum/Gln free media in 3D, as determined by ELISA. **M**. Quantification of PDACc survival after co-culture with CON, GS KD, or NetG1 KD CAFs in the presence of active NK cells (8*10^4^). *Compared to CON. One-Way ANOVA, Dunnett’s multiple comparison test. *p<0.05, **p<0.01, ***p<0.001; ****p<0.0001**. N.** Representative images of the immunohistochemical staining of GS in patient tissue, segregated into high or low fibroblastic GS expression. Inserts correspond to magnified regions, in which is stromal cells are more evident. Scale bars represent 100 μm.

To address the contribution of GS to the support provided by CAFs for PDACc survival, we genetically and pharmacologically inhibited GS in CAFs, using CRISPRi and increasing concentrations of methionine sulfoximine (MSO), respectively. GS was knocked down with high efficiency (**Fig 6I**), and MSO effectively inhibited GS expression at concentrations greater than 0.1 mM (**S11A**). GS KD CAFs exhibited decreased levels of VGlut1, FRA1, and p-4E-BP1, while maintaining expression of NetG1, and p-AKT, and increasing p-p38 (**Fig 6I**). Transient inhibition with MSO resulted in decreases of VGlut1 and NetG1, as well as downstream mediators p-p38 FRA1, and p-4E-BP1, with p-AKT increasing in expression (**S11B**). We attribute the differences in signaling molecules to the nature of the treatment (transient vs. permanent), which may have resulted in compensatory up-regulation of p-AKT, for example. Nevertheless, blockade of GS led to an overall downregulation of many proteins in the NetG1 signaling circuit. Therefore, due to the similar profile, we predicted that loss of GS would functionally mimic the loss of NetG1 and VGlut1 in CAFs and reduce PDAC survival in our metabolic stress assay and when challenged with NK cells. Accordingly, PDAC cells that were co-cultured with either GS KD or MSO treated CAFs under serum/Gln free conditions survived significantly less than when cultured with CON CAFs, similarly to co-culture with NetG1 KO CAFs (**Fig 6J, S11C**). This was likely due to the decreased levels of Glu/Gln detected in the CM of GS KD CAFs (**Fig 6K**), comparable to other inhibited CAFs in the NetG1 signaling hub that do not support PDAC survival. The immunosuppressive activity was determined in both GS KD and MSO treated CAFs by measuring cytokine secretion and protection of PDAC cells from NK cell induced death. Both GS KD and MSO treated CAFs showed a decrease in most cytokines tested (**Fig 6L, S11D**), and this resulted in a corresponding decrease in PDAC cell survival when co-cultured with GS KD or MSO treated CAFs, similarly to co-culture with NetG1 KO CAFs (**Fig 6M, S11E**). Finally, we performed IHC on patient PDAC tissue to detect fibroblastic expression of GS and found that 7/15 patients expressed high levels of GS in fibroblasts, and 8/15 had low expression (**Fig 6N**). Collectively, these data demonstrate how NetG1, VGlut1, and GS may be all linked in an emerging signaling circuit in CAFs, driving the metabolic and immunosuppressive functions of CAFs.

### Therapeutic inhibition of NetG1 ablates the pro-tumor functions of CAFs, stunting tumorigenesis *in vivo*

Having established the roles that NetG1 played in driving pro-tumor CAF functions, we next explored the therapeutic potential of targeting NetG1 *in vitro* and *in vivo*. We identified a neutralizing monoclonal antibody against NetG1 (mAb), which was determined by testing its ability to modulate NetG1 mediated signaling, Glu/Gln/cytokine production, and functions in CAFs *in vitro*. We measured the changes in expression of the proteins in the signaling hub, by dose and time responses. Anti-NetG1 mAb began to decrease expression of the majority of the signaling circuit at a concentration of 1.25 µg/mL, with maximal efficacy at 10 µg/mL (**Fig 7A**). We proceeded with the minimal dose that inhibited all proteins (2.5 µg/mL) to explore the changes in protein expression over time. Interestingly, most proteins lost substantial expression by 24 hours of treatment, with only GS having a transient increase in expression between 30-60 minutes of treatment (**Fig 7B**), suggesting a post-translational stabilization of this protein. p-AKT/p-4E-EP1 remained downregulated after 60 minutes of mAb treatment, while p-p38 levels rapidly decreased starting at 5 minutes, and rebounded by 120 minutes of treatment. These results largely confirm the putative signaling circuit uncovered in CAFs that we observed through the systematic inhibition of each individual component. Functionally, anti-NetG1 mAb treated CAFs co-cultured with PDACc or PANC-1 cells resulted in a ∼25-40% reduction in PDAC cell survival under metabolic stress compared to IgG treated CAFs, similar to co-culture with NetG1 KO CAFs (**Fig 7C**), owed likely to the reduced Glu/Gln production in mAb treated CAFs (**Fig 7D**). In addition, anti-NetG1 mAb treated CAFs produced less pro-tumor cytokines compared to IgG treated CAFs, showing a dose dependent decrease in IL-6, IL-8, and TGF-β, suggesting less immunosuppressive capacity (**Fig 7D**). Thus, inhibition of NetG1 by mAb was extremely effective at modulating all pro-tumor signaling and functions of CAFs, serving as the impetus to move the treatment *in vivo*.

**Figure 7.**
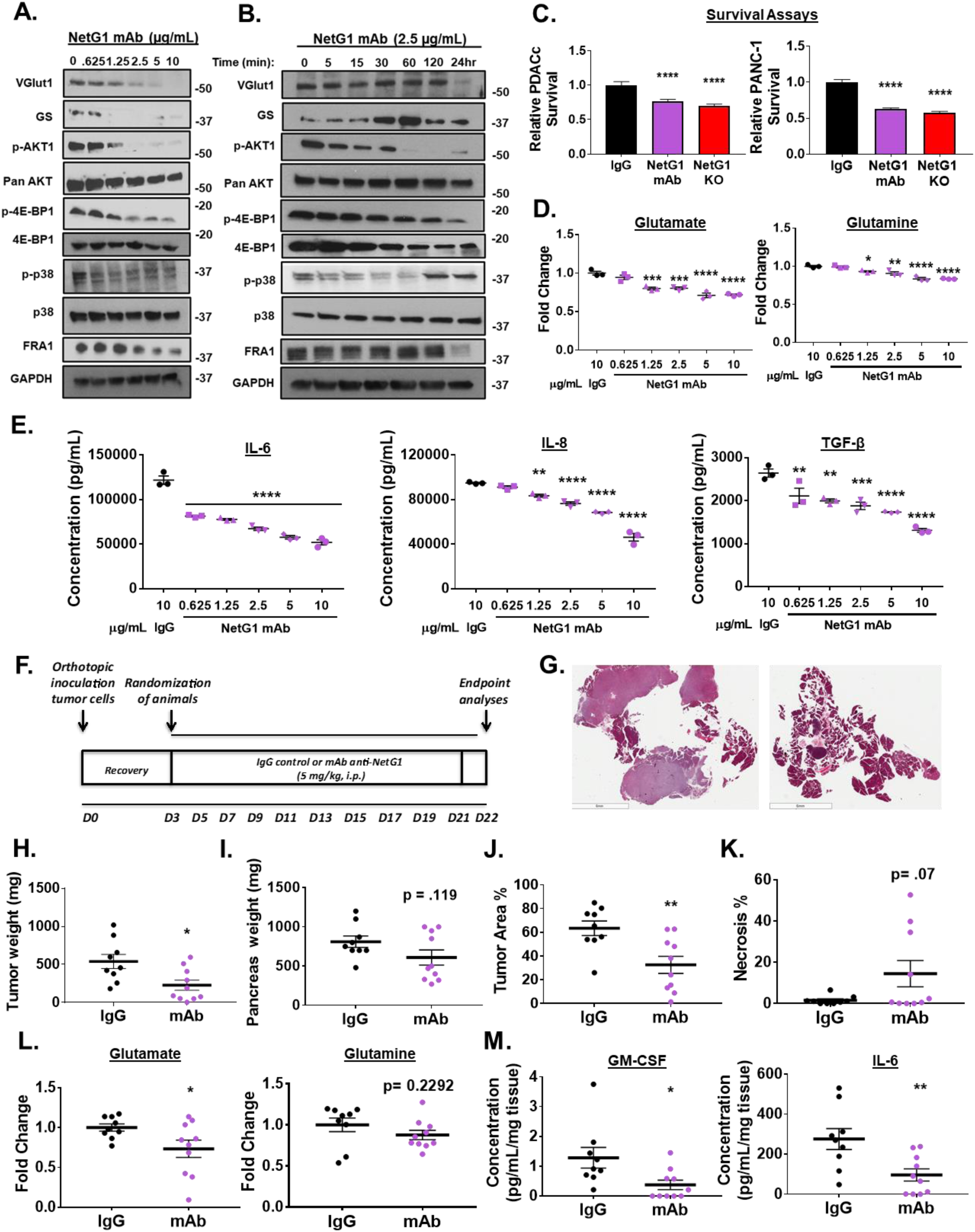
A neutralizing monoclonal antibody targeting NetG1 inhibits pro-tumor properties of CAFs and decreases tumor burden *in vivo*. ***A+B.*** Representative western blots demonstrating a dose (**A**) and time (**B**) dependent decrease in proteins associated with pro-tumorigenic pathways in CAFs after anti-NetG1 monoclonal antibody (mAb) treatment in serum and glutamine free media, with 48 hour treatment for CAFs in (**A**) and indicated timepoints for CAFs in (**B**). **C.** Quantification of PDAC cell survival (PDACc or PANC-1) under serum and glutamine starvation after 48 hour co-culture with CON (10 µg/mL IgG or mAb treated) or NetG1 KO CAFs, demonstrating the efficacy of NetG1 mAb in a key functional assay. **D.** Graphs depict amounts of secreted Glu and Gln detected in the conditioned media of CAFs treated with IgG or increasing doses of mAb for 48 hours, as determined by ELISA. **E.** Graphs depict amounts of secreted pro-tumor cytokines detected in the conditioned media of CAFs treated with IgG or increasing doses of mAb for 48 hours, as determined by ELISA. Note the dose dependent decrease in these key secreted factors in (**D)** and (**E**), important for PDAC tumorigenesis. * Compared to IgG. One-way ANOVA: * p <.05, ** p<.01, *** p<.001, **** p<.0001. **F.** Treatment Strategy: C57/BL6 mice were injected orthotopically with 10^6^ murine pancreatic cancer cells and mice were allowed to heal for 3 days. Starting on day 4, mice were separated into two groups, IgG (n=9) and mAb (n=10), and received either 5mg/kg IgG or mAb against NetG1. Mice were treated 3 times per week until the completion of the model, at day 22. Pancreata were isolated from the mice and H+E sections were developed from a cut of the entire pancreas. **G.** Representative images of the H+E sections of the pancreas from each treatment group. Scale bar: 6 mm. **H.** Graph depicting relative tumor weights. **I.** Graphs depicting the weights of the pancreata isolated from the mice. **J.** Quantification of % area of the pancreas that was classified as tumor. **K.** Quantification of the % area of the pancreas that was classified as necrotic. **L+M**. A small piece of tumor tissue (30-60 mg) was excised from the pancreas of each mouse and was cultured overnight in DMEM lacking serum, glutamate (Glu) and glutamine (Gln). The resultant media was collected and the amount of Glu and Gln (**L**) or cytokines (**M**) were measured by ELISA and normalized to the weight of tumor tissue cultured. Student’s T-Test was used to determine if samples were significantly different from IgG. * p <.05, ** p<.01, *** p<.001, **** p<.0001. n=9 for IgG, n=10 for mAb.

Next, we decided to test the efficacy of mAb *in vivo,* relying upon our previously performed syngeneic orthotopic injection model of PDAC in mice. Mice were injected with KPC3 PDAC cells and allowed to recover for 3 days, and were then split randomly into 2 groups, one receiving 5 mg/kg isotype control IgG and the other receiving 5 mg/kg anti-NetG1 mAb, three times a week for the duration of the experiment (22 days) (**Fig 7F**). Excitingly, anti-NetG1 mAb treatment was effective at limiting tumorigenesis, evident in the representative images of the pancreata isolated from IgG or mAb treated animals (**Fig 7G**). Mice treated with anti-NetG1 mAb had less tumor weight, a trend towards less pancreas weight, less % tumor area, and greater necrosis, all indicative of the efficacy of the therapy (**Fig 7H-K**). Moreover, analysis of the tumor culture of isolated pieces of the pancreas from each group demonstrated a significant decrease in Glu, GM-CSF, and IL-6 protein levels, with a trend towards less Gln, supporting our *in vitro* findings (**Fig 7L-M**). Overall, in an extremely aggressive orthotopic model, anti-NetG1 mAb treatment was successful at limiting PDAC tumorigenesis, as well as at reducing key CAF generated metabolites and cytokines, highlighting NetG1 as an important druggable target in PDAC.

## Discussion

The present study has uncovered a previously unknown target for the potential treatment of pancreatic cancer: the neural glutamatergic synaptic stabilizing protein NetG1, which has been shown to engage its sole postsynaptic receptor NGL-1 [56]. We demonstrated that NetG1 expressed in CAFs, and NGL-1 in cancer cells, promoted tumorigenesis in multiple murine models *in vivo*, and that NetG1 was a viable therapeutic target. In PDAC patients, fibroblastic expression of NetG1 inversely correlated with overall survival. Functionally, CAFs supported PDAC cell survival by simultaneously providing nutritional support, through Glu/Gln, as well as suppressing NK cell induced PDAC cell death, all in a NetG1 dependent manner. Mechanistically (**S12**), NetG1, VGlut1, and GS were all associated upstream to two major kinase pathways, p38/FRA1 and AKT/4E-BP1, in which the ablation of any of the three proteins resulted in the downregulation of both of these kinase pathways and regulated the pro-tumor functions of CAFs. Overall, our results reveal a complex signaling hub in CAFs, governed by NetG1, and that therapeutic targeting of fibroblastic NetG1 could significantly stunt tumorigenesis.

NetG1 (also known as Laminet-1 and not to be confused with Netrin-1) is a member of the netrin family of proteins, but is unique because it is anchored to the plasma membrane through a glycosylphosphatidylinositol (GPI) linkage, and is not secreted like the majority of netrin family members [34]. NetG1 is a synapse stabilizing protein, located pre-synaptically on neurons, and its only known receptor is NGL-1, discovered on post-synaptic neurons [35, 36]. Dysregulation of NetG1 has been associated with Rett syndrome, schizophrenia, bipolar disorder, and anxiety [57–59]. In cancer, NetG1 displays a low mutation rate [60] and NetG1 mRNA expression, methylation, or mutational status has been linked to colon and uterine cancers [60–62]. Nevertheless, no studies have ascribed a functional role to NetG1 in the context of cancer.

PDAC is the most common form of pancreatic cancer, and has a dismal prognosis [2]. CAFs are a major cell type present in the PDAC microenvironment, whose role is incompletely understood. CAFs regulate at least three critical hallmarks of PDAC: 1.) the deposition of desmoplastic ECM; 2.) overcoming nutrient deprivation; and 3.) vast immunosuppression of anti-tumor immune cells [3, 25]. This study aimed to characterize the functions of NetG1 in PDAC, with an emphasis on the contribution of the desmoplastic stroma. Our results confirm that all three hallmarks are interconnected, and that NetG1 expression in CAFs is a central driver of all of them in the PDAC microenvironment.

The PDAC microenvironment is nutrient deprived, due to the collapse of blood vessels enabled by desmoplasia, and this drives PDAC cells to use an alternative nutritional supply [16]. Recent studies have demonstrated that CAFs supply PDAC cells with important metabolites, through exosomes [50] and amino acid secretion [49]. Moreover, the ECM has been shown to be utilized as fuel for cancer cells [48], which potentially positions the scaffolding produced by CAFs in the PDAC microenvironment as a vital source of energy for PDAC cells. Here, we found that CAFs lacking NetG1 were less supportive of PDAC cell survival, in direct co-culture, with their CM, or ECM (**Fig 2**). These findings prompted us to perform a screen comparing amino acid production and secretion and revealed a NetG1 dependent increase in extracellular Glu/Gln in CAFs.

Interestingly, it has long been recognized that patients with pancreatic cancer have elevated serum levels of Glu [63], with a corresponding depletion of serum Gln levels [64]. PDAC cells have a high capacity to utilize Glu, by catabolizing Gln through the enzyme glutaminase (GLS), in a process known as glutaminolysis [21]. While it is known that inhibition of GLS can decrease cancer cell growth *in vitro* and *in vivo* in a number of cancer models [19, 51], a recent study using a potent small molecular inhibitor of GLS displayed little efficacy in preclinical mouse models, in part through up-regulation of compensatory Gln procuring pathways in PDAC cells [65]. Additionally, it has also been demonstrated that the conditions used in cell culture provide an overabundance of nutrients that does not accurately mimic the tumor microenvironment *in vivo*, and culturing cells in a medium with nutrients at levels more representative of tumors *in vivo* resulted in tumor cells with a decreased dependence on Gln, due to increased utilization of cystine [66]. Intriguingly, an elegant recent study suggested that in mouse interstitial fluid of PDAC tumors, Gln levels were surprisingly higher than previously thought [67]. It is tempting to speculate that CAFs may be responsible for the maintenance of Gln levels, by utilizing the high levels of Glu produced and secreted by tumor cells. Thus, our findings that CAFs can assist PDAC cells in overcoming nutritional stress by supplying key metabolites, like Glu/Gln, adds to the ways in which cancer cells in PDAC can acquire Gln from the tumor microenvironment.

Our results demonstrated that ablation of NetG1 in CAFs reduced GS (catalyzing conversion of Glu to Gln), but not GLS protein levels (**Fig 2**). Indeed, GS is up-regulated in CAFs in ovarian cancer compared to their normal counterparts, and inhibition of stromal GS resulted in decreased tumor development [52]. Additionally, we found that deletion of NGL-1 from PDAC cells resulted in a decrease in GLS expression (**S4**), signifying an even greater dependence of these cells on the microenvironment, evident by the lack of rescue during Glu/Gln addition in NGL-1 deficient PDAC cells. An additional interpretation of the data could be that CAF driven metabolism of Glu/Gln is required to first process these amino acids into other metabolites that are then used by PDAC cells. Analysis of TGCA data demonstrated that patients expressing the highest NGL-1 levels have enriched pathway signatures associated with increased glutamate receptor binding genes (**S4**), suggesting a link between NGL-1 and Glu uptake. Further analysis into the expression and activity of receptors that mediate uptake of extracellular amino acids in PDAC, such as Glu/Gln, as well as TCA cycle metabolites produced by CAFs, will be necessary to gain more insight into the general uptake ability of PDAC cells.

We also observed a dramatically reduced expression of VGlut1 upon NetG1 loss in CAFs (**Fig 2**). Interestingly, VGlut1 is present in glutamatergic synapses in neurons, where it loads Glu into pre-synaptic vesicles [53], and VGlut1 expression in neurons was shown to be dependent on NetG1 expression [35]. These studies agree with our data in CAFs, where the KO of NetG1 resulted in downregulation of VGlut1 expression, and provide a potential mechanistic explanation for the decreased levels of Glu in NetG1 KO CAF CM. Interestingly, VGlut1, 2 and 3 have been implicated in pancreatic neuroendocrine tumors (PNET), and Glu signaling in PNET tumor cells drives tumorigenesis [68]. Glu is a key neurotransmitter, but to maintain homeostasis and avoid excitotoxic stress, excess Glu loaded vesicles are then taken up by neighboring astrocytes, which express high levels of GS, is recycled into Gln, and subsequently secreted back to neurons, where the cycle can continue [69]. Our results suggest that NetG1^+^ CAFs may serve as an “astrocyte-like” recycling cell in the PDAC microenvironment, providing a constant supply of Gln to cancer cells, fueling tumorigenesis.

Another hallmark of PDAC tumors is an extremely immunosuppressive microenvironment. Numerous studies have confirmed a major role for CAFs in generating factors, such as secreted cytokines and chemokines [26, 27], that contribute to this environment, as well as physically sequestering anti-tumor immune cells [28]. Furthermore, compared to many other tumor types, PDAC has one of the lowest mutational burdens and display of neoantigens [70, 71], which correlates directly with low response rates to immune checkpoint inhibitors, such as PD-1 [72]. Collectively, these findings suggest that even if cytotoxic T-cells could penetrate the tumor and CAF driven immunosuppression was inhibited, the lack of tumor antigens would render them unable to kill PDAC cells effectively. However, the innate cytotoxic immune cell population— natural killer (NK) cells—can destroy PDAC cells independent of neoantigen presentation, and thus represents an attractive anti-tumor cell type for PDAC therapy [73]. Additionally, NK cells play a natural role in the pancreas, which is to remove activated stromal cells during acute fibrosis in wound healing by a process known as nemosis [74]. Importantly, they are unable to accomplish this in the setting of PDAC, due to the overwhelming numbers of CAFs and immunosuppressive signals, resulting in a situation akin to unresolved chronic wound healing [74, 75]. Thus, reverting the immunosuppressive phenotype of CAFs could allow for NK cell stimulating therapies (e.g. IL-15 supplementation) [76] to more efficiently clear pro-tumoral CAFs as well as PDAC cells. In the present work, we confirmed that CAFs produce the immunosuppressive cytokines TGF-β and GM-CSF, which can expand immunosuppressive myeloid cells [77, 78], as well as chemokines known to attract immunosuppressive immune cells (IL-8 and CCL20) (**Fig 3**). Intriguingly, we showed that CAFs, compared to tumor adjacent fibroblasts (TAs), upregulate IL-15, a potent stimulator of NK cell activity. Functionally, however, IL-15 activity is likely overwhelmed by the greater number of immunosuppressive factors secreted by CAFs, especially TGF-β, as CAFs significantly inhibited NK cell activation and function. Our results support this hypothesis, as CAFs lacking NetG1 failed to protect PDAC cells from NK cell induced death, and that IL-15 was responsible for activation of NK cells. Collectively, the data demonstrate that NetG1 in CAFs shapes the immunosuppressive microenvironment in PDAC, and its ablation reprograms CAFs to an anti-tumor phenotype allowing NK cells to eliminate cancer cells.

In recent years, there has been growing interest in linking microenvironmental metabolites with immune cell function [30], and CAFs in the PDAC microenvironment are important players in this connection [21]. Thus, we postulated that CAF metabolism, driven by NetG1, and by extension, VGlut1 and GS, could also be partially responsible for the immunosuppressive phenotype of CAFs. Importantly, we observed a significant decrease in amounts of immunosuppressive cytokines in the CM of VGlut1 or GS KD CAFs (**Fig 6**), akin to CM from NetG1 KO CAFs. Furthermore, both VGlut1 and GS KD CAFs had a functional effect on NK cells, as PDAC cell survival was significantly decreased in the NK cell elimination assay, probably due to the decreased secretion of immunosuppressive factors. Therefore, our results demonstrate a link between the metabolic circuit, NetG1-VGlut1-GS, and suppression of anti-tumor immunity. Taking lessons from the brain and synaptic biology, it is known that cytokines, such as TNF-α, IFN-γ, IL-6, TGF-β, and IL1-β regulate glutamatergic synaptic activity and subsequently Glu release and uptake by neurons and astrocytes [79–81]. Our data suggests that a reciprocal signaling mechanism may exist in CAFs, whereby disruption of synaptic proteins like NetG1 and VGlut1 result in modulation of cytokine secretion from CAFs, and Glu/Gln cycling may also play a role, as GS inhibition also affected cytokine secretion in CAFs. In sum, our findings illustrate that simply modulating NetG1 expression in CAFs can simultaneously “disarm” two major pro-tumorigenic features of CAFs: metabolic support of PDAC cells and protection from immune cell killing. These data suggest that synaptic biology in CAFs might be a missing link that controls metabolic and immunosuppressive programs in pancreatic cancer. Indeed, PDAC has been found to have a general dysregulation of axonal guidance genes [82], and neo-neurogenesis and perineural invasion have been shown to play roles in promoting tumorigenesis [83]. Thus, PDAC could have a general neural reprograming that is hijacked for the benefit of the tumor.

Mechanistically, little is known about NetG1 mediated signaling. Thus far, studies have shown that NetG1 physically interacts with its receptor, NGL-1, *in trans* on post-synaptic neurons [36] and *in cis* to leukocyte antigen receptor (LAR), a tyrosine protein phosphatase [35], on presynaptic neurons. Additionally, NetG1 has also been shown to modulate expression levels of the potassium channel KATP [84], but no downstream signaling pathways have been associated with any of these findings. Thus, we sought to uncover signaling downstream to NetG1. Our results revealed that NetG1 controlled the activity of the kinases p38 and AKT, as well as the known mediators of those pathways (**Fig 4-5**). In particular, FRA1 and 4E-BP1 were drastically downregulated and, collectively, these pathways could largely account for the pro-tumor functions in CAFs regulated by NetG1. Previously, p38 MAP kinase activity in CAFs has been shown to promote breast and K-Ras driven lung carcinogenesis, by enhancing the secretion of pro-tumor factors (akin to senescence associated secretory proteins) and activating CAFs to produce hyaluronan networks [85, 86]. Additionally, AKT activity in CAFs, mediated through the actin binding protein Girdin, has been shown to enhance lung tumorigenesis [87]. Downstream to p38 is the AP-1 transcription factor complex, which is a dimer that can be comprised of members of the FOS, JUN, ATF, and MAF families, with FRA1 belonging to FOS family [88]. AP-1 has long been associated with cellular responses to stress (activated by MAP kinases such as p38) and inflammatory signals [89]. FRA1 has been linked to activated CAF methylation signatures in PDAC [90], as well as in certain CAF subtypes in breast, as determined by single cell RNAseq [91]. FRA1 is upregulated in a variety of tumors, including pancreatic cancer [88]. Our results demonstrated that FRA1 regulated cytokine production in CAFs and NK cell induced death of PDAC cells, supporting the notion that FRA1 acts as an AP-1 subunit. We also found that 4E-BP1, a critical regulator of cap dependent translation [92], was highly expressed in CAFs and lost upon ablation of NetG1 (**Fig 5**). Previous work has shown that 4E-BP1 plays a major role in fibrogenesis [54], and a protein synthesis inhibitor that blocks mTOR/4E-BP1 in pancreatic CAFs, reverts chemotherapeutic resistance in mice, likely by blocking secretory factors from CAFs [93]. This supports our findings that CAFs lacking 4E-BP1 display a less tumorigenic profile, with the downregulation of Glu/Gln and cytokines. The identification of a neutralizing antibody allowed us to transiently block NetG1, and we were able to further confirm the signaling network downstream to NetG1 (**Fig 7**). This positions NetG1 as an attractive target, as the blockade of a single protein inhibits multiple pathways known to sustain pro-tumor functions in CAFs. This was further confirmed by the anti-tumor effects seen *in vivo*, when genetically targeting NetG1 in CAFs **(Fig 2)** or blocking it with a neutralizing antibody during tumor development **(Fig 7)**. The developing NetG1 signaling circuit is presented in **Supplemental Figure 12.**

A major question in the field is what role do CAFs play in PDAC tumorigenesis? Several studies have demonstrated that CAFs are tumor promoting [9, 10], while others have suggested tumor restrictive roles [12, 13]. One possible explanation for the seemingly paradoxical findings is that not all CAFs are created equally, as it is now appreciated that there is great heterogeneity in the fibroblastic population within the tumor microenvironment [94]. Recent studies have revealed multiple CAF subsets in a variety of tumors [38, 43, 95–97], each with different functions. Our group has identified two distinct CAF subsets in PDAC that are predictive of patient outcome, as patient OS inversely correlated with p-FAK and activated α5β1 integrin expression [38]. Accordingly, NetG1 KO CAFs displayed lower levels of both p-FAK and activated α5β1 integrin. We now propose that NetG1 expression in CAFs stratifies PDAC associated CAFs into two major subtypes: Class 1 (***C1***) anti-tumor (**NetG1 low/negative CAFs, with low GS, VGlut1, p38/FRA1, and AKT/4E-BP1**) and Class 2 (***C2***) pro-tumor (**NetG1 positive CAFs, maintaining high GS, VGlut1, p38/FRA1, and AKT/4E-BP1**). ***C2*** CAFs functionally support PDAC cell survival and shield them from NK cell induced death, while ***C1*** CAFs fail to provide metabolic or immunosuppressive support, independent of myofibroblastic features. These findings help explain how CAFs can be tumor restrictive in some settings, while tumor promoting in others, highlighting the need to fully characterize CAF biology and heterogeneity in order to develop more effective therapeutics. For example, systemic pFAK inhibition in a variant of the KPC mouse model, was shown to revert the desmoplastic phenotype of tumors, including immunosuppression, which improved immunotherapeutic responses [98], in agreement with our *in vivo* targeting of NetG1 (**Fig 7**).

Overall, we have identified a new potential desmoplastic target, NetG1, for the treatment of pancreatic cancer, for which there are no effective therapeutic interventions. We characterized two phenotypes of CAFs, the anti-tumor ***C1*** and the pro-tumor ***C2***, and by targeting NetG1 in CAFs, we can revert ***C2*** CAFs back into tumor restrictive ***C1*** CAFs, ultimately limiting tumorigenesis *in vivo*. Finally, NetG1 was established as a viable therapeutic target in pre-clinical murine models. It is our hope that this study provides strong rationale to consider effects of the stroma on cancer development and progression, and that specific normalization of the stroma is a reliable therapeutic option.

## Methods

### Reproducibility: Key Resources Table

All information (manufacturer/source, RRID, website) regarding key antibodies, mouse models, chemical and biological reagents, software, and instruments has been compiled into a master “Key Resources” Table.

### Cell Culture

The PDAC cells used for the majority of the assays, termed “PDACc” in this study [39], were maintained in media containing 1 part M3 Base F (INCELL, San Antonio, TX) to 4 parts Dulbeco Modified Eagle Medium (DMEM) low glucose (1 g/mL) + sodium pyruvate 110 mg/mL + 1% Penicillin-Streptomycin (10,000 U/mL; Thermo Fisher Scientific, Waltham, MA), supplemented with 5% fetal bovine serum (Atlanta Biologicals, Flowery Branch, GA). PANC-1 cells were maintained in DMEM supplemented with 5% FBS and 1% penicillin-streptomycin (P/S) (10,000 U/mL). Patient matched tumor adjacent fibroblasts and CAFs [38] were maintained in DMEM containing 15% FBS, 1% P/S, and 4 mM glutamine. For individual experiments, cells were cultured as described in the figure legends or methods, either with the maintenance conditions outlined above, or in DMEM without serum (serum free), or DMEM without serum and without glutamine (-). NK-92 cells were maintained in Alpha Minimum Essential medium without ribonucleosides and deoxyribonucleosides supplemented with the following components: 2 mM L-glutamine, 1.5 g/L sodium bicarbonate, 0.2 mM inositol, 0.1 mM 2-mercaptoethanol, 0.02 mM folic acid, 400 units/mL IL-2, 12.5% horse serum and 12.5% fetal bovine serum.

### Isolation and immortalization of patient derived fibroblasts

Fresh patient pancreatic tissue collected after surgery was digested enzymatically, using our established protocol [38]. Briefly, tissue was initially stored in PBS with antibiotics (P/S, Fungizone) Next, tissue was physically minced with a sterile scalpel, and was digested overnight in collagenase (2mg/mL). After a series of filtration and centrifugation steps, cells were plated on gelatin coated plates for subsequent expansion. Cells were characterized as done previously [37, 38], and fibroblasts were confirmed by a lack of cytokeratins, expression of vimentin, cell shape, ability to create substantive ECM, and the presence of lipid droplets (tumor adjacent fibroblasts) or acquisition of lipid droplets after TGF-β signaling inhibition (CAFs). For the immortalization, cells were retrovirally transduced as previously described [38], with the pBABE-neo-hTERT vector, which was a gift from Dr. Robert Weinberg (Addgene plasmid # 1774). In this study, the procedure above generated the following lines: TA (tumor adjacent fibroblasts) and CAFs (CAF1 and CAF2), which were subsequently used in the 3D system (outlined below) or *in vivo* (CAF1 CON and NetG1 KO).

### Generation of Patient Derived PDAC cell lines from patient derived xenographs (PDX)

Procedure was performed with an approved protocol from Fox Chase Cancer Center Institutional Review Board and was done as previously described [100]. Briefly, tumor fragments were obtained after surgery and washed in RPMI media. The washed fragments were then resuspended in 1:1 mixture of RPMI and Matrigel on ice and were implanted subcutaneously into 5-8 week old *C-B17.scid* mice (Taconic Bioscience). Once tumors developed to a size of at least 150 mm^3^, but no larger than 1,500 mm^3^, mice were sacrificed, and the tumors were collected. These tumors were then digested with a solution containing collagenase/hyaluronidase, and dispase (StemCell Technologies, Cambridge, MA). After several washes, dissociated PDX cells were plated in DMEM supplemented with 10% FBS, 1% penicillin/streptomycin, and 2 mM glutamine. All of the following reagents were from Thermo Fisher Scientific (Waltham, MA) and supplemented into the media: F12 nutrient mix, 5 µg/mL insulin, 25 µg/mL hydrocortisone, 125 ng/mL EGF, 250 µg/mL fungizone, 10 µg/mL gentamycin,10 mg/mL nystatin, 11.7 mmol/L cholera toxin, and 5 mmol/L Rho kinase inhibitor Y27632 (Sigma-Aldrich, St. Louis, MO). After 1 month of passages, cells were lysed and subjected to western blot analysis for the detection of NGL-1 expression.

### Generation of immortalized murine fibroblasts

Murine fibroblasts were isolated from pancreas by enzymatic digestion and clonal selection. Briefly, mouse pancreas was minced in 0.125mg/ml Collagenase and 0.125mg/ml Dispase in DMEM supplemented with 1% FBS and 1x Pen-Strep. The mixture was transferred to 15 mL tube and left at room temperature for 1-2 minutes to separate and aspirate the fat floating on the top. Subsequently, the tissue was digested by incubation for about 1.5-2 hours at 37°C with gentle agitation. Every 10-15 minutes digestion media was replaced with a fresh batch and pancreas tissue was pipetted to break down big chunks. Digested tissue was plated in DMEM supplemented with HEPES, 10% FBS and 1x Pen-Strep. Media was replaced twice a week. Clonal selection, immortalization, and phenotype confirmation was performed as in the above section “Isolation and immortalization of patient derived fibroblasts”.

### 3D microenvironment preparation (Cell-derived ECM (CDM) production)

Production of cell-derived ECM (often referred to as CDM) was conducted as previously described [38]. Briefly, CON or NetG1 KO CAFs (1.25×10^5^/mL) were seeded onto crosslinked (1% (v/v) glutaraldehyde then 1M ethanolamine) 0.2% gelatin coated wells of a 24 well or 6 well plate. These cells were grown in DMEM supplemented with 15% FBS, 1% Penicillin-Streptomycin (10,000 U/mL), 4mM glutamine, and 50 µg/mL L-ascorbic acid (Sigma-Aldrich, St. Louis, MO), provided fresh, daily. After 5 days of ECM production, cultures were used to collect secreted factors (described below for conditioned media preparation) or cells were lysed and extracted from ECMs using 0.5% Triton X-100 (Sigma-Aldrich, St. Louis, MO) supplemented with 20 mM Ammonia Hydroxide (Sigma-Aldrich, St. Louis, MO), followed by extensive washes with PBS. The resulting cell-free ECMs were used as the 3D scaffold environment for subsequent functional assays.

### Conditioned media (CM) preparation

Fibroblast CM was prepared by growing cells during 3D CDM production (see above) for 5 days and then replacing media with glutamine/serum free (-) media. Cells were allowed to condition the media for 2 days. The CM was then collected and used immediately or aliquoted and frozen at −80°C until use (for up to one month).

### mRNA Microarray

RNA was isolated from two sets of patient matched tumor adjacent fibroblasts and tumor associated fibroblasts (CAFs) at the end of CDM production (see above), which were generated in two technical repetitions. The RNA was processed by the Genomics Facility at Fox Chase Cancer Center to render differentially expressed genes between the two sets of matching tumor adjacent fibroblasts and CAFs. Briefly, an Agilent Technologies Gene Chip (∼44,000 genes) was used and RNA was hybridized to the chip and run through the Agilent Technologies Instrument (S/N: US22502680). Raw data were processed using the Agilent Feature Extraction 9.5.1.1 Software, outputting processed data into excel spreadsheets. Quantile normalization was applied to microarray data. We applied limma [101] to identify differentially expressed genes between TA and CAFs. Top differentially expressed genes were selected based on p-value < 0.01 and depicted as a heatmap (z-scores). In order to identify pathways that are enriched between these two cell types, we applied GSEA (preranked with default parameters) on significantly differentially expressed genes (p-value < 0.05) with no fold-change cutoff. To depict the resulting enriched pathways (FDR < 0.25) we applied enrichment map method [102]. The full data set, the top hits with p < 0.01, and GSEA list with FDR <0.25, and complete GSEA analysis for positive and negative enrichment, are all available in Supplemental Table 1, organized by tabs.

### Quantitative Reverse Transcriptase Polymerase Chain Reaction (qRT-PCR)

The Ambion PureLink kit (Life Technologies, Carlsbad, CA) was used, following manufacturer’s instructions, to extract total RNA cells from the various experimental conditions and tested for RNA quality using a Bioanalyzer (Agilent, Santa Clara, CA). Contaminating genomic DNA was removed using Turbo DNA free from Ambion. RNA concentrations were determined with a spectrophotometer (NanoDrop; Thermo Fisher Scientific, Waltham, MA). RNA was reverse transcribed (RT) using Moloney murine leukemia virus reverse transcriptase (Ambion, Life Technologies, Carlsbad, CA) and a mixture of anchored oligo-dT and random decamers (Integrated DNA Technologies, Skokie, Illinois). Two reverse-transcription reactions were performed for each biological replicate using 100 and 25 ng of input RNA. Taqman assays were used in combination with Taqman Universal Master Mix and run on a 7900 HT sequence detection system (Applied Biosystems, Foster City, CA). Cycling conditions were as follows:

95°C, 15 minutes, followed by 40 (two-step) cycles (95°C, 15 s; 60°C, 60 s).

Ct (cycle threshold) values were converted to quantities (in arbitrary units) using a standard curve (four points, four fold dilutions) established with a calibrator sample. Values were averaged per sample and standard deviations were from a minimum of two independent PCRs. Polymerase (RNA) II (DNA directed) polypeptide F (POLR2F) was used as internal control. Identification numbers of commercial assays (Life Technologies, Carlsbad, CA) and sequences of the NTNG1 primers (FW and RV) and probe for the POLR2F assay are as follows:

GGAAATGCAAGAAGAATTATCAGG (forward),

GTTGTCGCAGACATTCGTACC (reverse),

6fam-CCCCATCATCATTCGCCGTTACC-bqh1 (probe)

For other genes, the same RNA extraction was performed, but RT reaction was performed using High Capacity cDNA RT Kit and qRT-PCR reactions were performed in a barcoded 96 well plate using Power SYBR Green PCR master mix, gene specific primers, and template DNA, using the StepOnePlus Real Time-PCR system. All reagents and instruments were from Applied Biosystems/Thermo Fisher Scientific, Waltham MA. All primer sequences were obtained from PrimerBank [103] and are available in the Key Resources Table. Relative mRNA expression for each gene was calculated by the ΔΔCt method, with 18s used as a housekeeping control, and then normalized to CON CAFs to determine fold change differences in gene expression.

### SDS-PAGE and Western Blot Analysis

Cell lysates were prepared by adding 150 µL standard RIPA buffer to 6 well plates containing 10^6^ cells. The lysates were homogenized using a sonicator. After centrifugation, the supernatant was collected and 30 µg of lysate (determined by the Bradford Assay) was diluted in 2x loading buffer (Bio-Rad, Hercules, CA) and was loaded onto 4-20% polyacrylamide gels (Bio-Rad, Hercules, CA). The gels were run at 70 volts for 2 hours, and transferred to PVDF membranes (Millipore, Burlington, MA) using the semi-dry transfer system (Bio-Rad, Hercules, CA). Membranes were blocked in 5% nonfat milk in 0.1% Tween in tris buffered saline (TBST) for 1 hour at room temperature (RT), and primary antibodies were incubated with the membranes overnight at 4°C. Next, membranes were washed 5 times with 0.1% TBST and then secondary antibodies linked to HRP were diluted in 5% nonfat milk in 0.1% TBST and were applied for 2 hours at RT. Membranes were washed again as before, and incubated for 5 minutes in Immobilon Western Chemiluminescent HRP substrate (Millipore, Burlington, MA). Membranes were developed using film and scanned to create digital images. Primary and secondary antibodies are listed in the Reagents table.

### Indirect Immunofluorescence

CAFs were allowed to generate ECM (3D microenvironment) as outlined above (CDM production). After 5 days of ECM production, cells on coverslips were fixed and simultaneously permeabilized for 5 minutes at RT in 4% paraformaldehyde containing 0.5% TritonX-100 and 150 mM sucrose and allowed to continue fixation, using the same preparation lacking triton, for an additional 20 minutes at RT. Fixed/permeabilized cells were then blocked for 1 hour at RT with Odyssey Blocking Buffer (PBS base; Li-Cor, Lincoln, NE). Cells were stained for 1 hour at RT with the following antibodies diluted in blocking buffer: fibronectin (Sigma-Aldrich, St. Louis, MO) and α-smooth muscle actin (α-SMA) (Sigma-Aldrich, St. Louis, MO). After 1 hour incubation, cells were washed 3 times with 0.05% Tween in phosphate buffered saline (PBST), and secondary antibodies diluted in blocking buffer were applied for 1 hour at RT (TRITC for α-SMA; Cy5 for fibronectin, Jackson ImmunoResearch, West Grove, PA). Next, cells were washed 3 times with 0.05% PBST, and nuclei were counter stained with SYBR green (1:10,000; Invitrogen, Carlsbad, CA). Stained coverslips were imaged using Spinning disk confocal microscopy (Perkin Elmer, Waltham, MA), captured by a CoolSNAP CCD Camera (Photometrics, Tucson, AZ) attached to an Eclipse 2000-S inverted microscope (Nikon, Tokyo, Japan). Images were exported from Volocity 3D Image Analysis Software (Perkins Elmer, Waltham, MA). The integrated intensity of α-SMA was quantified using Metamorph Image Analysis Software (Molecular Devices, San Jose, CA). Fibronectin alignment was quantified as described below.

### ECM fiber orientation analysis

ECM fiber orientation, indicative of alignment, was performed as previously described [38]. Briefly, confocal images of fibronectin in CAF generated ECM’s were analyzed for fiber orientation using the OrientationJ plugin [104] for ImageJ. Fiber angles generated by the plugin were normalized to fit within the angle range of −90° to +90°. Next, normalized angles were plotted in Excel to generate orientation curves (alignment output at every angle). Finally, the percentage of fibers between −15° to +15° was plotted as a measurement indicative of ECM fiber alignment. In general, tumor adjacent fibroblasts tend to have disorganized −15° to +15° angle fibers (<50%), while CAFs tend to have fibers with angles >50%.

### PDAC Cell Proliferation Assay

CON or NGL-1 KO PDACc cells (2×10^4^) were seeded in CAF-derived 3D ECMs and allowed to proliferate overnight. Ki67, a nuclear marker of proliferation, and nuclei (DAPI) were stained by IF (as outlined above). The number of Ki67 positive cells/nuclei per field was calculated using Metamorph image analysis software.

### CRISPR/Cas9 Mediated Knockout of NetG1 and NGL-1

Knockout of NetG1 in CAFs, or NGL-1 in PDACc or PANC-1 cells, was achieved by introducing a frameshift mutation in coding regions of each gene, induced by cuts made by CRISPR/Cas9 and gene specific gRNAs introduced into cells by lentiviral transduction.

#### gRNA design

Using the MIT Optimized CRISPR Design website: http://crispr.mit.edu/, the first 200 bp of exon 5 for NetG1 (first common exon for all splice variants) and first 220 bp of the NGL-1 gene were used to generate gRNAs. The top two hits for each gene were selected and designed to include BsmBI/Esp3I overhangs (bold, underline, italics), the specific gRNA sequence (plain text). G (italics, underline) was added to the .1 gRNAs (and corresponding C on the .2 gRNA) to optimize its expression for the human U6 promoter.

The gRNA sequences are listed below:

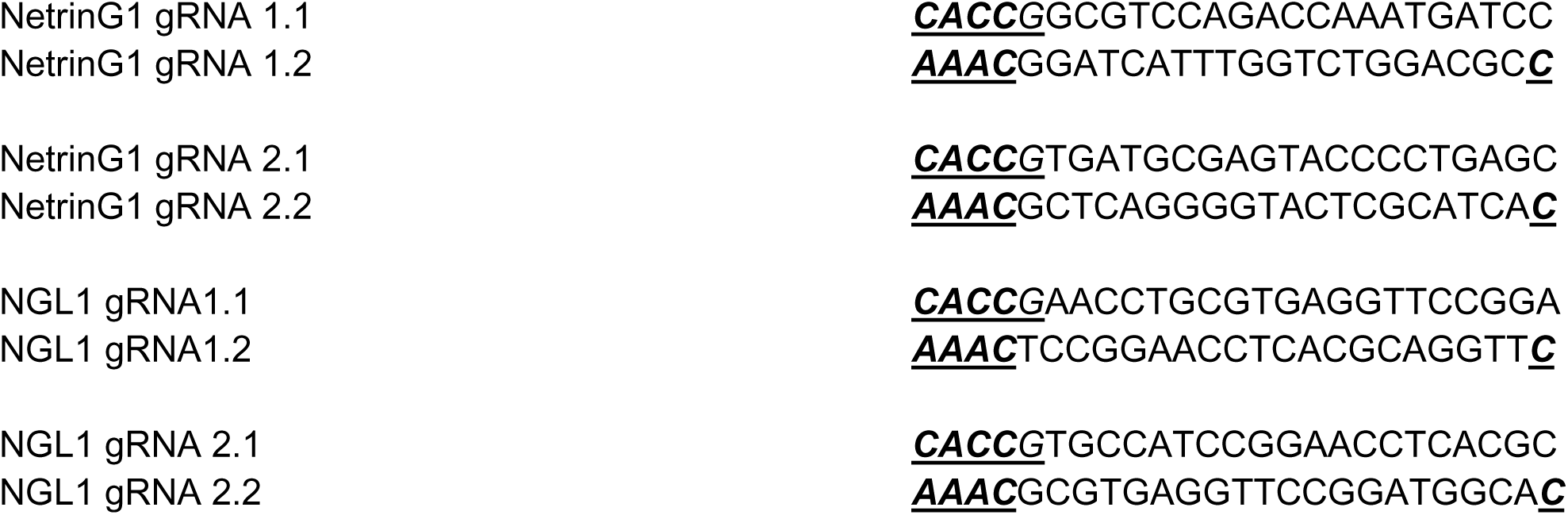

Empty vector or Nontargeting gRNA against eGFP was used as a control:

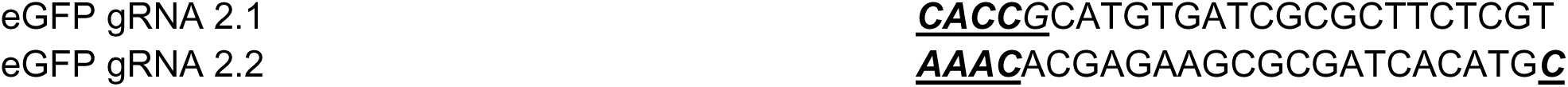

#### Generation of Lentiviral Vectors

Designed oligos were ordered from Integrated DNA Technologies (Skokie, Illinois) and cloned into the LentiCRISPR v2 vector, as previously published (a gift from Feng Zhang; Addgene plasmid # 52961; [38, 105]. Briefly, oligos were phosphorylated using T4 PNK (New England Biolabs, Ipswich, MA) and annealed in a thermal cycler. Annealed oligos were cloned into a gel purified (GeneJet Gel Extration Kit; Thermo Fisher Scientific, Waltham, MA), dephosphorylated (Fast AP; Thermo Fisher Scientific, Waltham, MA), and digested vector (Fast Digest Esp3I; Thermo Fisher Scientific, Waltham, MA). Annealed oligos were ligated into the digested vector using quick ligase (New England Biolabs, Ipswich, MA). The ligated vectors were transformed into Stbl3 strain of *Escherichia coli* (Thermo Fisher Scientific, Waltham, MA) and selected with 100 µg/mL ampicillin on LB/agar plates. The following day, single colonies of bacteria were screened by colony PCR using a forward primer for the U6 promoter (GAGGGCCTATTTCCCATGATT) and the reverse primer sequence (gRNA x.2) for each gene. Positive clones were subjected to sequencing and subsequently expanded for plasmid purification and lentivirus production.

#### Lentiviral Transduction

To package the LentiCRISPRv2 plasmid into functional virus particles, 10 µg LentiCRISPRv2, 5 µg psPAX2 (psPAX2 was a gift from Didier Trono; Addgene plasmid # 12260) and 2 µg VSVg were mixed with 30 µL X-tremeGene9 transfection reagent (Sigma-Aldrich, St. Louis, MO) in 1 mL of serum free/antibiotic free DMEM and held at RT for 30 minutes. The transfection mixture was added dropwise to 293T cells containing 5 mL serum free/antibiotic free DMEM and the cells were incubated at 37°C overnight. The next day, the media was replaced with complete DMEM media. On days 2 and 4 post-transfection, the media with viral particles was collected and syringe filtered through a 0.45 µM filter (Millipore, Burlington, MA). The filtered viral supernatant was used immediately to transduce target cells (PDACc/PANC-1 or fibroblasts) with 10 µg/mL polybrene (Santa Cruz Biotechnology, Dallas, Texas) or stored at −80°C for use in the future. Target cells were selected with puromycin (2 µg/mL for CAFs, 5 µg/mL PANC-1, 12 µg/mL KRAS) for 2 weeks. Cells that survived selection were used for future experiments.

#### Generation of RFP+ and GFP+ CRISPR/Cas9 transduced cells

eGFP and mCherry (RFP) were PCR amplified using Phusion HF polymerase master mix (Thermo Fisher Scientific, Waltham, MA) to add overhangs compatible with restriction enzymes. Then cloned into XbaI/XhoI (New England Biolabs, Ipswich, MA) digested, Fast AP dephosphorylated pLV-CMV-H4-puro vector (kindly provided by Dr. Alexey Ivanov, West Virginia University School of Medicine, Morgantown, WV, USA). Empty vector transduced (CON) or NetG1 KO CAF were transduced with pLV-CMV-H4-puro-GFP and the CON or NGL-1 KO PDACc were transduced with pLV-CMV-H4-puro-RFP, with 10 µg/mL polybrene. Cells were selected by flow cytometry. The selected cells were used for future experiments.

### CRISPR interference (CRISPRi) mediated knockdown of target genes

We generated a lentiviral vector named CRISPRi-Puro (modified from pLV hU6-sgRNA hUbC-dCas9-KRAB-T2a-Puro; Addgene Plasmid #71236; a gift from Charles Gersbach) [106] containing dead Cas9 (dCas9) fused to Krueppel-associated box (KRAB) domain, gRNA cloning site, and puromycin resistance. The gRNA cloning site was modified to contain a “stuffer” (cut from LentiCRISPRv2 plasmid using the restriction enzyme BsmBI and ligated into our vector) that resulted in a higher gRNA cloning efficiency. Upon insertion of gene specific gRNA, dCas9-KRAB mediated knockdown of target genes could be achieved. Three gRNA oligos were selected from the top ten gRNA sequences generated from the study “Compact and highly active next-generation libraries for CRISPR-mediated gene repression and activation” [107]. Generation of the vector and transduction of the target cells was performed as described above for the generation and transduction of the LentiCRISPRv2 vector. The best two knockdown cell lines generated were subsequently used for all experiments. Cells transduced with vector alone were used as a control.

The selected oligos are listed below:

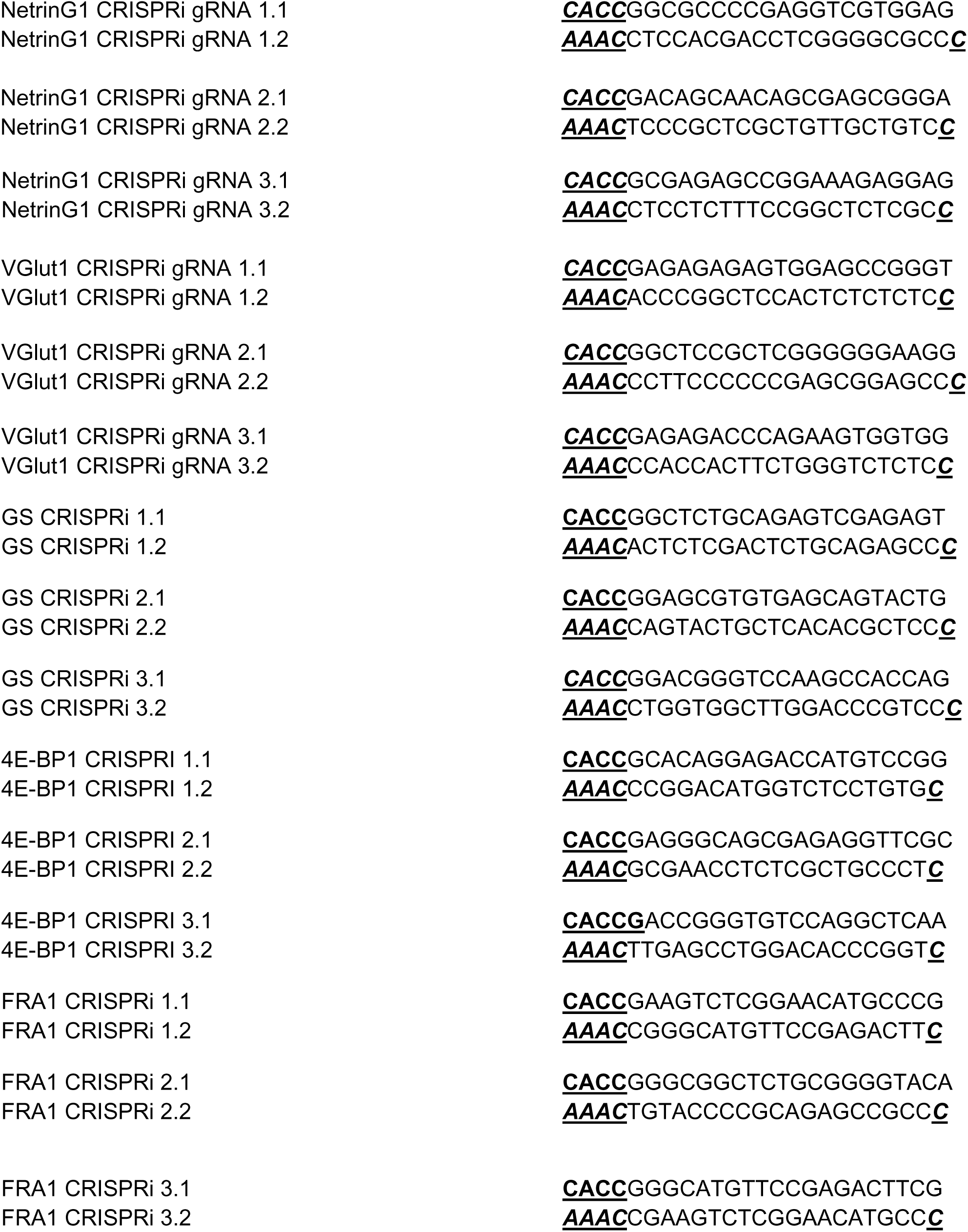

### Cell Engagement/Motility Assays

RFP+ CON or NGL-1 KO PDACc (5×10^4^) were plated with GFP+ CON or NetG1 KO CAFs (5×10^4^) in 3D and allowed to attach for 2 hours. Following cell attachment, 3 regions per well, where there was obvious clustering of red PDACc and green CAFs, were selected to be imaged every 15 minutes for 24 hours, using a motorized XYZ stage (Optical Apparatus Co., Ardmore, PA) controlled by MetaMorph 6.3r7 (Molecular Devices, Downingtown, PA) software. Time-lapse movies were built with images from each corresponding fluorescence channels acquired every 15 min for a period of 24 hours using either 4X or 20X Pan Fluor objectives mounted on a Nikon TE-2000U wide field inverted microscope (Optical Apparatus Co., Ardmore, PA) with a Cool Snap HQ camera rendering 3 time-lapse movies per condition for a total of 9 videos generated over 3 independent experimental repetitions. The resulting time-lapse videos were analyzed to quantify heterotypic cell engagement (yellow interacting areas using both channels simultaneously in merged file stacks) and PDACc cell motility (using only the monochromatic images corresponding to the red channel), as outlined below.

#### Cell Engagement

For image analysis MetaMorph offline 7.8.1.0 software was used. To achieve sequence coherence, out of focus or blank images from particular fluorescence channels were removed, together with its fluorescence counterpart image. All movies were normalized (trimmed) to 18 hours data for analysis. Images were scaled and pseudocolored (RFP in red, GFP in green) similarly for all conditions. After merging of fluorescence channels, cell engagement areas (CEA) obtained from regions of fluorescence colocalization (in yellow) were digitally detected, isolated and measured for each condition over time.

#### PDAC Cell Motility

Monochromatic channel of RFP+ PDAC cells were tracked using the Track Object function in MetaMorph 7.8.1.0 offline software (Molecular Devices, Downingtown, PA). A minimum of 5 cells per region were chosen to carefully follow. Red PDACc that were touching other red PDACc, moving out of the imaging plane, or that began to round up prior to mitosis were not counted in this experiment. Motility was analyzed according to the total distance (µm) traveled over the timespan of the experiment (24 hr), expressed as velocity (µm/hr).

### Material Transfer Assay

2×10^4^ RFP+ CON or NGL-1 KO PDACc and/or 2×10^4^ GFP+ CAFs were seeded in 3D and were co-cultured for 36 hours, in serum free DMEM. Spinning disk confocal images (Perkin Elmer, Waltham, MA) were captured by a CoolSNAP CCD Camera (Photometrics, Tucson, AZ) attached to an Eclipse 2000-S inverted microscope (Nikon, Tokyo, Japan). Images were exported from Volocity 3D Image Analysis Software (Perkins Elmer, Waltham, MA) and quantified for the integrated intensity of GFP+ pixels under RFP+ pixels, using Metamorph Image Analysis Software (Molecular Devices, San Jose, CA), which was indicative CAF derived GFP material transfer to RFP+ PDACc.

### PDAC Survival Assay

2×10^4^ RFP+ CON or NGL-1 KO PDAC cells (PDACc or PANC-1) and/or 2×10^4^ GFP+ CON or NetG1 KO CAFs were seeded in 3D (CON CAF, NetG1 KO, or tumor adjacent fibroblast ECMs, as indicated) in 24 well plates. Cells were co-cultured for 96 hours in glutamine free/serum free DMEM. Next, at least 10 images were taken of red PDAC cells and green CAFs using a Cool Snap 1HQ camera (Roper Scientific, Vianen, Netherlands) on an Eclipse 2000-U inverted microscope (Nikon, Tokyo, Japan). PDAC cell survival was quantified as the number of RFP+ per field of view, using Image J. Dead cells were excluded by Sytox Blue (1 µM; Thermo Fisher Scientific, Waltham, MA) positivity or cell size. Briefly, identical thresholds were established for each color channel for each batch of images per experiment. Then the images were batch processed using a macro that measures cell counts and % area coverage of cells in each image, at a given threshold. The mean RFP+/Sytox blue negative cells (live) from each condition was calculated (counts or % area, depending on the experiment) and normalized to the CON PDAC/CON CAF condition. Cell counts were chosen when cells were clearly isolated (accurate cell counts), and % area was chosen when cells had more overlap (could not accurately determine cell counts). The 96 hour timepoint was determined empirically, when cell death became evident by eye. Other PDAC survival assays were performed with CAF conditioned media (CM), VGlut1, GS, 4E-BP1, and FRA1 KD CAFs, or CAFs pretreated 24 hours with 1mM L-Methionine sulfoximine (glutamine synthetase inhibitor; Sigma-Aldrich, St. Louis, MO), 10 nM SB 202190 (P38 inhibitor; Tocris Bioscience, Bristol, UK), 10 µM MK-2206 (AKT inhibitor; Cayman Chemical, Ann Arbor, MI), or 2.5 µg/mL NetG1 mAb (Santa Cruz, Dallas, TX), as indicated.

### NK-92 cell killing assay

2×10^4^ RFP+ CON or NGL-1 KO PDAC cells and/or 2×10^4^ GFP+ CON or NetG1 KO CAFs were seeded in 3D in 24 well plates and were co-cultured in 500 µL NK-92 media without IL-2 to establish their niche. The following day, wells were seeded with 8×10^4^ resting or active NK-92 cells, in 500 µL of NK-92 media without IL-2. Cells were co-cultured for 48 hours and PDAC survival was assessed identically to the PDAC cell survival assay. To produce active NK-92 cells, cells were maintained with 400 units/mL IL-2 and for resting NK-92 cells, cells were maintained for one week in 100 units/mL IL-2 and switched overnight to NK culture media containing no IL-2 the day before the experiment. For 2D conditions, all experimental conditions were the same, except the endpoint of the experiment was at 6-8 hours, as lack of CAF derived ECM (3D microenvironment) allowed NK-92 cells to kill at a faster rate. For the IL-15 neutralizing experiments, an IL-15 neutralizing antibody (10 µg/mL; R&D Systems, Minneapolis, MN) or Nonspecific IgG isotype control (10 µg/mL; Jackson Immunoresearch, West Grove, PA) were added to the co-cultures where specified. Whenever CON or NetG1 KO CAF CM was used, it was diluted 1:1 in NK-92 media without IL-2. Images were quantified for live PDAC cells as in the “PDAC Survival Assay” section above.

### NK-92 cell activation assay

8×10^4^ IL-2 activated NK-92 cells were directly co-cultured (CC) with CAFs or with conditioned media (CM) derived from CAFs for 2 hours. After the co-culture period, NK-92 activation status was determined by staining Granzyme B intracellularly, or by using the IFNγ secretion assay detection kit. Stained cells were subjected to flow cytometry using by LSRII (Beckton Dickinson, San Jose, CA) and analyzed with FlowJo 9.9 software (Tree Star, San Carlos, CA). IL-2 activated NK-92 cells cultured alone were used as the positive control. Percentage of cells that were positive for either Granzyme B or IFNγ were considered activated.

### Enzyme-linked immunosorbent assay (ELISA)

ELISAs for IL-15, IFN-β, TGF-β, IL-6, IL-8, GM-CSF, mTGF-β, mGM-CSF, and mIL-6 (R&D Systems, Minneapolis, MN), as well as glutamate (Abcam, Cambridge, UK) and glutamine (MyBioSource, San Diego, CA), were performed as recommended by the manufacturers. Briefly, buffers for the IL-15, IFN-β, IL-6, IL-8, GM-CSF, mGM-CSF, and mIL-6 ELISAs were prepared using the DuoSet ELISA Ancillary Reagent Kit 2, and buffers were prepared using the DuoSet ELISA Ancillary Reagent Kit 1 for the TGF-β and mTGF-β ELISAs (R&D Systems, Minneapolis, MN). Standards, CM or lysates from various fibroblastic lines or murine tumor cultures were incubated for 2 hours on plates pre-coated overnight with capture antibody. For the TGF-β ELISAs, samples first underwent activation using the Sample Activation Kit 1 (R&D Systems, Minneapolis, MN) before sample incubation. Next, detection antibody linked to biotin was added to the wells and incubated for 2 hours. Streptavidin-HRP was then added to the wells and incubated for 20 minutes. Samples were rinsed 3 times with wash buffer in between all steps. Substrate was added for 20 minutes and was neutralized with stop buffer. Plates were read using Spark® Multimode Microplate Reader (Tecan, Switzerland) at 450 nm to measure the specified protein present in sample. Plate imperfections, measured at 570 nm, were subtracted from the readings at 450 nm, to produce corrected readings. Finally, media alone (BLANK) was subtracted from the corrected readings for each sample. To determine the concentration of the indicated protein in each sample, a standard curve was generated using known concentrations of analyte, and the blank corrected values determined for each sample were compared to the standard curve.

### U-Plex Assay

U-Plex Assay (multiplex ELISA) was carried out as recommended by the manufacturer (Meso Scale Discoveries, Rockville, MD). Briefly, TA and CON or NetG1 KO CAF CM samples were generated and added to a pre-coated plate containing 10 capture antibodies per well for the detection of the following proteins: IL-1β, IL-2, IFNγ, GM-CSF, IL-6, IL-8, IL-10, IL-12p70, MIP3α, and TNFα. After 3 washes, sample, standards, or calibrator were added to the wells and incubated for 1 hour. Next, after 3 washes, detection antibody was added to the wells and incubated for 1 hour. After a final set of 3 washes, read buffer was added to the wells and the plate was read using the MSD SECTOR Imager 2400 (Meso Scale Discoveries, Rockville, MD) and the data was processed using the MSD Workbench 4.0 software (Meso Scale Discoveries, Rockville, MD).

### Amino Acid Screen: Biochrom Method

*Media*: 20 µL of media was mixed with 80 µL of water and 100 µL of 10% Sulfosalicylic acid (Sigma-Aldrich, St. Louis, MO) followed by incubation for 1 hour at 4°C and then centrifugation at 13,000 rpm (15 min; 4°C). No visible pellet was seen. Supernatant was filtered using Ultrafree-MC-GV centrifugal filters (Millipore, Burlington, MA) and used for amino acid analysis employing Biochrom 30 amino acid analyzer. Results are in terms of µM.

*Cell Lysates*: 14.55 µg of cell lysate protein in RIPA buffer was brought to a total volume of 50 µL in water and was mixed with 50 µL of 10% Sulfosalicylic acid (Sigma-Aldrich, St. Louis, MO) followed by incubation for 1 hour at 4°C and then centrifugation at 13,000 rpm (15 minutes; 4°C). Supernatant was filtered using Ultrafree-MC-GV centrifugal filters (Millipore, Burlington, MA) and used for amino acid analysis employing the Biochrom 30 amino acid analyzer (Biochrom US, Holliston, MA). Results were provided in nmol/mg protein units.

These results are available in Supplemental Tables 7-9.

### Amino Acid Screen: Mass Spectrometry

#### Metabolite Extraction and Quantitation

Metabolites were extracted from media and quantitated as described previously [66]. 10μL of media was mixed with 10μL of a mixture of isotopically labeled metabolites and amino acids of known concentrations (Cambridge Isotope Laboratories, Tewksbury, MA) and extracted with 600μL ice-cold HPLC grade methanol. Samples were vortexed for 10 minutes at 4°C, centrifuged at max speed on a tabletop centrifuge for 10 minutes 4°C, and then 450uL of supernatant was dried under nitrogen gas and frozen at −80°C until analyzed by GC-MS.

#### Gas Chromatograpy-Mass Spectrometry Analysis

Polar metabolites were analyzed by GC-MS as described previously [66]. Dried metabolite extracts were derivatized with 16μL MOX reagent (Thermofisher, Waltham, MA) for 90 minutes. at 37°C, followed by 20 μL of N-tertbutyldimethylsilyl-N-methyltrifluoroacetamide with 1% tert-butyldimethylchlorosilane (Sigma Aldrich, St. Louis, MO) for 60 minutes. at 60°C. After derivatization, samples were analyzed by GC-MS using a DB-35MS column (Agilent Technologies, Santa Clara, CA) installed in an Agilent 7890A gas chromatograph coupled to an Agilent 5997B mass spectrometer. Helium was used as the carrier gas at a flow rate of 1.2 mL/minute. One microliter of sample was injected in split mode (all samples were split 1:1) at 270°C. After injection, the GC oven was held at 100°C for 1 minute. and increased to 300°C at 3.5 °C/minute. The oven was then ramped to 320°C at 20 °C/minute and held for 5 minutes. at this 320°C. The MS system operated under electron impact ionization at 70 eV and the MS source and quadrupole were held at 230°C and 150°C respectively. The detector was used in scanning mode, and the scanned ion range was 100–650 m/z.

These results are available in Supplemental Table 10.

### Human NK cell Isolation and Activation Experiments

#### Isolation of Primary NK cells

Blood was collected into heparinized tubes from healthy volunteer donors that consented using HIPAA-compliant procedures approved by the Institutional Review Board of Fox Chase Cancer Center. NK cells were enriched from healthy donor PBMCs using the EasySep NK cell negative selection kit (StemCell Technologies, Cambridge, MA) and confirmed to be >95% purity using flow cytometry.

#### Primary NK Cell Activation Assay

Purified NK cells were incubated overnight at 37°C in indicated percentages of CON or NetG1 CAF CM diluted in NK cell media (10%-65%) in the presence of IL-2 (500 units/mL) and IL-12 (10 ng/mL). NK cells activated with IL-2 and IL-12 were used as a positive control. After overnight incubation, the population of IFNγ secreting cells was identified using the IFNγ Secretion assay detection kit (PE) (Miltenyi Biotec, Bergisch Gladbach, Germany) according to the manufacturer’s protocol. After a 2 hour incubation with IFNγ catch reagent, cells were washed twice. Next, cells were stained with IFNγ detection antibody and antibodies against NKp80 (anti-NKp80-APC, Biolegend, San Diego, CA), CD3 (anti-CD3-Cy7APC, BD Pharmigen, San Jose, CA), CD56 (anti-CD56-BUV395, BD Horizon, San Jose, CA), CD69 (anti-CD69-Pac Blue, San Diego, CA). Cells were centrifuged and rinsed twice, with propidium iodide in the second wash. NK cells were defined as NKp80+CD56+CD3- cells.

#### Flow cytometry and data analysis

Stained cells were analyzed on a BD ARIA II flow cytometer (Becton Dickinson, Franklin Lakes, NJ). Data was collected with BD FACS Diva software (v6) and analyzed with FlowJo (v9.9; Tree Star Inc., Ashland, OR). IFNγ and CD69 expression were identified as the percentage of IFNγ+ cells or mean fluorescence intensity (MFI) of CD69+ cells. The data were normalized to the positive control (NK cells with IL2/IL12 treatment alone).

### Mouse Models

Tissue for stromal analysis was obtained from the LSL-**<u>K</u>**RAS^G12D^, PDX1-**<u>C</u>**re (KC) model [46], and KPC4B and KPC3 murine PDAC cell lines were derived from tumors generated from a variant of LSL-**<u>K</u>**RAS^G12D^, T**<u>P</u>**53^-/-^, PDX1-**<u>C</u>**re (KPC) mice [47].

### PDAC cell orthotopic injections

All animal work was performed under protocols approved by IACUC at Fox Chase Cancer Center. A small incision was made in the abdominal skin of the mice, and gently moved aside the spleen to reveal the pancreas. Next, 10^6^ murine CON or NGL-1 KO #1 or #2 PDAC cells (derived from KPC tumors), were injected (100 µL) orthotopically into the pancreas of syngeneic C57BL/6 mice and tumors were allowed to develop for 3.5 weeks. Mice were monitored by observation, palpation, and weekly MRIs and sacrificed at 3.5 weeks. Pancreas weights were obtained and pancreatic tissue was further analyzed as described below.

To determine the effect of the immune system on tumorigenesis in the orthotopic model, the injections were performed as above, with the following mice used as hosts:

C57BL/B6 (immunocompetent; FCCC), C.B17 Severe Combined Immunodeficient, (SCID; lacks T and B cells; Taconic Bioscience), and NOD.Cg-Prkdc^scid^ IL2rg^tm1Wjl^/SzJ: NOD *scid* gamma (NSG; lacks NK, T, and B cells; Jackson Laboratories).

For the mouse tumor cell/normal fibroblast co-injections, 2.5*10^5^ murine PDAC cells (CON or NGL-1 KO) and 7.5*10^5^ normal pancreatic fibroblasts (CON, NetG1 KD1+2, GFP, NetG1 WT, NetG1 Mut) were co-injected into the pancreas of C57BL/B6 mice and tumors were allowed to develop over 3.5 weeks (as above). Murine cancer cell and normal fibroblast injected alone were used as controls.

For the NetG1 neutralizing monoclonal antibody (mAb) experiment, after mice recovered from the tumor cell (10^6^) orthotopic injections for 3 days, mice were divided into two groups (IgG; n=9 or mAb; n=10) and were treated with 5 mg/kg isotype specific IgG (IgG_1_; West Lebanon, New Hampshire) or mAb (Santa Cruz, Dallas, TX) 3 times per week, until the completion of the model (3.5 weeks, as above).

For the human tumor cell/CAF co-injections, RFP^+^ PDACc cells (CON or NGL-1 KO) and CAFs (CON or NetG1 KO) were co-injected at a 1:3 ratio (2.5*10^5^ tumor: 7.5*10^5^ CAF) into SCID mice. CAFs and tumor cells injected alone were used as controls. Tumors were allowed to form for 4 weeks. Subsequently, mice were sacrificed and tissue was analyzed as outline below.

### Magnetic Resonance Imaging (MRI)

Animals were imaged in a 7 Tesla vertical wide-bore magnet, using a Bruker DRX 300 spectrometer (Billerica, MA) with a micro-imaging accessory, which included a micro 2.5 gradient set, a 30 cm radiofrequency coil, and Paravision software (Bruker, Billerica, MA). Animals were anesthetized with a mixture of oxygen and isoflurane (2-3%). An injection of 0.2 mL of 10:1 diluted Magnevist (gadopentetate dimeglumine; Bayer, Whippany, NJ) was made into the shoulder region immediately preceding the scan. Scout scans were performed in the axial and sagittal orientations, permitting us to accurately prescribe an oblique data set in coronal and sagittal orientations that included the organs of interest. A two-dimensional spin echo pulse sequence was employed with echo time 15 msec, repetition time 630 msec, field of view = 2.56 cm, acquisition matrix= 256×256, slice thickness= 0.75 mm, 2 averages, scan time = 5 minutes. Fat suppression (standard on Bruker DRX systems) was used for all scans. Raw data files were converted to Tiff format image files using the Bruker plugin on ImageJ.

### Murine tissue preparation and histopathological analysis

#### Tissue Preparation

Tissues were collected and fixed in 10% phosphate-buffered formaldehyde (formalin) 24-48 hours, dehydrated and embedded in paraffin. Tissues were processed by dehydration in a series of ethanol followed by xylene (70% ethanol 3 hours, 95% ethanol 2 hours, 100% ethanol 2 hours, 100% ethanol Xylene mixed 1hours, Xylene 3hours, then immersed in Paraffin). Paraffin blocks were cut into 5 µm sections, mounted on microscope slides, and stored at room temperature until used. Sections were stained with hematoxylin and eosin for pathologic evaluation. All H&E slides were viewed with a Nikon Eclipse 50i microscope and photomicrographs were taken with an attached Nikon DS-Fi1 camera (Melville, NY, USA).

#### Tumor area analysis

H&E stained slides were also scanned using Aperio ScanScope CS scanner (Aperio, Vista, CA). Scanned images were then viewed with Aperio’s image viewer software (ImageScope). Selected regions of interest for PDAC, necrotic, and normal tissue were outlined respectively by a pathologist (Dr. Kathy Q. Cai), who was blinded to the treatment groups. The ratio of PDAC, necrotic, and normal tissue to the total pancreatic areas scanned was calculated for each animal.

#### Differentiation status of tumor cells

H&E images of pancreata injected with CON or NGL-1 KO cells were scored for the percentage of cells that were well differentiated, moderately differentiated, or poorly differentiated, as determined by a pathologist, who was blinded to the treatment groups.

#### TUNEL Staining

Apoptotic cells were quantified using the DeadEnd™ Fluorometric TUNEL System (Promega, Madison, WI), per manufacturer’s instructions. Briefly, tissue was deparaffinized and fixed in 4% formaldehyde for 15 minutes at RT. After 2 PBS washes, tissue was treated with 20 µg/mL proteinase K solution for 10 minutes at RT. After 2 PBS washes, slides were fixed again in 4% formaldehyde for 5 minutes at RT. Tissue was then submerged in Equilibration buffer at RT for 5 minutes. Next, apoptotic cells were labeled with TdT reaction mixture for 1 hour at 37°C. After 1 hour, the reaction was stopped by adding SSC buffer for 15 minutes at RT. Tissue was then washed 3x in PBS and then stained with DAPI (1:10,000) for 15 minutes at RT. Finally, after an additional wash, tissue was mounted and imaged using the Eclipse 2000-U inverted microscope. Metamorph software was used to quantify dead cells (green)/nuclei (blue) in each field of view. At least 10 images (20X) per sample were acquired.

#### Ki67 staining (proliferation)

IF on tissue sections was done as previously published [38]. Briefly, deparaffinized tissue underwent antigen retrieval by boiling in buffer containing 10 mM Sodium citrate, 0.05% Tween 20, pH 6.0 for 1 hour. Next, tissue was blocked in Odyssey Blocking Buffer (PBS base) (LI-COR Biosciences, Lincoln, NE) for 1 hour at RT. Tissue was then stained with primary antibody against Ki67 (Abcam, Cambridge, UK) overnight at 4°C. The following day, tissue was washed 3 times in 0.05% PBST, and secondary antibody conjugated to TRITC (Jackson ImmunoResearch, West Grove, PA) was applied to the samples for 2 hours at RT. After 3 additional washes, nuclei were stained with DAPI (1:10,000) for 15 minutes at RT. After one final wash, samples were mounted and images (20X) were captured using an Eclipse 2000-U inverted microscope. Proliferating cells (red)/nuclei (blue) in each field of view were quantified using Metamorph Software.

### Supernatant Collection of Tumor Cultures

Small pieces of tissue (30-60 mg) were excised from the pancreata isolated from mice treated with IgG or mAb (described above) and were cultured in serum/Gln free media overnight. The following day, the media was centrifuged for 15 minutes at 15,000 rpm to remove cellular debris. The resultant supernatant was collected and subjected to ELISAs to measure levels of Glu, Gln, mTGF-β, mGM-CSF, and mIL-6.

### Acquisition and Use of Human Tissue

All human tissues used in this study were collected using HIPAA approved protocols and exemption-approval of the Fox Chase Cancer Center’s Institutional Review Board, after patients signed a written informed consent in which they agreed to donate surgical specimens to research. To protect patient’s identity, samples were classified, coded and distributed by the Institutional Biosample Repository Facility. Normal pancreatic tissue was obtained through US Biomax Inc. (Rockville, MD) following their rigorous ethical standards for collecting tissue from healthy expired donors.

### Tissue Microarray (TMA)

#### Acquisition of the TMAs

The FCCC tissue microarray was generated by the Biosample Repository at Fox Chase Cancer Center. There were 80 patient samples generated from three independent TMAs. The MDA TMA was generated at the University of Texas MD Anderson Cancer Center. There were 143 patient samples generated from three independent TMAs. The clinical parameters for each TMA are available in Supplemental Tables 2 and 3, respectively.

#### Immunohistochemistry (IHC)

TMA sections were cut from formalin fixed paraffin embedded (FFPE) blocks. After deparaffinizing in xylene, sections were hydrated in graded alcohol and quenched in 3% H_2_O_2_. Slides were subjected to heat induced epitope retrieval in citrate buffer (pH 6.0) for 1 hour. After cooling to room temperature, sections were permeabilized in Triton X-100 for 20 minutes, followed by a 1 hour incubation in blocking buffer (1%BSA, 2% Normal Goat Sera, 0.2% Cold water fish skin gelatin in PBS). Antibodies against NGL-1 (GTX 121508), NetG1 (GTX 115637), GS (Abcam), VGlut1 (ThermoFisher), and NKp46 (R&D Systems, AF1850) were used at 1:100 dilution and incubation was carried out overnight at 4° C. The sections were washed and incubated with anti rabbit-HRP (Vector Lab MP-7451) and then treated with DAB (ImmPACT SK-4105) for 2.5 minutes. Sections were counter stained in Mayer’s Hematoxylin to show nuclei, dehydrated in graded alcohol and mounted in Cytoseal-60 after clarifying in toluene.

#### Blind Pathological Scoring of TMAs

A pathologist (Dr. Klein-Szanto) blindly scored the 80 and 143 stained TMA cores for fibroblastic NetG1 or tumoral NGL-1 on a scale of 0-4, which is based on staining intensity and coverage. NKp46 was also scored on a 0-4 scale, but with each value representing # of cells staining positive for NKp46, per core. 0 = 0 cells, 1 = 1-10 cells, 2 = 11-20 cells, 3 = 21-30 cells, and 4 = >30 cells. Each patient had two cores stained per marker and these were averaged together to generate the IHC score. These values, along with overall survival data, are available in Supplemental Tables 4 and 5 for FCCC and MDA TMAs, respectively.

#### Generation of Kaplan-Meier Plots

Kaplan-Meier (KM) plots were generated for overall survival correlated with immunohistochemical staining of fibroblastic NetG1, tumoral NGL-1, and NKp46.

### Simultaneous Multichannel Immunofluorescence (SMI)

#### Qdot antibody conjugation

Corresponding antibodies were linked to Qdot molecules using SiteClick Qdot labeling from Molecular Probes-ThermoFisher Scientific. In brief, approximately 100 mg of specific IgG were pre-purified to remove excessive carrier proteins (if needed) and each conjugation was performed following manufacturer’s three-step protocol requiring antibody’s carbohydrate domain modifications to allowing an azide molecule attachment to facilitate linking DIBO-modified nanocrystals to the antibodies.

#### Formalin fixed and paraffin embedded (FFPE) tissue processing

FFPE sections were deparaffinized using xylene and rehydrated in progressive ethanol to water dilutions. Tissues then were processed through heat-induced epitope retrieval using pH 6.0 Citrate buffer, followed by permeabilization with 0.5% TritonX-100. After treating the specimens with Odyssey (PBS base) blocking buffer (LI-COR Biosciences, Lincoln, NE) samples were labeled by a three-day indirect immunofluorescence scheme. Initially, primary antibodies against mouse monoclonal NetG1 (D-2, Santa Cruz. Dallas, TX) and rabbit polyclonal NGL-1 (N1N3, Genetex Inc., Irvine, CA) were incubated at 4°C overnight, followed by anti-mouse Q625 and anti-rabbit Q655 secondary antibodies, together with primary rabbit polyclonal Y397P-FAK (Genetex Inc., Irvine, CA) pre-linked to Q605, for an overnight incubation (see Q-dots antibody conjugation details above). After washes, specimens were treated with corresponding monovalent Fab IgG fragments (Jackson Immuno Research Inc., West Grove, PA) to avoid cross-reaction during the incubation of the following cocktail of primary antibodies: mouse monoclonal anti-pan-cytokeratin (clones AE1/AE3, DAKO) to detect the epithelial compartment; rabbit monoclonal anti-vimentin (EPR3776, Abcam) antibodies for mesenchymal (stromal) components. Secondary antibodies used were donkey anti-mouse Cy3 and donkey anti-rabbit Cy2 (Jackson Immuno Research Inc., West Grove, PA). Nuclei were labeled by DRAQ5 (Invitrogen-Thermo Fisher Scientific, Waltham, MA). Lastly, sections were dehydrated in progressive ethanol concentrations and clarified in Toluene before mounting using Cytoseal-60 (Thermo Fisher Scientific, Waltham, MA). Slides were cured overnight in dark at RT before the imaging.

#### Image acquisitions for SMI analysis

Imaging of fluorescently labeled tissue sections was performed utilizing Nuance-FX multispectral imaging system (Caliper LifeSciences, PerkinElmer), using full capabilities of its Tunable Liquid Crystal imaging module, attached to a Nikon Eclipse TE-2000-S epifluorescence microscope. Images were acquired using a Plan Fluor 40X/0.75 objective. A specific wavelength based spectral library for each system was built using control pancreatic tissues (i.e. murine and human), individually labeled for each Qdot and fluorophore used. Unstained specimens were also included to digitally discard specific auto-fluorescence spectra. Each particular spectral library was used for the subsequent image acquisition and analysis. All excitations were achieved via a high-intensity mercury lamp using the following filters for emission-excitation, for Nuance-FX, DAPI (450-720), FITC (500-720), TRITC (580-720), CY5 (680-720). Excitation was conducted from the infrared to ultraviolet spectrum, attempting to avoid energy transfer and excitation of nearby fluorophores. Emission was collected using ‘DAPI’ filter (wavelength range 450–720) for all Qdot labeled markers, while masks used the conventional FITC, TRITC, and CY5 filters.

Collected image cubes (multi-spectral data files) were unmixed to obtain 16-bit (gray scale) individual monochromatic files. Next, the images were processed in bulk for similar levels adjustment and 8-bit monochromatic image conversion for each channel, using Photoshop’s ‘Levels’ and ‘Batch processing’ automated functions. The resulting images were sampled to set identical threshold values for each marker (NetG1, Y397P-FAK and NGL-1) and masks (vimentin and nuclei). Threshold values distinguishing cytokeratin in normal versus tumor samples needed to be independently adjust for each tissue set to assure variations in intensity staining patterns between the two types of epithelium (i.e., normal vs tumoral) were accounted for, as these were needed to be used as “gating masks” identifying epithelial/tumoral area pixels. Processed images were used to analyze distribution, intensity and co-localization of positive-selected pixels using SMIA-CUKIE (SCR_014795) algorithm, available online at https://github.com/cukie/SMIA [38].

#### SMIA-CUKIE usage and outputs

As described previously [38], SMIA-CUKIE algorithm (SCR_014795) was used to analyze high throughput monochromatic images acquired from SMI-processed FFPE tissue sections. In brief, the algorithm is based on the usage of digital masks, constructed from pixel-positive area distribution, resulting in tissue compartmentalization (i.e. cytokeratin positive/vimentin negative area: epithelium/tumor compartment *vs.* vimentin positive/cytokeratin negative area: stromal compartment). Next, we queried mean, median, coverage area, integrated intensity of the markers of interest (i.e. NetG1, NGL-1, Y397P-FAK), under each mask. Each mask and marker required a numeric threshold (0-255), indicating the value of pixel intensity considered positive for each channel (corresponding to each mask and marker). These were kept identical for each marker and mask throughout the study. Finally, the analysis generated an Excel file containing numerical values, which were scrutinized for statistical significance as described in the corresponding statistical analysis section. Black lines through a row indicate an unquantifiable image. The algorithm is available at the public domain: https://github.com/cukie/SMIA_CUKIE. This dataset is available as the supplemental material file: “Human Single Patient SMI”.

### RNA sequencing (RNAseq)

Stranded mRNA-seq library: 1000ng total RNAs from each sample were used to make library according to the product guide of Truseq stranded mRNA library kit (Illumina, Cat# 20020595). In short, mRNAs were enriched twice via poly-T based RNA purification beads, and subjected to fragmentation at 94 degree for 8 minutes via divalent cation method. The 1st strand cDNA was synthesized by Superscript II reverse transcriptase (ThermoFisher, Cat# 18064014) and random primers at 42 degree for 15 minutes, followed by 2nd strand synthesis at 16 degree for 1hour. During second strand synthesis, the dUTP was used to replace dTTP, thereby the second strand was quenched during amplification. A single ‘A’ nucleotide is added to the 3’ ends of the blunt fragments at 37 degree for 30 minutes. Adapters with illuminaP5, P7 sequences as well as indices were ligated to the cDNA fragment at 30 degree for 10 minutes. After SPRIselect beads (Beckman Coulter, Cat# B23318) purification, a 15-cycle PCR reaction was used to enrich the fragments. PCR was set at 98 degree for 10 sec, 60 degree for 30 sec and extended at 72 degree for 30 sec. Libraries were again purified using SPRIselect beads, had a quality check on Agilent 2100 bioanalyzer (serial # DE34903146) using Agilent high sensitivity DNA kit (Cat# 5067-4626), and quantified with Qubit 3.0 flurometer (ThermoFisher Scientific, Cat#Q33216) using Qubit 1x dsDNA HS assay kit (Cat#Q33230). Sample libraries were subsequently pooled and loaded to the Hiseq 2500 (Illumina, serial number SN930). Single end reads at 50bp were generated by using Hiseq rapid SR cluster kit V2 (Illumina, Cat# GD-402-4002) and HiSeq rapid SBS kit v2 50 cycles (Illumina, Cat# FC-402-4022). Faseq files were obtained at Illumina base space (https://basespace.illumina.com) and aligned to Hg38 using Tophat2 [108] and resulting BAM files were used to estimate the gene counts using HT-Seq [109]. Subsequently, to identify differentially expressed genes between CON and NetG1 KO CAFs, we applied DESeq2 [110]. The resulting differentially expressed genes were used as input to GSEA [111] (default parameters with enrichment statistic set as classic). Heatmap depicting differentially expressed genes using mean-centered rlog transformed gene counts were plotted using pheatmap package available in R. Data are organized in tabs (Gene list p<0.001, GSEA list with FDR <0.25, and complete GSEA analysis for positive and negative enrichment), available in Supplemental Table 6.

### Pancreatic Cancer Datasets

We obtained the gene expression data from [42] (processed data available at ArrayExpress ID: E-MTAB-6134), [43] (processed expression data available at Gene Expression Omnibus ID GSE21501) and TCGA (rsem normalized gene expression data available at GDAC Firehose repository for 76 high tumor purity samples). Clinical outcomes data indicating patient overall survival was obtained from the respective data repositories. In the case of Moffitt and TCGA studies, we used Moffitt *et al.* classification to stratify PDAC tumors into basal and classical subtypes. In the case of Puleo *et al.* we applied 5 different classes as described by that study. In order to assess the association between expression of *LRRC4C* and *NTNG1* in PDAC tumors to overall survival (OS), we used expression and clinical data for TCGA and Puleo *et al.* We stratified the expression data into lower (DN) and upper quartiles (UP) for both LRRC4C and NTNG1 and categorized them into 4 different groups (DN-DN; DN-UP; UP-DN and UP-UP). We then estimated the distributions of OS and RFS using Kaplan-Meier methods and were compared using log-rank tests. Analyses to compare the overall survival among the basal and classical subtypes were performed by comparing survival curves with log-rank tests. These were calculated using the R ‘survival’ package Therneau, T. M. & Grambsch, P. M. Modeling Survival Data: Extending the Cox Model (Springer-Verlag, 2010). We applied Gene Set Variation Analysis (GSVA) [112] with mx.diff parameter set to true on all the 76 PDAC tumor cases in TCGA using 12,913 gene sets available from MSigDB [113]. To classify NGL-1 into low- and high-expressors, we selected cases on Z-scores (85th CI for mean). Wilcoxon-rank sum test was applied to identify significantly enriched MSigDB datasets (p<0.005). Select enriched pathways that are significantly different between low- and high-expressors of NGL-1 were plotted as waterfall plots with low- (green color) and high-expressors (red). Cases that did not satisfy the Z-score cut-off were plotted as gray bars.

### KM Plots Generated with KM Plotter

To generate KM Plots correlating with overall survival (OS) and relapse free survival (RFS) data with *NTNG1* and *LRRC4C* mRNA expression from the TCGA dataset, we used the online tool KM Plotter (kmplot.com; [114]). We selected the pancreatic adenocarcinoma dataset (N=177) from the Pan-Cancer RNAseq section of the website for overall survival. For the RFS analysis, we selected the RFS dataset, for which there was data available for 69 patients [115]. We used the default auto cutoff algorithm to establish the high and low expressors of *NTNG1* and *LRRC4C* (cutoff = 22) combined. The false discovery rate (FDR) for the cutoff values for *NTNG1* and *LRRC4C* were >0.5. KM plots were generated and Hazard Ratios (HR) and p-values (log rank test) were obtained.

### Statistics

For all *in vitro* and animal experiments, Prism 7.0 (Graph Pad Software, San Diego, California) was used for all statistical analysis. For comparison between two groups, a two tailed Student’s T-Test was performed. For comparisons between more than two groups, a One-Way ANOVA was performed, using either a Dunnett’s multiple comparisons test (compared to control condition) or Tukey’s multiple comparisons test (comparing all conditions to one another), depending on the experiment. Groups were deemed statistically significantly different from one another if the p-value was less than or equal to 0.05. On graphs, significance was defined as follows: * p<0.05; ** p<0.01; *** < 0.001; **** p< 0.0001. For the correlation of the SMI analysis, linear regression was applied to the SMIA generated values of two markers at the designated components, and the R^2^ value was calculated. All *in vitro* experiments were performed at least three times (unless explicitly stated in the figure legend) and the orthotopic injections were performed in two cohorts each time, with the following resultant sample sizes:

-Human co-injection SCID model: N=6 for all co-injections; N=4 and 5 for CON or NGL-1 KO PDAC cells alone; N= 3 and 4 for CON or NetG1 KO CAFs alone.
-Murine cancer cell model: N=7 for Control, and N=19 for combined NGL-1 KO injected mice.
-Murine immunocompromised mouse models: N=5 for all groups
-Murine co-injection models: N= 5 for all PDAC/fibroblast co-injected groups, N=4 and 3 for CON and NGL-1 KO cancer cell alone, N= 2 for all fibroblast groups (CON, NetG1 KD1+2, GFP, NetG1 WT, NetG1 mut) injected alone.
-mAb model: N=9 for IgG group; N =10 for mAb group.

## Supporting information

Key Resources Table

## Author Contributions

Conceptualization, E.C., R.F., D.B.V.C.; Methodology, E.C., R.F., J.F.B, I.A., L.G., J.W, T.P., K.S.C., A.M., A.N.L., S.G, K.Q.C., R.T., S.B., M.G.V.H., H.H., E.N., N.S., T.L, S.P., R.T., Y.T.; Validation, T.L., R.F., D.B.V.C, J.W., A.M. D.Ro., D.Re., Y.T.; Formal Analysis, R.F., D.B.V.C, A.M., A.N.L., J.F.B., J.W., T.L., N.S., T.P., K.D., E.M., K.Q.C., E.C., D.Ro., Y.Z., S.P., A.J.K., H.W.; Investigation, R.F., D.B.V.C., J.F.B., T.L., N.S., T.P., S.G., L.G., A.M., A.N.L., E.M., H.H.; Resources, E.C., I.A., M.V.H., S.B., W.D.K., W.S.E.D., K.C., H.W.; Data Curation; J.F.B., N.S., D.K., Y.Z., D.Ro., T.L, S.P., Y.T.; Writing-original draft, R.F., D.B.V.C.; Writing-review & editing, E.C., I.A., J.W., J.F.B., T.P., S.B., K.S.C., A.M., R.F., D.B.V.C., S.P., M.G.V.H; Visualization, E.C., J.F.B., R.F., D.B.V.C., J.W., A.M., A.N.L., T.P., D.Ro., T.L.; Supervision, E.C., I.A., K.S.C.; Funding Acquisition, E.C., I.A., K.S.C., M.G.V.H.

## Acknowledgments

This study is dedicated to the memories of the late Dr. Patricia Keely (pioneer in the field of ECM biology and an amazing mentor) and Neelima Shah (our amazing co-author on this study, with over 30+ years in research and the most beautiful soul), who continue to inspire our work. We would like to thank Dr. Martin Humphries for the SNAKA 51 antibody. We appreciate the advice from Dr. Alexander Macfarlane on NK cells and flow cytometry. We thank Jim Oesterling for his great feedback and technical assistance with flow cytometry. We greatly appreciate the NetG1/NGL-1 discussions with Dr. Shigeyoshi Itohara. We would also like to acknowledge Dr. Stephen M. Sykes, Dr. Daniela Di Marcantonio, and Esteban Martinez for their time and expertise with RT-qPCR. We are grateful for the kind gifts provided by Robert Weinberg (pBABE-neo-hTERT vector; Addgene plasmid # 1774), Dr. Feng Zhang (LentiCRISPRv2; Addgene plasmid # 52961), Dr. Didier Trono (psPAX2; Addgene plasmid # 12260), Charles Gersbach (pLV hU6-sgRNA hUbC-dCas9-KRAB-T2a-Puro; Addgene plasmid #71236), and Dr. Alexey Ivanov (pLV-CMV-H4-puro vector). This work was supported in part by gifts donated to the memory of Judy Costin, funds from the Martin and Concetta Greenberg Pancreatic cancer Institute, Pennsylvania’s DOH Health Research Formula Funds, the Greenfield Foundation, the 5th AHEPA Cancer Research Foundation, Inc., Fox Chase In Vino Vita Institutional Pilot Award, as well as NIH/NCI grants R01CA113451-10 (EC), T32CA009035 (RF), R01CA113451-09S1 (JFB/EC), R21 CA231252 (EC/IA), R01CA188430 (IA), R03CA212949 (IA), R01GM116911 (IA), R01CA194263 (KC) Core grant CA06927 in support to Fox Chase Cancer Center’s facilities including: Bio Sample Repository, Light Microscopy, Small Animal Imaging, Biostatistics and Bioinformatics, Flow Cytometry, Laboratory Animal, Cell Culture, DNA sequencing, Genotyping and Real-time PCR, Histopathology, Immune Monitoring Facility, and Talbot Library. DOD grant WX81XWH-15-1-0170 (EC/RF/JFB). The authors also acknowledge support from F32CA213810 (AM), the Damon Runyon Cancer Research Foundation (ANL), R01CA168653, R01CA201276, and P30CA1405141 (M.G.V.H), the Lustgarten Foundation (MGVH), the MIT Center for Precision Medicine (MGVH), SU2C (MGVH), and the Ludwig Center at MIT (MGVH). MGVH is a Howard Hughes Medical Institute Faculty Scholar.

## Supplemental Figure Legends

**Figure S1.**
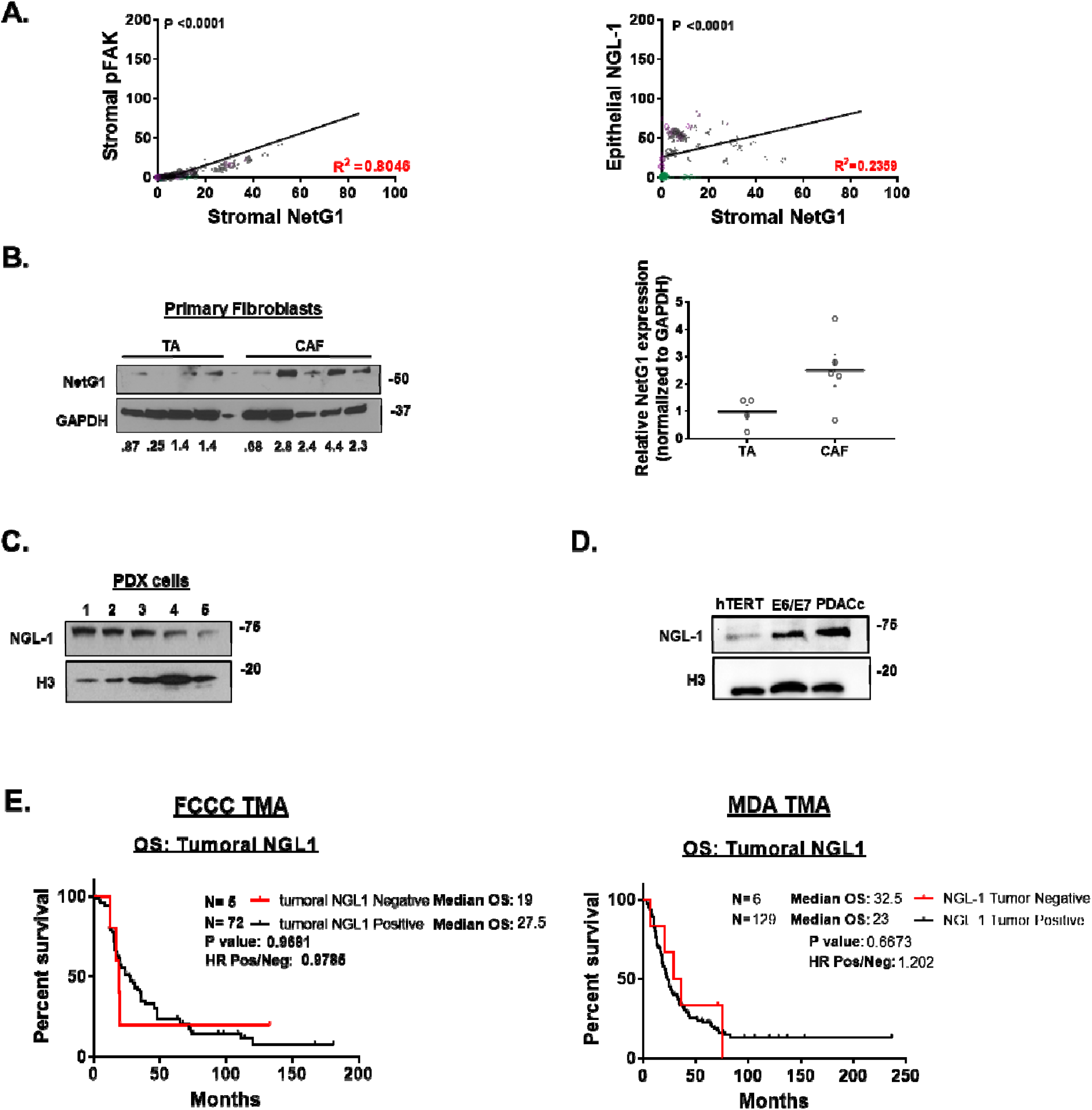
*Stromal* NetG1 expression correlates with stromal pFAK and tumoral NGL-1 expression. **A. (LEFT)** Linear regression analysis comparing expression of stromal pFAK with stromal NetG1 and **(RIGHT)** epithelial NGL-1 with stromal NetG1. Each dot represents the integrated intensity of each parameter from a single image; normal (green dots), tumor adjacent (purple), and PDAC tissue (gray). P-values are shown in black and R^2^ values are shown in red. **B.** Representative western blot comparing NetG1 expression in 4 TA and 5 CAFs obtained from PDAC patient tissues, and subsequently passaged 1 time in 3D to obtain lysates, with the respective quantifications normalized to GAPDH and mean TA expression. **C.** Representative western blot comparing NGL-1 expression in cell lines derived from PDAC PDX models. H3 was used as a loading control. **D.** Representative western blot comparing NGL-1 expression in an isogenic cell line of human pancreatic epithelial cells (HPNE system) with progressive mutations, simulating different stages of pancreatic cancer. hTERT = immortalized epithelia; E6/E7 = Rb and P53 inactivated hTERT cells; PDACc = E6/E7 cells with KRAS^G12D^. H3 was used as a loading control. **E.** Kaplan-Meier plots showing overall patient survival over time, based on NGL-1 IHC score in tumor cells. **LEFT:** FCCC TMA. **RIGHT:** MDA TMA.

**Figure S2.**
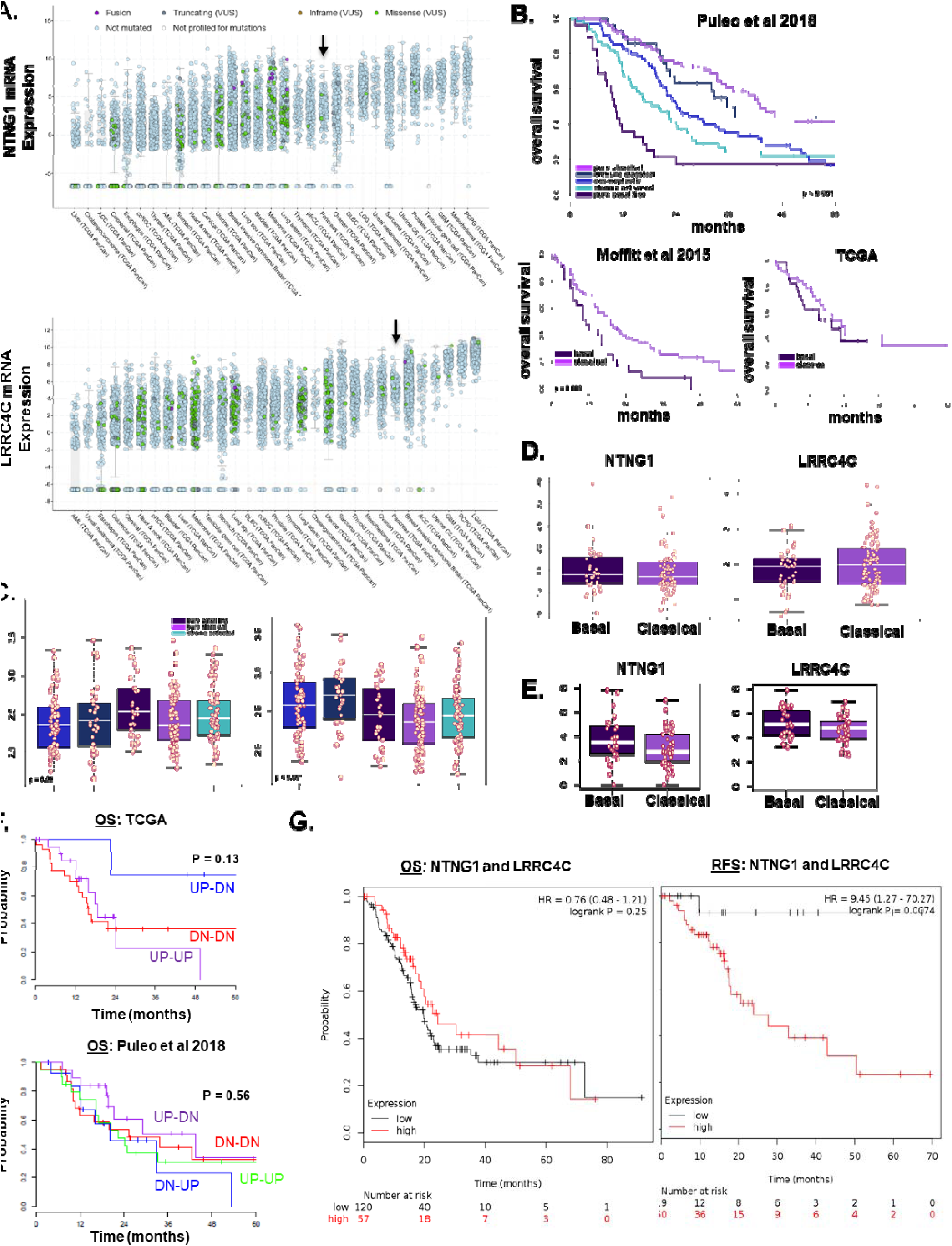
Expression of NetG1 and its ligand NGL-1 in cancers. **A.** Plots show the mRNA expression of NetG1 (*NTNG1*) and its ligand NGL-1 (*LRRC4C*) in various cancers. The expression data (Z-scores of RNASeq V2 RSEM expression data) are shows as box plots where each dot represents a patient with colors describing the molecular alterations within that patient. Pancreatic cancer is highlighted with an arrow. **B.** Kaplan-Meier curves for overall survival based on 5 different subtypes of PDAC cases as described by Puleo *et al.* (**LEFT**) (**MIDDLE)** Moffitt *et al.* and TCGA (**RIGHT**) are shown. **C-E.** Box plots show the expression of *NTNG1* and *LRRC4C* in PDAC tumors for various subtypes across Puleo *et al.* **(C)**, Moffitt *et al*. (**D**) and TCGA (**E**) studies. Significance of differences in expression of these two genes across subtypes is estimated using Kruskal-Wallis test. **F.** Kaplan-Meier curves depicting overall survival (OS) versus expression of *NTNG1* and *LRRC4C* in the PDAC TCGA (**LEFT**) and Puleo *et al*. (**RIGHT**) datasets. Data were categorized into four groups based on *NTNG1* and *LRRC4C* mRNA expression (lower and upper quartiles): DN-DN (*LRRC4C* low, *NTNG1* low), DN-UP (*LRRC4C* low, *NTNG1* high), UP-DN (*LRRC4C* high, *NTNG1* low) and UP-UP (*LRRC4C* high, *NTNG1* high). Note that no instances of *LRRC4C* high/*NTNG1* low expression (UP-DN) could be found in the TCGA dataset. **G.** Kaplan-Meier plots depicting OS (**LEFT**) and RFS (relapse free survival) (**RIGHT**) correlated with the combined expression of *NTNG1* and *LRRC4C* generated from the TCGA dataset using the online tool KM Plotter. Log rank test was used for all graphs to determine statistical significance of OS/RFS associated with gene expression.

**Figure S3.**
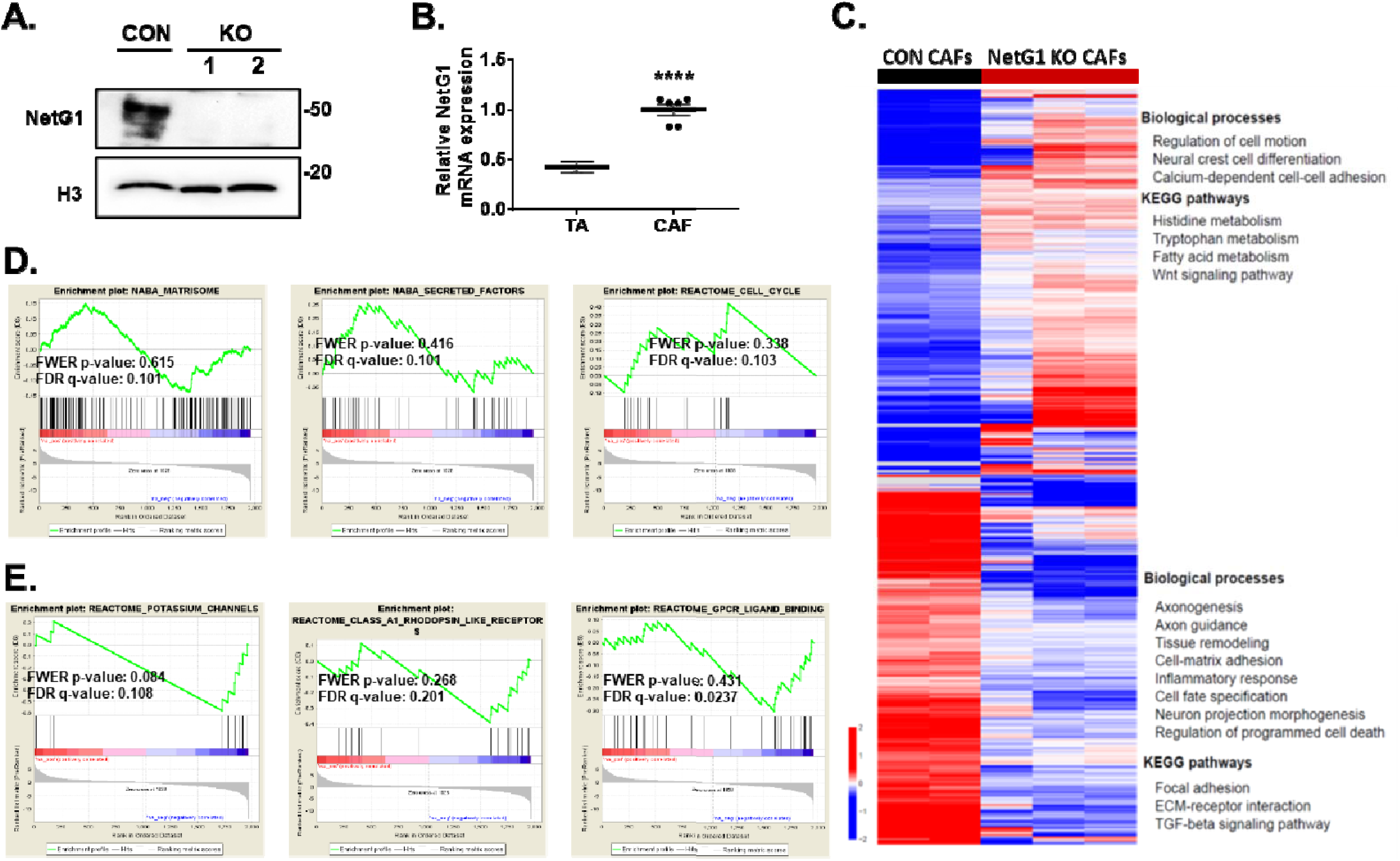
*Genetic* ablation of NetG1 results in a more “normalized” fibroblastic gene expression profile. **A.** Generation of two human NetG1 KO CAF lines, using CRISPR/Cas9, as shown in the western blot. H3 was used as a loading control. Cell line containing vector targeting GFP was used as control**. B.** Representative mRNA expression of NetG1 in TA and CAFs from a single patient cultured in 3D. RNAseq analysis comparing CON CAFs to NetG1 KO CAFs revealed a normalization of CAF function upon NetG1 deletion. **C.** Heatmap showing differentially expressed genes (p < 0.001) in NetG1 KO CAFs with genes increased (red) and decreased (blue) expression levels. Over-represented biological process or a pathway among these genes are shown. **D and E** show GSEA enrichment plots for positive and negative enrichment respectively in NetG1 KO CAFs.

**Figure S4.**
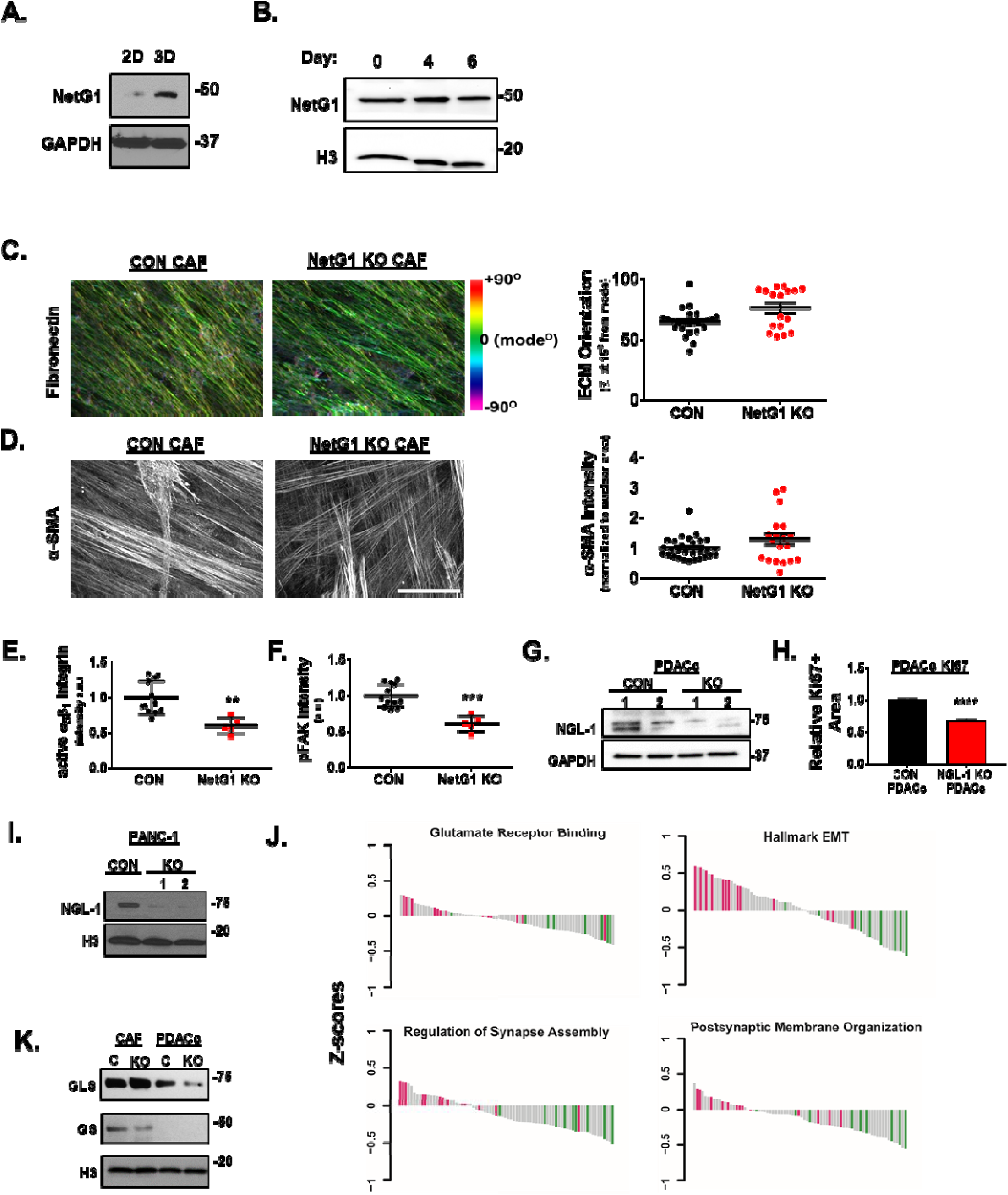
CAFs maintains canonical myofibroblastic features while decreasing traits associated with tumor promoting properties of CAFs. **A.** Representative western blot demonstrating NetG1 expression in 2D vs. 3D culturing conditions. Note the significantly higher expression of NetG1 in 3D. GAPDH served as a loading control. **B.** Representative western blot of NetG1 illustrating consistent expression of the protein throughout 6 days of 3D ECM production. H3 was used as a loading control. **C.** ECM fiber orientation, as indicated by fibronectin IF and pseudo-colored to highlight fiber orientation using the OrientationJ plugin in ImageJ. Color bar: color coded angle orientation showing green color used for normalized mode angle fibers. FN fibers within 15 degrees of the mode angle were scored as an indicator of ECM orientation and were graphed on the right. **D. (LEFT)** Representative images of α-smooth muscle actin (α-SMA) IF. **(RIGHT)** Quantification (intensity) displayed in the graph. Scale bar is 50 µm and serves for both (**C**) and (**D**). **E.** Quantification of active α_5_β_1_-integrin; IF intensity, detected using the SNAKA51 antibody in CON or NetG1 KO CAFs. **F.** Quantification of phospho-FAK^397^ (pFAK) IF intensity in CON or NetG1 KO CAFs. **G.** Validation of CRISPR/Cas9 mediated NGL-1 KO in PDACc cells, as shown in the representative western blot. GAPDH was used as a loading control. Cell lines containing vector targeting GFP were used as control. **H.** Quantification of the relative Ki67 staining in CON and NGL-1 KO PDACc grown in 3D, in CON CAF generated ECMs. Student’s T-test, ** p< 0.01; *** p< 0.001; **** p< 0.0001. **I.** Validation of CRISPR/Cas9 mediated KO of NGL-1 in PANC-1 cells, as shown in the representative western blot. H3 was used as a loading control. Cell lines containing vector targeting GFP were used as control. **J.** Select enriched pathways that are significantly different between cases with low- and high-expression of NGL-1 were plotted as waterfall plots with low- (green color) and high-expressors (red). Cases that did not satisfy the Z-score cut-off were plotted as gray bars. **K.** Representative western blots of GLS and GS in control CAFs (C) and NetG1 KO CAFs (KO) in comparison to CON and NGL-1 KO PDACc. H3 was used as a loading control.

**Figure S5.**
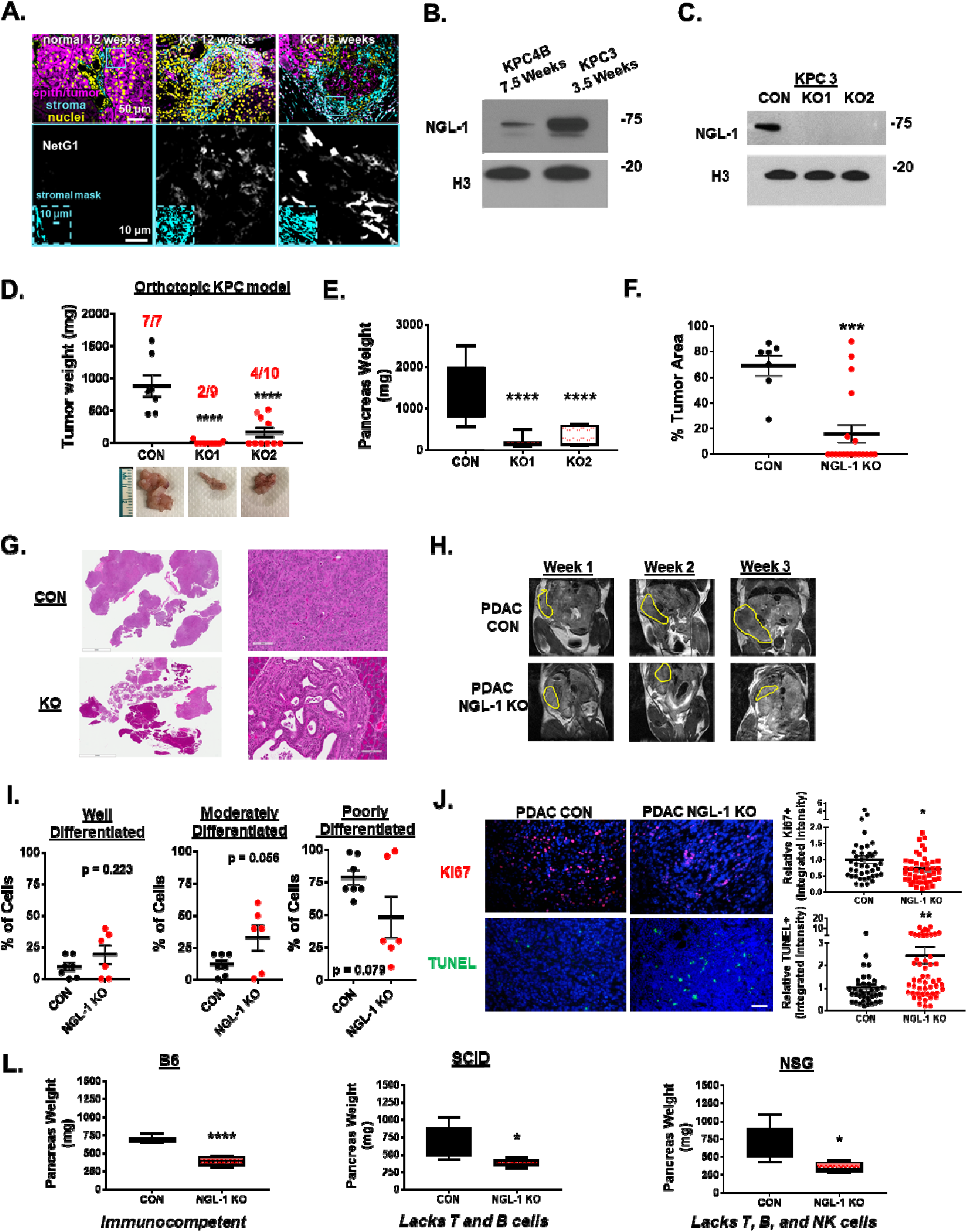
NGL-1 KO in murine PDAC cells stunts tumorigenesis in an orthotopic murine model. **A.** SMI was conducted on murine pancreatic tissue from wild type C57BL/B6 (normal) or KC mice (12 weeks or 16 weeks old). **(TOP Panel)** Epithelium (purple), stroma (cyan), and nuclei (yellow) were labeled and used as “masks” to query stromal (i.e., cyan^+^/purple^-^ pixel areas shown in **BOTTOM Insert** and marked by cyan square) expression of NetG1 (white, **BOTTOM Panel**). Representative scale bars are shown**. B.** Representative western blot of NGL-1 in two different murine PDAC cell line clones derived from KPC mice (KPC4B and KPC3). H3 was used as a loading control. Note the time for orthotopically injected tumor progression in weeks listed above the lanes. **C.** Western blot of NGL-1, demonstrating knockout in two independent clones of KPC3 murine PDAC cells. H3 was used as a loading control. **D.** C57BL/6 mice were injected orthotopically with CON or NGL-1 KO KPC3 cells (10^6^) and were sacrificed 3.5 weeks later. Graph depicting quantification tumor weight from each mouse group. Red numbers above the bars represent tumor incidence over total mice for each group. Images below display representative pancreata from each group. **E.** Graphs depicting the weight of the pancreata isolated from each group of mice. **F.** Quantification of the % area that tumor cells occupy in the pancreata of mice injected with CON KPC3 or NGL-1 KO KPC3 cells (pooled KO mice together). **G.** Representative images of the entire pancreas displaying tumor area coverage. Scale bar is 5 mm**. H.** Representative MRI images for pancreata injected with CON or NGL-1 KO cells, taken at 1, 2, and 3 weeks after injection. Yellow outline marks the pancreas in each image. **I.** Quantification of the differentiation status of tumors within the pancreatic tissue that formed tumors (CON, N = 7; KO, N = 6), as measured by % cells that were classified as well differentiated, moderately differentiated, or poorly differentiated by a blinded pathologist. **J. (LEFT)** Representative images of Ki67 (red) and TUNEL (green) staining of tumors that developed from mice injected with CON KPC3 or NGL-1 KO KPC3 cells. Nuclei is stained by DAPI (blue). Scale bar is 50 µm**. (RIGHT)** Quantification of Ki67 or TUNEL staining (integrated intensity) normalized to nuclei, relative to the control PDAC group. Each dot represents the quantification of Ki67 or TUNNEL image from a single image. N= 7 CON and 6 pooled KO tumors. **L**. B6, SCID (lacks T and B cells), or NSG (lacks T, B, and NK cells) mice were injected with 10^6^ CON KPC3 or NGL-1 KO #2 cells. Animals were sacrificed after 3.5 weeks and pancreas weight (mg) was assessed. (n=5, for each cell type, in each background). One-Way ANOVA, Dunnett’s multiple comparison test (**D-E**) or Student’s T-Test (**F**, **I, J, L**). *p<0.05, **p<0.01, ***p<.001, ****p<0.0001.

**Figure S6.**
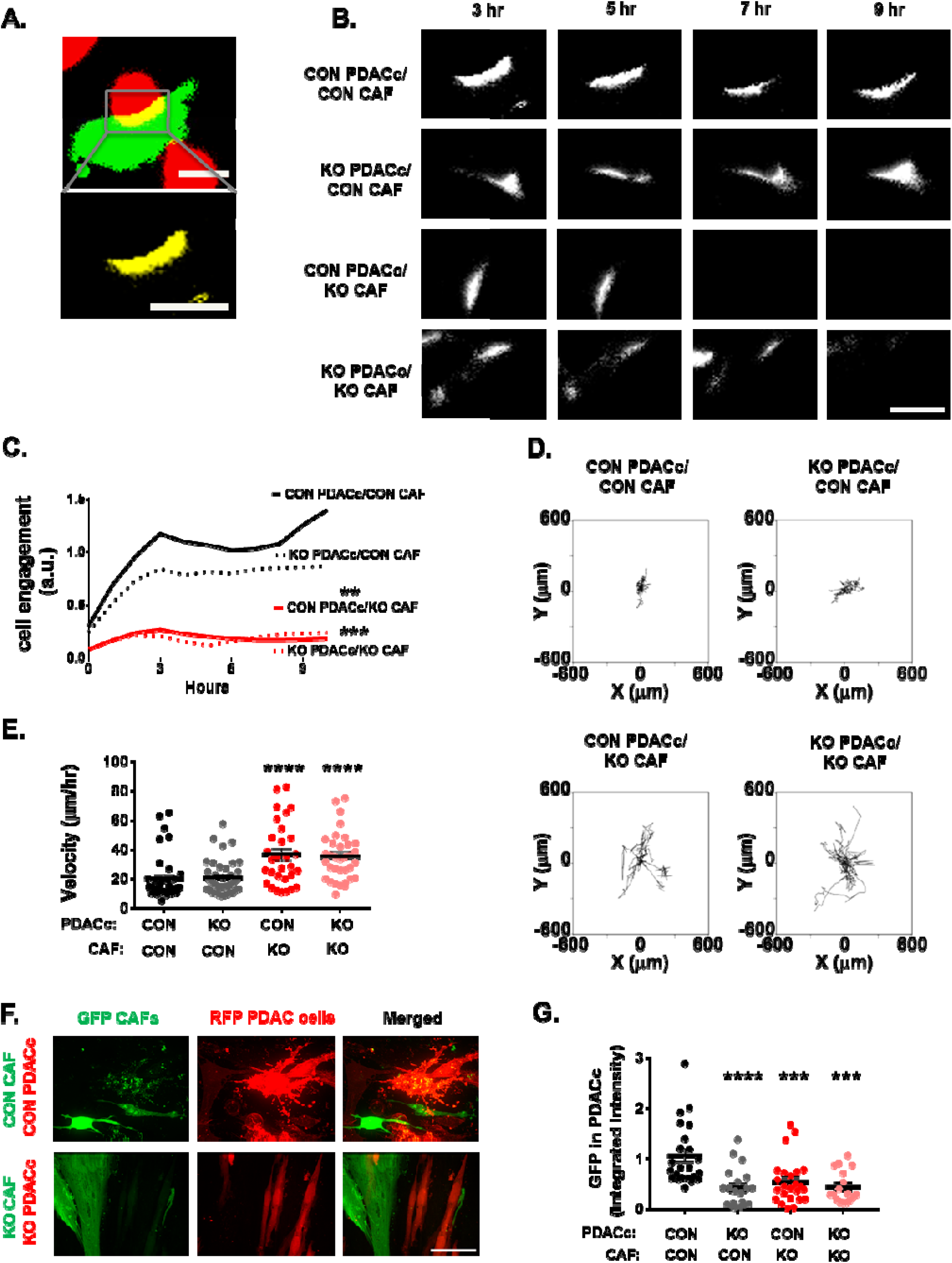
NetG1 promotes heterotypic cell interactions and material transfer from CAFs to PDAC cells. A cell engagement assay was undertaken by co-culturing 2×10^4^ CON or NGL-1 KO PDACc (RFP^+^) in 3D with 2×10^4^ CON or NetG1 KO CAFs (GFP^+^) for 24 hours, and areas of cell engagement were measured. **A.** Example of the type of areas analyzed in the cell engagement assay. Note the yellow area in the top panel (representing the engagement area), with the zoomed in region of the yellow in the panel below. Scale bar is 10 µm. **B.** Representative images of the areas of cell engagement for each co-culture condition, over the time span of the assay. Scale bar is 10 µm. **C.** Quantification of cell engagement areas during 10 hours of engagement. **D. 3D** Cell movement, following cells invading through CAF generated ECMs was performed by co-culturing RFP^+^ CON or NGL-1 KO PDACc (2×10^4^) with GFP^+^ CON or NetG1 KO CAFs (2×10^4^) for 24 hours; time lapse videos measured PDACc cell movement over that timespan. Shown are all motility tracks of PDACc, moving through ECMs, that were acquired from each co-culture experimental condition. **E.** Velocity (µm/hour) of PDACc from (**D**). * compared to CON PDACc/ CON CAF group. **F.** Representative images from a 36 hour 3D co-culture of GFP labeled CON or NetG1 KO CAFs (2×10^4^) with RFP labeled CON or NGL-1 KO PDACc (2×10^4^), in SF media. Note the yellow color inside of RFP^+^ PDACc in the CON/CON condition, indicative of PDACc intracellular GFP, acquired from CAFs (Merged). Scale bar is 50 µm**. G.** Quantification of the amount of GFP^+^ PDACc cells, calculated as integrated GFP fluorescence intensity, from the 36 hour material transfer assay in (**F**), relative to the CON PDACc /CON CAF co-culture condition. Each dot represents the quantification of one image. N = 3 independent experiments with a minimum of 7 images per condition. Note how loss of stromal NetG1 or tumoral NGL-1 sufficed to prevent CAF material transfer into PDAC cells. One-way ANOVA, Dunnett’s multiple comparison test ** p< 0.01; *** p< 0.001; **** p< 0.0001.

**Figure S7.**
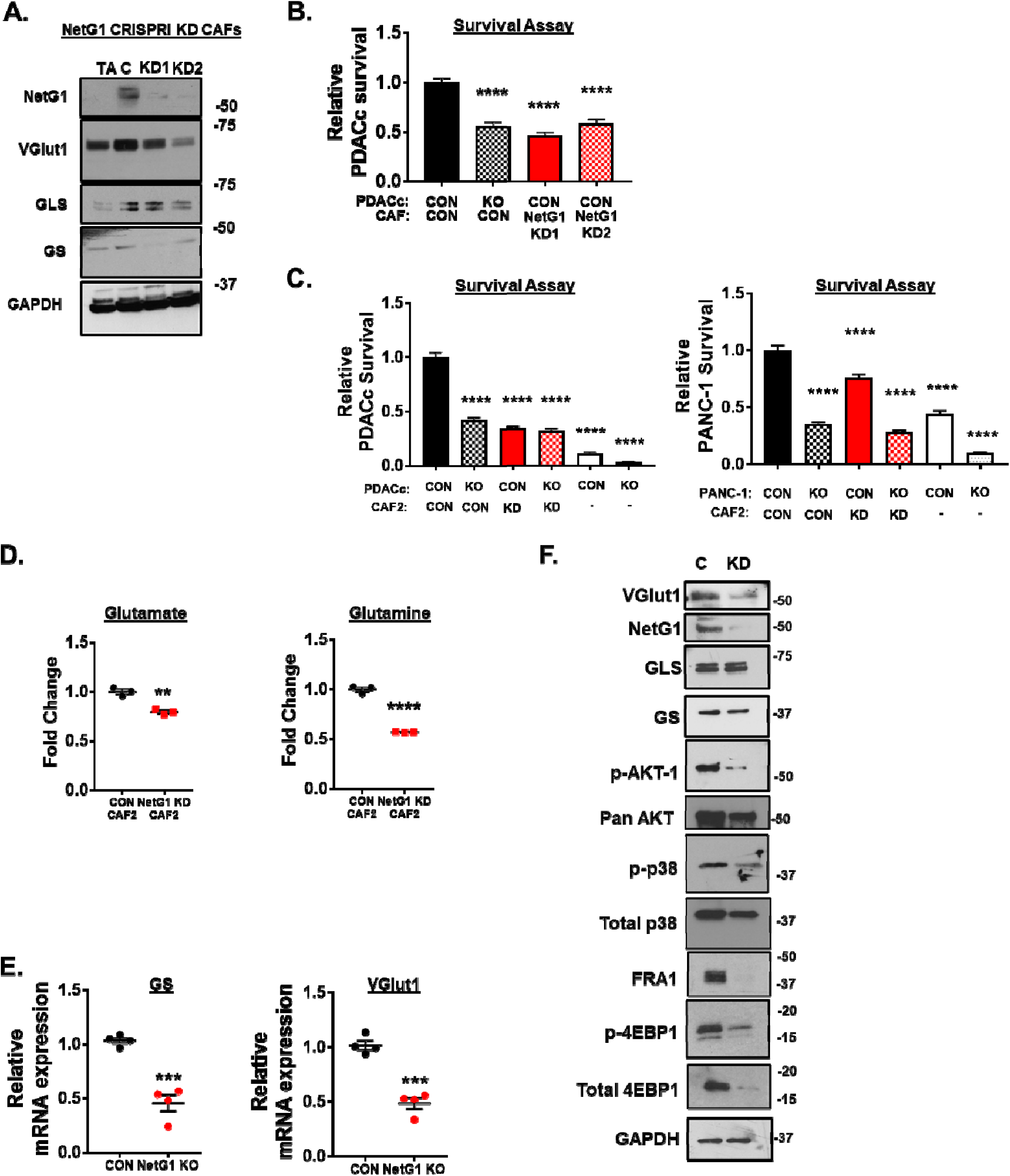
NetG1 KD CAFs, generated with CRISPRi, and a second CAF line phenocopy the metabolic functions of NetG1 KO CAFs. **A.** Representative western blots displaying the expression levels of NetG1, VGlut1, GLS, GS in TA, CON CAFs, and NetG1 KD CAFs (KD1 and KD2). Note the effective knockdown of NetG1, as well as downregulation of VGlut1 and GS expression. **B.** PDAC cell survival was determined after RFP^+^ CON or NGL-1 KO PDACc (2×10^4^) were cultured with CON or NetG1 KD CAFs for 96 hours in SF/Gln free media. **C.** RFP^+^ CON or NGL-1 KO PDACc or PANC-1 cells (2×10^4^) were co-cultured in 3D with CON or NetG1 KD CAF2 (2×10^4^) or alone in the absence of serum and Gln for 4 days followed by cell survival assessments. * compared to CON PDAC/CON CAF2. One-way ANOVA, Dunnett’s multiple comparison test, ****p<.0001. **D.** Quantification of the relative amounts of Glu and Gln in the conditioned media of CON or NetG1 KD CAF2, after 48 hours of 3D culture in serum/Gln free media. **E.** Graphs depicting relative mRNA expression of GS and VGlut1 genes in CON and NetG1 KO CAFs, normalized to CON expression levels. *compared to CON CAFs. Student’s T-Test, **p<0.01, ***p<0.001, ****p<0.0001. **F.** Representative western blots of VGlut1, NetG1, GLS, GS, p-AKT, AKT, p-p38, p38, FRA1, p-4E-BP1, and 4E-BP1 in CON or NetG1 KD CAF2. GAPDH was used as a loading control.

**Figure S8.**
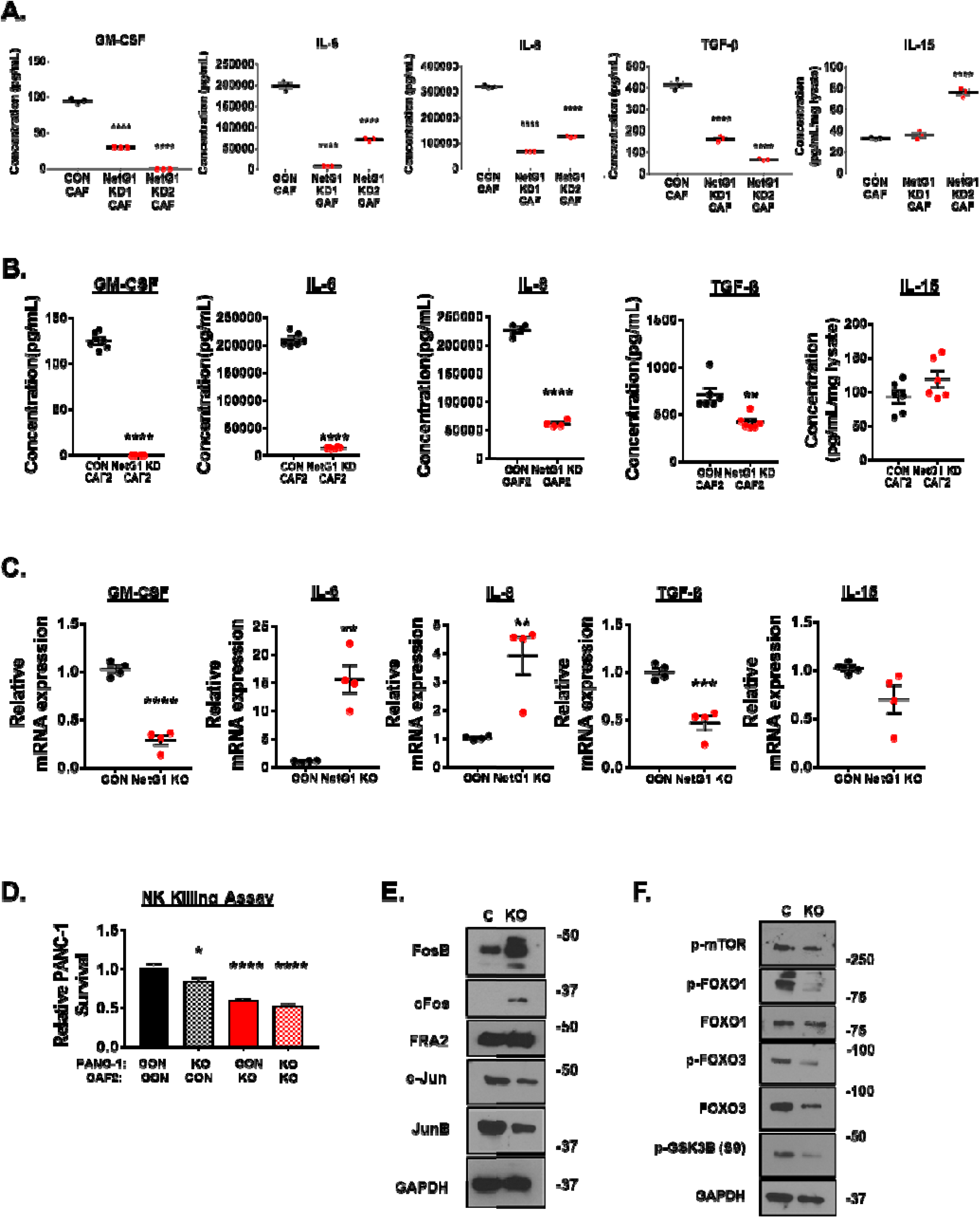
Knockdown of NetG1 by CRISPRi in two patient CAF lines results in a loss of pro-tumor immunosuppressive capacities. **A.** CON CAFs or NetG1 KD CAFs were allowed to produce CM for 48 hours in 3D. The CM was then subjected to ELISA analysis for the quantification of the following cytokines: GM-CSF, IL-6, IL-8, TGF-β, and IL-15. N= 3 biological replicates. One-way ANOVA, Dunnett’s multiple comparison test, * p< 0.05; ** p< 0.01; *** p< 0.001; **** p< 0.0001. **B.** CON or NetG1 KD CAF2 were allowed to produce CM for 48 hours in 3D. The CM was then subjected to ELISA analysis for the quantification of the following cytokines: GM-CSF, IL-6, IL-8, TGF-β, and IL-15. N= 6 biological replicates. Student’s T-Test, ** p<.01, *** p<.001, **** p<.0001. **C.** Graphs depicting relative mRNA expression GM-CSF, IL-6, IL-8, TGF-β, and IL-15 genes in CON and NetG1 KO CAFs, normalized to CON expression levels. *compared to CON CAFs. Student’s T-Test, **p<0.01, ***p<0.001, ****p<0.0001. **D.** RFP^+^ CON or NGL-1 KO PANC-1 cells (2×10^4^) were co-cultured in 3D with CON or NetG1 KD CAFs (2×10^4^) in the presence of active NK cells (8×10^4^) for 48 hours and relative PANC-1 survival was measured. * compared to CON PANC-1/CON CAF2. One-way ANOVA, Dunnett’s multiple comparison test. * p< 0.05; ** p< 0.01; *** p< 0.001; **** p< 0.0001. **E**. Representative blots of various AP-1 transcription factor subunits, downstream to p38 signaling, comparing expression in CON and NetG1 KO CAFs. **F.** Representative blots of downstream factors of AKT signaling, comparing expression in CON and NetG1 KO CAFs. GAPDH is used as a loading control in (**E**) and (**F**).

**Figure S9.**
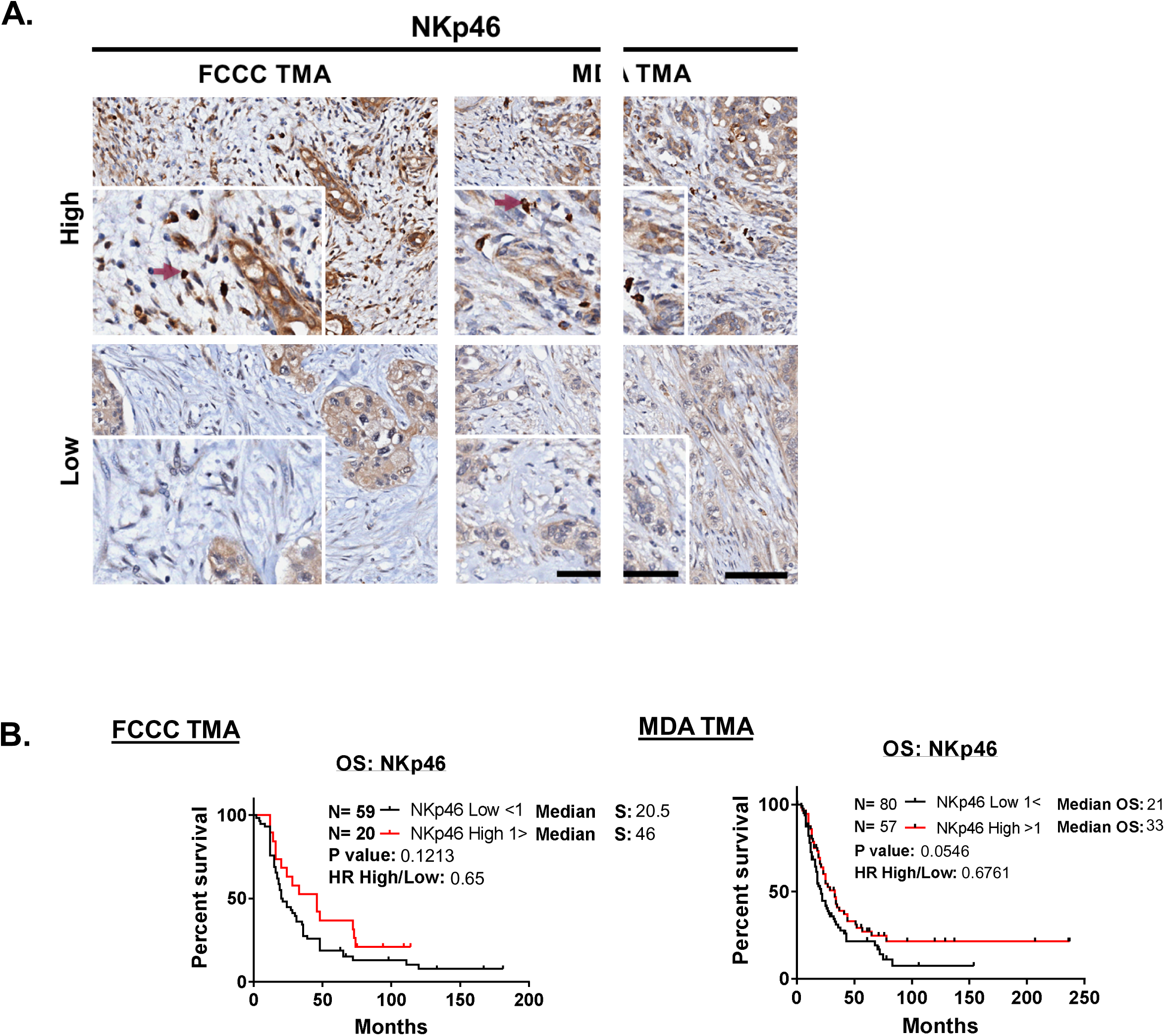
NK cell infiltration into the tumor trends towards directly correlating with patient overall survival. NKp46, a marker of NK cells, was stained immunohistochemically in both TMAs and blindly scored by a pathologist (0-4 scale; see methods for details). **A.** Representative images from both TMAs demonstrating patients that presented with a high or low amount of NKp46^+^ cell infiltration. Inserts magnify regions to display NKp46 positive cells (dark red arrows). Scale bars represent 100 μm. Note that there was a high degree of background staining of epithelial cells, and this was not considered in the scoring of the staining. **B.** Kaplan-Meier plots displaying overall survival as a function of NKp46 immunohistochemical score. **LEFT:** FCCC TMA; **RIGHT:** MDA TMA. Log-rank test was used to determine statistical significance between the groups.

**Figure S10.**
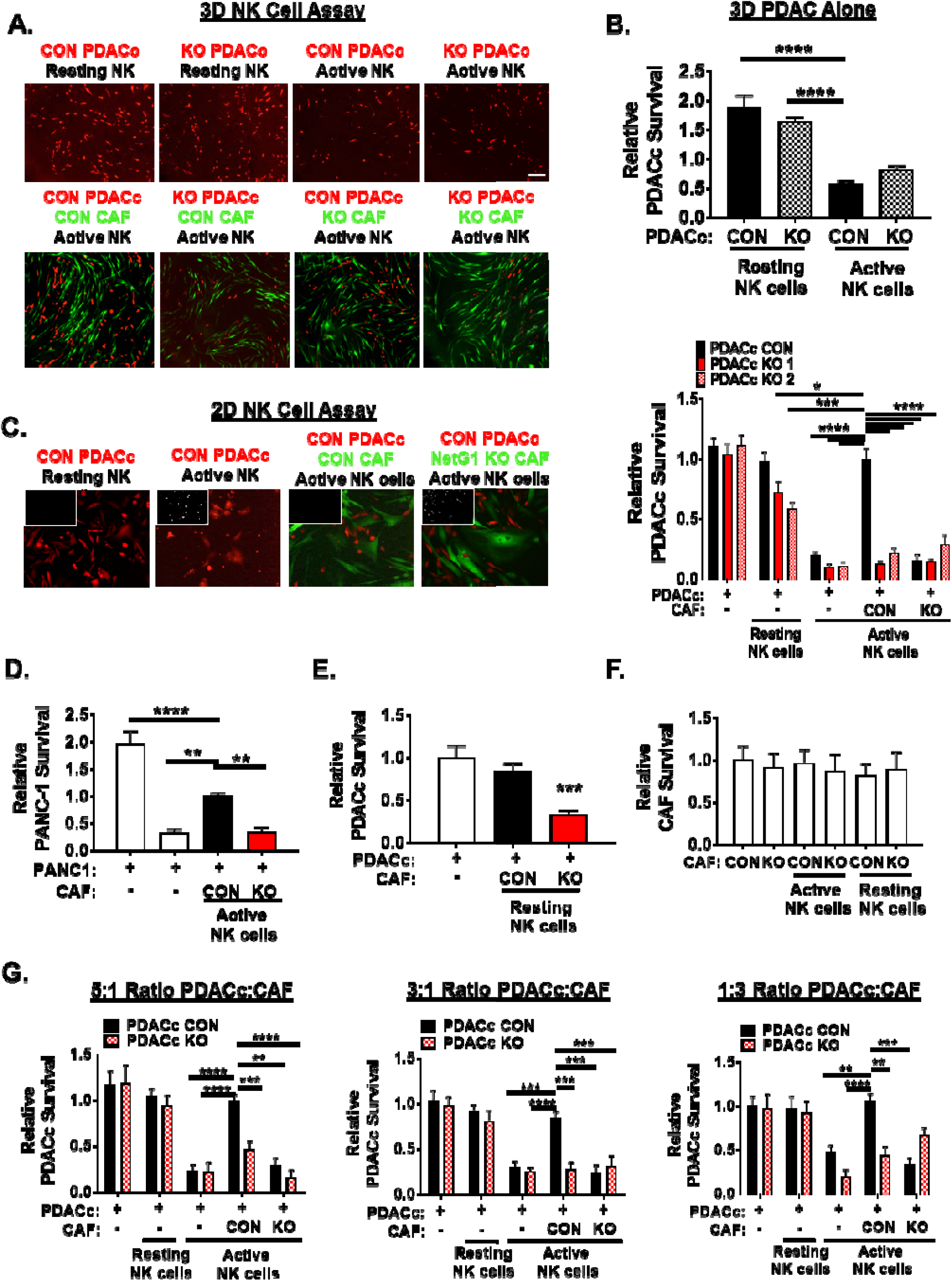
NetG1 KO CAFs support NK cell killing of PDAC cells. **A.** Representative images of PDAC cell survival. RFP^+^ CON or NGL-1 KO PDACc (2×10^4^) were co-cultured in 3D in the absence **(TOP panel)** or presence **(BOTTOM panel)** of GFP^+^ CON or NetG1 KO CAFs (2×10^4^), together with active or resting NK-92 cells (8×10^4^) for 48 hours. Scale bar is 500 µm**. B.** Quantification of **(A); BOTTOM** panel. * compared to CON with active NK cells. **C. (LEFT)** 2D NK cell assay: RFP^+^ CON or NGL-1 KO PDACc were co-cultured 1:1 (2×10^4^ cells) with resting or activated NK-92 cells (8×10^4^) alone, or with GFP^+^ CON and NetG1 KO CAFs for 8 hours. PDACc cell survival was assessed. Representative images of PDACc (red) and CAFs (green) from the various assay conditions. White insert in the top left-hand corner shows dead cell channel, marked by Sytox blue. **(RIGHT)** Quantification of the 2D NK cell killing assay. * compared to CON PDACc/CON CAF cultured with active NK-92 cells. **D.** The same NK cell assay performed in (**C**), but with PANC-1 cells instead of PDACc cells. * compared to CON PANC-1/CON CAF cultured with active NK-92 cells. **E**. Quantification of the co-culture conditions with resting NK cells from (**C**). * compared to CON PDACc/CON CAF cultured with resting NK-92 cells. **F.** Quantification of CAF control conditions from the experiment in (**C**). **G.** The NK cell killing assay was performed as in (**C**), but with the indicated ratios of PDACc:CAF (5:1, 3:1, 1:3). * compared to CON PDACc/CON CAF cultured with active NK-92 cells, for each ratio. Graphs show mean survival and standard errors of the three independent experiments. One-way ANOVA, Dunnett’s multiple comparison test, * p< 0.05; ** p< 0.01; *** p< 0.001; **** p< 0.0001.

**Figure S11.**
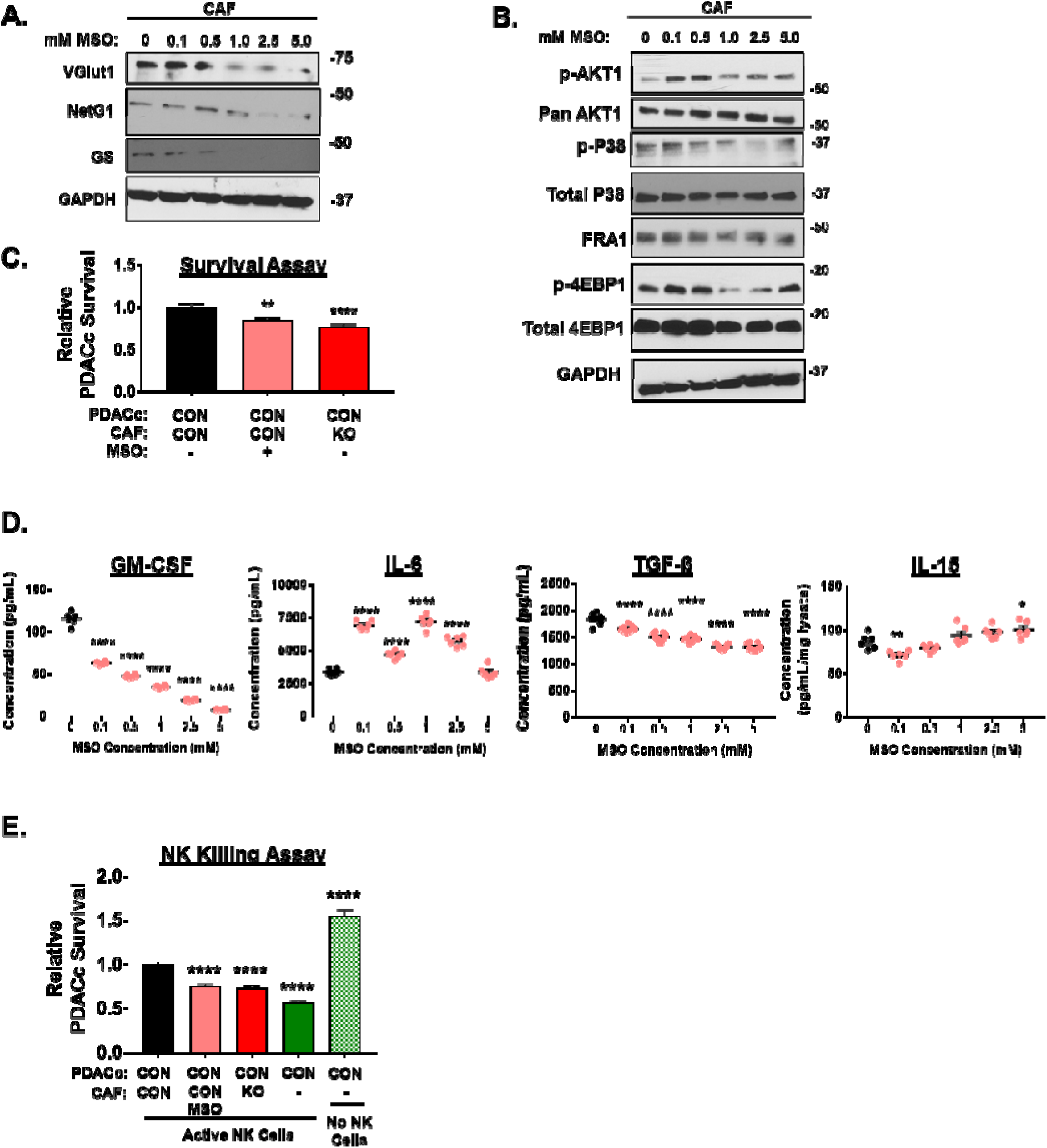
Pharmacological inhibition of GS in CAFs by MSO causes a decrease in CAF mediated support to PDAC cells. **A.** Representative western blots demonstrating changes in VGlut1, NetG1, GLS, and GS protein expression in CAFs treated for 24 hours in 3D with increasing concentrations of MSO (GS inhibitor). For all blots, GAPDH served as the loading control. **B.** Representative blots of p-AKT, AKT, p-p38, p38, FRA1, p-4E-BP1, and 4E-BP1 in vehicle or MSO treated CAFs. GAPDH was used as a loading control. **C.** RFP^+^ CON or NGL-1 PDAC (2×10^4^) were cultured, using 3D conditions as before, for 96 hours in serum/Gln free media with CON or NetG1 KO CAFs (2×10^4^) that were pre-treated 24 hours with vehicle (water) a GS inhibitor (1 mM methionine sulfoximine) and PDAC survival was evaluated. **D.** GS inhibited CAFs (MSO, 0, 0.1, 0.5, 1, 2.5, and 5 mM) were grown in 3D and allowed to generate CM for 48 hours in SF/Gln free media. Graphs show quantification of ELISAs of the indicated cytokines. * compared to CAFs treated with 0 mM MSO. **E.** CON RFP^+^ PDAC cells (2×10^4^) were co-cultured with vehicle (water) or MSO (1 mM) 24 hour pre-treated CON or NetG1 KO CAFs (2×10^4^), in the presence of active NK-92 cells (IL-2 preactivated), and PDACc cell survival was assessed 48 hours later. PDACc alone with and without active NK cells were used as controls. * compared to CON PDACc/CON CAF with NK cells. One-Way ANOVA, Dunnett’s multiple comparison test. *p<0.05, **p<0.01, ****p<0.0001.

**S12.**
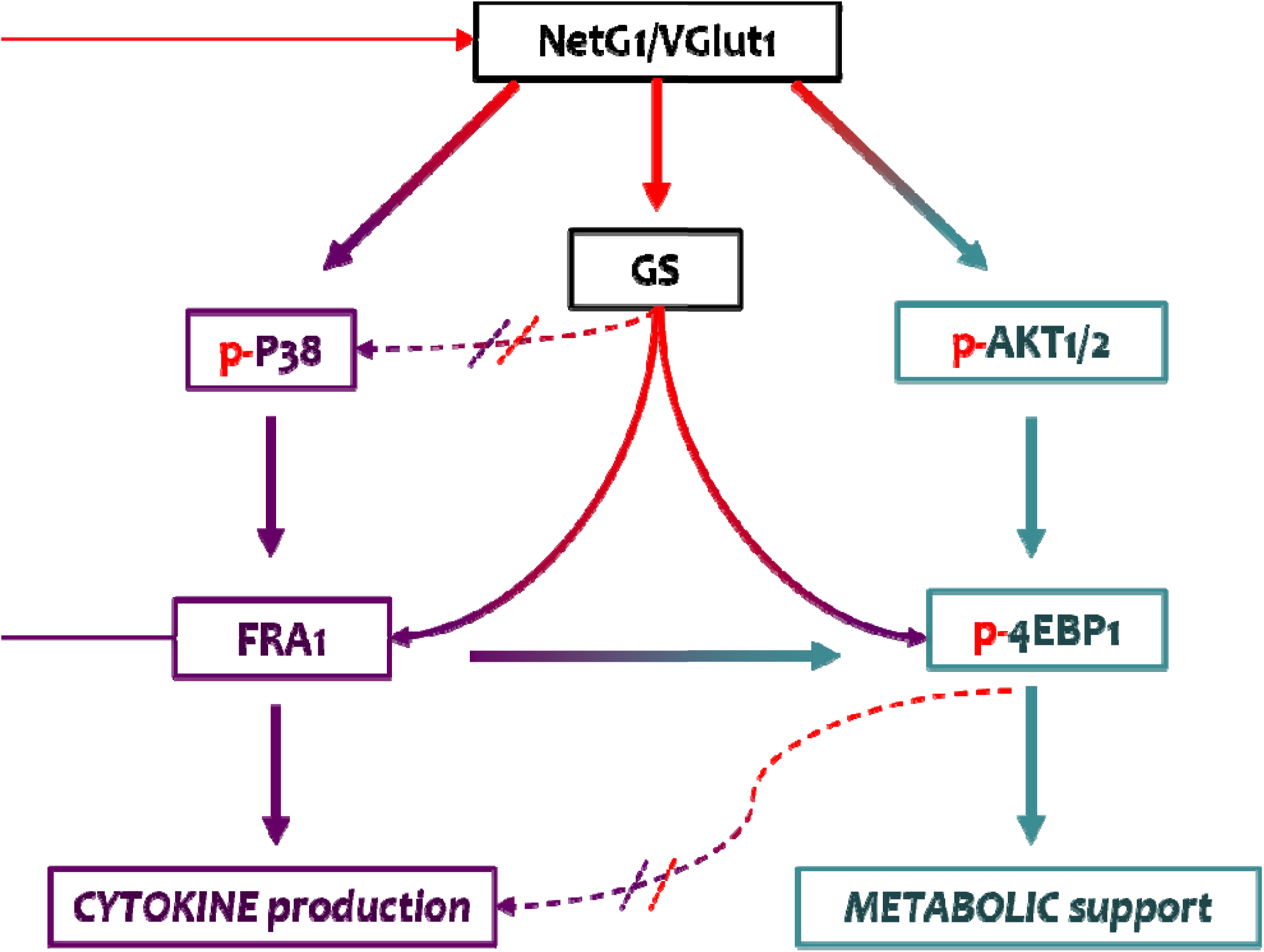
NetG1 Signaling Circuit in CAFs. There are two major arms of the signaling circuit regulated by NetG1, the cytokine arm and the metabolic arm. NetG1 and VGlut1 sit atop the signaling network, and increase levels of GS, p-p38, and p-AKT (red arrows). GS, p38, and AKT are further downstream, and they regulate FRA1 and 4E-BP1 levels, which begin to diverge in what CAF functions they control. FRA-1 regulates cytokine production in CAFs, as well as mediates partial effects on CAF generated metabolism (dotted arrow), through Gln production. 4E-BP1 has an inhibitory effect on cytokine production (blue lines), while directly controlling CAF generated Glu/Gln. There is also positive feedback to the top of the circuit (curved arrow), and crosstalk to the AKT/4E-BP1 arm, all originating at FRA1. Thus, NetG1 controls pro-tumor CAF functions, through a complex signaling network.

## References

1. Rahib, L., et al., Projecting cancer incidence and deaths to 2030: the unexpected burden of thyroid, liver, and pancreas cancers in the United States. Cancer Res, 2014. 74(11): p. 2913–21.

2. Siegel, R.L., K.D. Miller, and A. Jemal, Cancer statistics, 2020. CA Cancer J Clin, 2020. 70(1): p. 7–30.

3. Ryan, D.P., T.S. Hong, and N. Bardeesy, Pancreatic adenocarcinoma. N Engl J Med, 2014. 371(22): p. 2140–1.

4. Hidalgo, M., Pancreatic cancer. N Engl J Med, 2010. 362(17): p. 1605–17.

5. Sahai, E., et al., A framework for advancing our understanding of cancer-associated fibroblasts. Nat Rev Cancer, 2020. 20(3): p. 174–186.

6. Olive, K.P., et al., Inhibition of Hedgehog signaling enhances delivery of chemotherapy in a mouse model of pancreatic cancer. Science, 2009. 324(5933): p. 1457–61.

7. McCarroll, J.A., et al., Role of pancreatic stellate cells in chemoresistance in pancreatic cancer. Front Physiol, 2014. 5: p. 141.

8. Chan, T.S., et al., Metronomic chemotherapy prevents therapy-induced stromal activation and induction of tumor-initiating cells. J Exp Med, 2016. 213(13): p. 2967–2988.

9. Hwang, R.F., et al., Cancer-associated stromal fibroblasts promote pancreatic tumor progression. Cancer Res, 2008. 68(3): p. 918–26.

10. Djurec, M., et al., Saa3 is a key mediator of the protumorigenic properties of cancer-associated fibroblasts in pancreatic tumors. Proc Natl Acad Sci U S A, 2018. 115(6): p. E1147–E1156.

11. Erkan, M., et al., The activated stroma index is a novel and independent prognostic marker in pancreatic ductal adenocarcinoma. Clin Gastroenterol Hepatol, 2008. 6(10): p. 1155–61.

12. Özdemir BC, P.-H.T., Carstens JL, Zheng X, Wu CC, Simpson TR, Laklai H, Sugimoto H, Kahlert C, Novitskiy SV, De Jesus-Acosta A, Sharma P, Heidari P, Mahmood U, Chin L, Moses HL, Weaver VM, Maitra A, Allison JP, LeBleu VS, Kalluri R., Depletion of carcinoma-associated fibroblasts and fibrosis induces immunosuppression and accelerates pancreas cancer with reduced survival. Cancer Cell, 2014. 25(6): p. 719–34.

13. Rhim, A.D., et al., Stromal elements act to restrain, rather than support, pancreatic ductal adenocarcinoma. Cancer Cell, 2014. 25(6): p. 735–47.

14. Kim, E.J., et al., Pilot clinical trial of hedgehog pathway inhibitor GDC-0449 (vismodegib) in combination with gemcitabine in patients with metastatic pancreatic adenocarcinoma. Clin Cancer Res, 2014. 20(23): p. 5937–45.

15. Provenzano, P.P., et al., Enzymatic targeting of the stroma ablates physical barriers to treatment of pancreatic ductal adenocarcinoma. Cancer Cell, 2012. 21(3): p. 418–29.

16. Kamphorst, J.J., et al., Human pancreatic cancer tumors are nutrient poor and tumor cells actively scavenge extracellular protein. Cancer Res, 2015. 75(3): p. 544–53.

17. Erkan, M., M. Kurtoglu, and J. Kleeff, The role of hypoxia in pancreatic cancer: a potential therapeutic target? Expert Rev Gastroenterol Hepatol, 2016. 10(3): p. 301–16.

18. Son, J., et al., Glutamine supports pancreatic cancer growth through a KRAS-regulated metabolic pathway. Nature, 2013. 496(7443): p. 101–5.

19. Jin, L., G.N. Alesi, and S. Kang, Glutaminolysis as a target for cancer therapy. Oncogene, 2016. 35(28): p. 3619–25.

20. Seo, J.W., et al., Autophagy is required for PDAC glutamine metabolism. Sci Rep, 2016. 6: p. 37594.

21. Lyssiotis, C.A. and A.C. Kimmelman, Metabolic Interactions in the Tumor Microenvironment. Trends Cell Biol, 2017. 27(11): p. 863–875.

22. von Ahrens, D., et al., The role of stromal cancer-associated fibroblasts in pancreatic cancer. J Hematol Oncol, 2017. 10(1): p. 76.

23. Basso, D., E. Gnatta, and M. Plebani, Pancreatic Cancer Fostered Immunosuppression Privileges Tumor Growth and Progression. Journal of Clinical & Cellular Immunology, 2014(5): p. 16.

24. Clark, C.E., et al., Dynamics of the immune reaction to pancreatic cancer from inception to invasion. Cancer Res, 2007. 67(19): p. 9518–27.

25. Fu, Y., et al., The critical roles of activated stellate cells-mediated paracrine signaling, metabolism and onco-immunology in pancreatic ductal adenocarcinoma. Mol Cancer, 2018. 17(1): p. 62.

26. Harper, J. and R.C. Sainson, Regulation of the anti-tumour immune response by cancer-associated fibroblasts. Semin Cancer Biol, 2014. 25: p. 69–77.

27. Wu, Q., et al., Functions of pancreatic stellate cell-derived soluble factors in the microenvironment of pancreatic ductal carcinoma. Oncotarget, 2017. 8(60): p. 102721–102738.

28. Ene-Obong, A., et al., Activated pancreatic stellate cells sequester CD8+ T cells to reduce their infiltration of the juxtatumoral compartment of pancreatic ductal adenocarcinoma. Gastroenterology, 2013. 145(5): p. 1121–32.

29. Kumar, V., et al., Cancer-Associated Fibroblasts Neutralize the Anti-tumor Effect of CSF1 Receptor Blockade by Inducing PMN-MDSC Infiltration of Tumors. Cancer Cell, 2017. 32(5): p. 654–668 e5.

30. Buck, M.D., et al., Metabolic Instruction of Immunity. Cell, 2017. 169(4): p. 570–586.

31. Kouidhi, S., F. Ben Ayed, and A. Benammar Elgaaied, Targeting Tumor Metabolism: A New Challenge to Improve Immunotherapy. Front Immunol, 2018. 9: p. 353.

32. Renner, K., et al., Metabolic Hallmarks of Tumor and Immune Cells in the Tumor Microenvironment. Front Immunol, 2017. 8: p. 248.

33. Gardiner, C.M. and D.K. Finlay, What Fuels Natural Killers? Metabolism and NK Cell Responses. Front Immunol, 2017. 8: p. 367.

34. Nakashiba, T., et al., Netrin-G1: a novel glycosyl phosphatidylinositol-linked mammalian netrin that is functionally divergent from classical netrins. J Neurosci, 2000. 20(17): p. 6540–50.

35. Song, Y.S., et al., Trans-induced cis interaction in the tripartite NGL-1, netrin-G1 and LAR adhesion complex promotes development of excitatory synapses. J Cell Sci, 2013. 126(Pt 21): p. 4926–38.

36. Lin, J.C., et al., The netrin-G1 ligand NGL-1 promotes the outgrowth of thalamocortical axons. Nat Neurosci, 2003. 6(12): p. 1270–6.

37. Amatangelo, M.D., et al., Stroma-derived three-dimensional matrices are necessary and sufficient to promote desmoplastic differentiation of normal fibroblasts. Am J Pathol, 2005. 167(2): p. 475–88.

38. Franco-Barraza, J., et al., Matrix-regulated integrin alphavbeta5 maintains alpha5beta1-dependent desmoplastic traits prognostic of neoplastic recurrence. Elife, 2017. 6.

39. Campbell, P.M., et al., K-Ras promotes growth transformation and invasion of immortalized human pancreatic cells by Raf and phosphatidylinositol 3-kinase signaling. Cancer Res, 2007. 67(5): p. 2098–106.

40. Cerami, E., et al., The cBio cancer genomics portal: an open platform for exploring multidimensional cancer genomics data. Cancer Discov, 2012. 2(5): p. 401–4.

41. Gao, J., et al., Integrative analysis of complex cancer genomics and clinical profiles using the cBioPortal. Sci Signal, 2013. 6(269): p. pl1.

42. Puleo, F., et al., Stratification of Pancreatic Ductal Adenocarcinomas Based on Tumor and Microenvironment Features. Gastroenterology, 2018. 155(6): p. 1999–2013 e3.

43. Moffitt, R.A., et al., Virtual microdissection identifies distinct tumor- and stroma-specific subtypes of pancreatic ductal adenocarcinoma. Nat Genet, 2015. 47(10): p. 1168–78.

44. Cancer Genome Atlas Research Network. Electronic address, a.a.d.h.e. and N. Cancer Genome Atlas Research, Integrated Genomic Characterization of Pancreatic Ductal Adenocarcinoma. Cancer Cell, 2017. 32(2): p. 185–203 e13.

45. Cukierman, E., et al., Taking cell-matrix adhesions to the third dimension. Science, 2001. 294(5547): p. 1708–12.

46. Hingorani, S.R., et al., Preinvasive and invasive ductal pancreatic cancer and its early detection in the mouse. Cancer Cell, 2003. 4(6): p. 437–50.

47. Hingorani, S.R., et al., Trp53R172H and KrasG12D cooperate to promote chromosomal instability and widely metastatic pancreatic ductal adenocarcinoma in mice. Cancer Cell, 2005. 7(5): p. 469–83.

48. Muranen, T., et al., Starved epithelial cells uptake extracellular matrix for survival. Nat Commun, 2017. 8: p. 13989.

49. Sousa, C.M., et al., Pancreatic stellate cells support tumour metabolism through autophagic alanine secretion. Nature, 2016. 536(7617): p. 479–83.

50. Zhao, H., et al., Tumor microenvironment derived exosomes pleiotropically modulate cancer cell metabolism. Elife, 2016. 5: p. e10250.

51. Xiang, Y., et al., Targeted inhibition of tumor-specific glutaminase diminishes cell-autonomous tumorigenesis. J Clin Invest, 2015. 125(6): p. 2293–306.

52. Yang, L., et al., Targeting Stromal Glutamine Synthetase in Tumors Disrupts Tumor Microenvironment-Regulated Cancer Cell Growth. Cell Metab, 2016. 24(5): p. 685–700.

53. Liguz-Lecznar, M. and J. Skangiel-Kramska, Vesicular glutamate transporters (VGLUTs): the three musketeers of glutamatergic system. Acta Neurobiol Exp (Wars), 2007. 67(3): p. 207–18.

54. Woodcock, H.V., et al., The mTORC1/4E-BP1 axis represents a critical signaling node during fibrogenesis. Nat Commun, 2019. 10(1): p. 6.

55. Rajasekaran, S., et al., Expression profiling of genes regulated by Fra-1/AP-1 transcription factor during bleomycin-induced pulmonary fibrosis. BMC Genomics, 2013. 14: p. 381.

56. Matsukawa, H., et al., Netrin-G/NGL complexes encode functional synaptic diversification. J Neurosci, 2014. 34(47): p. 15779–92.

57. Zhang, Q., et al., Netrin-G1 regulates fear-like and anxiety-like behaviors in dissociable neural circuits. Sci Rep, 2016. 6: p. 28750.

58. Eastwood, S.L. and P.J. Harrison, Decreased mRNA expression of netrin-G1 and netrin-G2 in the temporal lobe in schizophrenia and bipolar disorder. Neuropsychopharmacology, 2008. 33(4): p. 933–45.

59. Nectoux, J., et al., Netrin G1 mutations are an uncommon cause of atypical Rett syndrome with or without epilepsy. Pediatr Neurol, 2007. 37(4): p. 270–4.

60. Hao, W., et al., The pan-cancer landscape of netrin family reveals potential oncogenic biomarkers. Sci Rep, 2020. 10(1): p. 5224.

61. Sho, S., et al., A prognostic mutation panel for predicting cancer recurrence in stages II and III colorectal cancer. J Surg Oncol, 2017. 116(8): p. 996–1004.

62. Andrew, A.S., et al., Hyper-Methylated Loci Persisting from Sessile Serrated Polyps to Serrated Cancers. Int J Mol Sci, 2017. 18(3).

63. Castro-Bello, F., et al., High serum glutamic acid levels in patients with carcinoma of the pancreas. Digestion, 1976. 14(4): p. 360–3.

64. Roux, C., et al., Endogenous glutamine decrease is associated with pancreatic cancer progression. Oncotarget, 2017. 8(56): p. 95361–95376.

65. Biancur, D.E., et al., Compensatory metabolic networks in pancreatic cancers upon perturbation of glutamine metabolism. Nat Commun, 2017. 8: p. 15965.

66. Muir, A., et al., Environmental cystine drives glutamine anaplerosis and sensitizes cancer cells to glutaminase inhibition. Elife, 2017. 6.

67. Sullivan, M.R., et al., Quantification of microenvironmental metabolites in murine cancers reveals determinants of tumor nutrient availability. Elife, 2019. 8.

68. Li, L. and D. Hanahan, Hijacking the neuronal NMDAR signaling circuit to promote tumor growth and invasion. Cell, 2013. 153(1): p. 86–100.

69. Popoli, M., et al., The stressed synapse: the impact of stress and glucocorticoids on glutamate transmission. Nat Rev Neurosci, 2011. 13(1): p. 22–37.

70. Chalmers, Z.R., et al., Analysis of 100,000 human cancer genomes reveals the landscape of tumor mutational burden. Genome Med, 2017. 9(1): p. 34.

71. Evans, R.A., et al., Lack of immunoediting in murine pancreatic cancer reversed with neoantigen. JCI Insight, 2016. 1(14).

72. Yarchoan, M., A. Hopkins, and E.M. Jaffee, Tumor Mutational Burden and Response Rate to PD-1 Inhibition. N Engl J Med, 2017. 377(25): p. 2500–2501.

73. Van Audenaerde, J.R.M., et al., Natural Killer cells and their therapeutic role in pancreatic cancer: A systematic review. Pharmacol Ther, 2018.

74. Vaheri, A., et al., Nemosis, a novel way of fibroblast activation, in inflammation and cancer. Exp Cell Res, 2009. 315(10): p. 1633–8.

75. Heneberg, P., Paracrine tumor signaling induces transdifferentiation of surrounding fibroblasts. Crit Rev Oncol Hematol, 2016. 97: p. 303–11.

76. Van Audenaerde, J.R.M., et al., Interleukin-15 stimulates natural killer cell-mediated killing of both human pancreatic cancer and stellate cells. Oncotarget, 2017. 8(34): p. 56968–56979.

77. Trotta, R., et al., TGF-beta utilizes SMAD3 to inhibit CD16-mediated IFN-gamma production and antibody-dependent cellular cytotoxicity in human NK cells. J Immunol, 2008. 181(6): p. 3784–92.

78. Takeuchi, S., et al., Chemotherapy-Derived Inflammatory Responses Accelerate the Formation of Immunosuppressive Myeloid Cells in the Tissue Microenvironment of Human Pancreatic Cancer. Cancer Res, 2015. 75(13): p. 2629–40.

79. Galic, M.A., K. Riazi, and Q.J. Pittman, Cytokines and brain excitability. Front Neuroendocrinol, 2012. 33(1): p. 116–25.

80. Haroon, E., A.H. Miller, and G. Sanacora, Inflammation, Glutamate, and Glia: A Trio of Trouble in Mood Disorders. Neuropsychopharmacology, 2017. 42(1): p. 193–215.

81. Vesce, S., et al., Glutamate release from astrocytes in physiological conditions and in neurodegenerative disorders characterized by neuroinflammation. Int Rev Neurobiol, 2007. 82: p. 57–71.

82. Biankin, A.V., et al., Pancreatic cancer genomes reveal aberrations in axon guidance pathway genes. Nature, 2012. 491(7424): p. 399–405.

83. Demir, I.E., H. Friess, and G.O. Ceyhan, Neural plasticity in pancreatitis and pancreatic cancer. Nat Rev Gastroenterol Hepatol, 2015. 12(11): p. 649–59.

84. Long, Y., et al., 2-Deoxy-D-Glucose Exhibits Anti-seizure Effects by Mediating the Netrin-G1-KATP Signaling Pathway in Epilepsy. Neurochem Res, 2019. 44(4): p. 994–1004.

85. Brichkina, A., et al., p38MAPK builds a hyaluronan cancer niche to drive lung tumorigenesis. Genes Dev, 2016. 30(23): p. 2623–2636.

86. Alspach, E., et al., p38MAPK plays a crucial role in stromal-mediated tumorigenesis. Cancer Discov, 2014. 4(6): p. 716–29.

87. Yamamura, Y., et al., Akt-Girdin signaling in cancer-associated fibroblasts contributes to tumor progression. Cancer Res, 2015. 75(5): p. 813–23.

88. Jiang, X., et al., Expression and function of FRA1 protein in tumors. Mol Biol Rep, 2020. 47(1): p. 737–752.

89. Atsaves, V., et al., AP-1 Transcription Factors as Regulators of Immune Responses in Cancer. Cancers (Basel), 2019. 11(7).

90. Bhagat, T.D., et al., Lactate-mediated epigenetic reprogramming regulates formation of human pancreatic cancer-associated fibroblasts. Elife, 2019. 8.

91. Busch, S., et al., Cellular organization and molecular differentiation model of breast cancer-associated fibroblasts. Mol Cancer, 2017. 16(1): p. 73.

92. Qin, X., B. Jiang, and Y. Zhang, 4E-BP1, a multifactor regulated multifunctional protein. Cell Cycle, 2016. 15(6): p. 781–6.

93. Duluc, C., et al., Pharmacological targeting of the protein synthesis mTOR/4E-BP1 pathway in cancer-associated fibroblasts abrogates pancreatic tumour chemoresistance. EMBO Mol Med, 2015. 7(6): p. 735–53.

94. Ohlund, D., E. Elyada, and D. Tuveson, Fibroblast heterogeneity in the cancer wound. J Exp Med, 2014. 211(8): p. 1503–23.

95. Costa, A., et al., Fibroblast Heterogeneity and Immunosuppressive Environment in Human Breast Cancer. Cancer Cell, 2018. 33(3): p. 463–479 e10.

96. Costea, D.E., et al., Identification of two distinct carcinoma-associated fibroblast subtypes with differential tumor-promoting abilities in oral squamous cell carcinoma. Cancer Res, 2013. 73(13): p. 3888–901.

97. Ohlund, D., et al., Distinct populations of inflammatory fibroblasts and myofibroblasts in pancreatic cancer. J Exp Med, 2017. 214(3): p. 579–596.

98. Jiang, H., et al., Targeting focal adhesion kinase renders pancreatic cancers responsive to checkpoint immunotherapy. Nat Med, 2016. 22(8): p. 851–60.

99. Franco-Barraza, J., et al., Preparation of Extracellular Matrices Produced by Cultured and Primary Fibroblasts. Curr Protoc Cell Biol, 2016. 71: p. 10 9 1–10 9 34.

100. Beglyarova, N., et al., Screening of Conditionally Reprogrammed Patient-Derived Carcinoma Cells Identifies ERCC3-MYC Interactions as a Target in Pancreatic Cancer. Clin Cancer Res, 2016. 22(24): p. 6153–6163.

101. Ritchie, M.E., et al., limma powers differential expression analyses for RNA-sequencing and microarray studies. Nucleic Acids Res, 2015. 43(7): p. e47.

102. Merico, D., et al., Enrichment map: a network-based method for gene-set enrichment visualization and interpretation. PLoS One, 2010. 5(11): p. e13984.

103. Wang, X., et al., PrimerBank: a PCR primer database for quantitative gene expression analysis, 2012 update. Nucleic Acids Res, 2012. 40(Database issue): p. D1144–9.

104. Rezakhaniha, R., et al., Experimental investigation of collagen waviness and orientation in the arterial adventitia using confocal laser scanning microscopy. Biomech Model Mechanobiol, 2012. 11(3-4): p. 461–73.

105. Sanjana, N.E., O. Shalem, and F. Zhang, Improved vectors and genome-wide libraries for CRISPR screening. Nat Methods, 2014. 11(8): p. 783–4.

106. Thakore, P.I., et al., Highly specific epigenome editing by CRISPR-Cas9 repressors for silencing of distal regulatory elements. Nat Methods, 2015. 12(12): p. 1143–9.

107. Horlbeck, M.A., et al., Compact and highly active next-generation libraries for CRISPR-mediated gene repression and activation. Elife, 2016. 5.

108. Kim, D., et al., TopHat2: accurate alignment of transcriptomes in the presence of insertions, deletions and gene fusions. Genome Biol, 2013. 14(4): p. R36.

109. Anders, S., P.T. Pyl, and W. Huber, HTSeq--a Python framework to work with high-throughput sequencing data. Bioinformatics, 2015. 31(2): p. 166–9.

110. Love, M.I., W. Huber, and S. Anders, Moderated estimation of fold change and dispersion for RNA-seq data with DESeq2. Genome Biol, 2014. 15(12): p. 550.

111. Subramanian, A., et al., Gene set enrichment analysis: a knowledge-based approach for interpreting genome-wide expression profiles. Proc Natl Acad Sci U S A, 2005. 102(43): p. 15545–50.

112. Hanzelmann, S., R. Castelo, and J. Guinney, GSVA: gene set variation analysis for microarray and RNA-seq data. BMC Bioinformatics, 2013. 14: p. 7.

113. Liberzon, A., et al., The Molecular Signatures Database (MSigDB) hallmark gene set collection. Cell Syst, 2015. 1(6): p. 417–425.

114. Nagy, A., et al., Validation of miRNA prognostic power in hepatocellular carcinoma using expression data of independent datasets. Sci Rep, 2018. 8(1): p. 9227.

115. Liu, J., et al., An Integrated TCGA Pan-Cancer Clinical Data Resource to Drive High-Quality Survival Outcome Analytics. Cell, 2018. 173(2): p. 400–416 e11.

