## Supplementary material for "Netrin G1 promotes pancreatic tumorigenesis through cancer associated fibroblast driven nutritional support and immunosuppression": Sup Tables 7-9

**Supplemental Table 4.** Concentration of amino acids in conditioned media and lysate of Normal PSCs

| Amino Acid | Media Concentration  (µM) | p-value  vs. CON CAF | p-value  vs. NetG1 KO CAF | Lysate Concentration (nmol/mg lysate) | p-value  vs. CON CAF Lysate | p-value  vs. NetG1 KO CAF Lysate |
| --- | --- | --- | --- | --- | --- | --- |
| Alanine | 96.22 ± 0.91 | ***8.26E-14*** | ***4.05E-10*** | 27.47 ± 3.71 | ***1.51E-06*** | ***0.00587*** |
| Arginine | 24.17 ± 2.06 | ***4.35E-06*** | ***8.21E-08*** | 6.17 ± 1.00 | ***7.21E-04*** | 0.331 |
| Cysteine | 7.48 ± 1.04 | ***7.41E-16*** | ***2.23E-14*** | 1.37 ± 1.57 | ***7.93E-05*** | ***2.76E-05*** |
| Cystine | -36.13 ± 0.80 | ***8.88E-05*** | ***1.28E-08*** | BD | N/A | N/A |
| Glutamate | 64.57 ± 0.75 | ***6.43E-14*** | ***3.06E-08*** | 191.98 ± 25.82 | 0.259 | 0.0894 |
| Glutamine | 12.70 ± 0.26 | ***2.06E-10*** | ***2.11E-08*** | BD | N/A | N/A |
| Glycine | 253.42 ± 20.40 | 0.717 | ***3.01E-04*** | 348.81 ± 48.90 | ***0.0123*** | 0.222 |
| Histidine | 12.15 ± 1.03 | ***8.52E-06*** | ***1.89E-07*** | 14.01 ± 1.87 | 0.418 | ***0.00126*** |
| Isoleucine | -73.70 ± 4.11 | ***3.19E-11*** | ***5.02E-11*** | 23.27 ± 4.10 | ***8.42E-04*** | 0.353 |
| Leucine | -72.25 ± 3.91 | ***3.05E-11*** | ***1.05E-11*** | 36.98 ± 2.36 | 0.200 | ***1.53E-05*** |
| Lysine | 51.55 ± 4.72 | ***1.36E-07*** | ***6.95E-06*** | 9.73 ± 1.68 | ***0.00696*** | 0.794 |
| Methionine | 13.25 ± 1.33 | ***2.27E-06*** | ***5.46E-10*** | 14.10 ± 2.18 | 0.169 | ***9.95E-05*** |
| Phenylalanine | 28.68 ± 2.65 | ***5.96E-06*** | ***2.45E-07*** | 32.23 ± 4.25 | 0.635 | ***0.00284*** |
| Proline | 36.73 ± 1.00 | ***5.81E-07*** | 0.392 | 52.52 ± 11/48 | ***0.00617*** | ***7.51E-04*** |
| Serine | -214.50 ± 0.84 | ***1.88E-17*** | ***5.34E-18*** | 28.94 ± 4.22 | ***2.86E-06*** | ***6.58E-06*** |
| Threonine | 44.03 ± 4.15 | ***4.25E-06*** | ***6.06E-08*** | 170.76 ± 24.54 | ***3.22E-05*** | ***0.00598*** |
| Tyrosine | 26.60 ± 2.27 | ***3.83E-07*** | ***4.42E-08*** | 31.79 ± 4.42 | 0.545 | ***4.36E-04*** |
| Valine | 20.80 ± 4.22 | ***8.21E-10*** | ***6.28E-10*** | 29.59 ± 4.45 | 0.454 | 0.724 |

Media concentration: (mean ± SE) was calculated as the net gain, signifying the amount of amino acid present in the media after blank media subtraction. Lysate Concentration: (mean ± SE) was calculated as the amount of amino acid present normalized to total protein concentration of the lysate. Student’s t-Test comparing normal PSC with CON CAF or NetG1 KO CAF. Values in bold represent statistical significance (p<0.05). BD: below detection. N/A: not applicable.

**Supplemental Table 5.** Concentration of amino acids in conditioned media and lysate of CON CAFs.

| Amino Acid | Media Concentration  (µM) | p-Value vs. Normal CM | p-value  vs. NetG1 KO CAF CM | Lysate Concentration (nmol/mg lysate) | p-Value vs. Normal Lysate | p-value  vs. NetG1 KO CAF Lysate |
| --- | --- | --- | --- | --- | --- | --- |
| Alanine | 34.32 ± 0.63 | ***8.26E-14*** | ***7.02E-12*** | 11.86 ± 0.82 | ***1.51E-06*** | ***2.08E-06*** |
| Arginine | -8.63 ± 3.03 | ***4.35E-06*** | ***0.000472*** | 8.58 ± 0.71 | ***7.21E-04*** | ***0.00521*** |
| Cysteine | 25.52 ± 2.67 | ***7.41E-16*** | 0.127 | 8.03 ± 2.03 | ***7.93E-05*** | ***0.00556*** |
| Cystine | -121.35 ± 0.51 | ***8.88E-05*** | ***1.51E-09*** | BD | N/A | N/A |
| Glutamate | 137.12 ± 1.02 | ***6.43E-14*** | ***6.48E-15*** | 177.58 ± 14.16 | 0.259 | ***0.00255*** |
| Glutamine | 25.05 ± 0.41 | ***2.06E-10*** | ***1.08E-11*** | BD | N/A | N/A |
| Glycine | 245.62 ± 4.68 | 0.717 | ***6.12E-08*** | 282.63 ± 22.21 | ***0.0123*** | ***0.0369*** |
| Histidine | -3.28 ± 1.55 | ***8.52E-06*** | ***0.00494*** | 14.80 ± 1.34 | 0.418 | ***0.00176*** |
| Isoleucine | -259.00 ± 4.43 | ***3.19E-11*** | 0.510 | 15.09 ± 1.35 | ***8.42E-04*** | ***5.53E-04*** |
| Leucine | -253.27 ± 4.39 | ***3.05E-11*** | ***0.00131*** | 38.61 ± 1.68 | 0.200 | ***2.00E-05*** |
| Lysine | -49.25 ± 6.14 | ***1.36E-07*** | ***0.0152*** | 12.42 ± 0.99 | ***0.00696*** | ***3.91E-04*** |
| Methionine | -10.00 ± 2.02 | ***2.27E-06*** | ***1.18E-05*** | 12.65 ± 0.96 | 0.169 | ***7.67E-06*** |
| Phenylalanine | -12.22 ± 3.92 | ***5.96E-06*** | ***0.0149*** | 33.27 ± 2.99 | 0.635 | ***0.00223*** |
| Proline | 52.18 ± 0.96 | ***5.81E-07*** | ***1.48E-05*** | 33.85 ±6.58 | ***0.00617*** | 0.138 |
| Serine | -347.70 ± 0.60 | ***1.88E-17*** | 0.0827 | 12.09 ± 1.25 | ***2.86E-06*** | ***0.0188*** |
| Threonine | -23.03 ± 6.22 | ***4.25E-06*** | ***0.000863*** | 95.59 ± 8.15 | ***3.22E-05*** | ***5.95E-05*** |
| Tyrosine | -18.80 ± 3.17 | ***3.83E-07*** | ***0.0215*** | 33.25 ± 3.64 | 0.545 | ***5.35E-04*** |
| Valine | -132.52 ± 5.52 | ***8.21E-10*** | 0.524 | 27.98 ± 2.37 | 0.454 | 0.204 |

Media concentration: (mean ± SE) was calculated as the net gain, signifying the amount of amino acid present in the media after blank media subtraction. Lysate Concentration: (mean ± SE) was calculated as the amount of amino acid present normalized to total protein concentration of the lysate. Student’s t-Test comparing CON CAF with normal PSC or NetG1 KO CAF. Values in bold represent statistical significance (p<0.05). BD: below detection. N/A: not applicable.

**Supplemental Table 6.** Concentration of amino acids in conditioned media and lysate of NetG1 KO CAFs.

| Amino Acid | Media Concentration  (µM) | p-Value vs. Normal CM | p-value  vs. CAF CM | Lysate Concentration (nmol/mg lysate) | p-Value vs. Normal Lysate | p-value  vs. CAF  Lysate |
| --- | --- | --- | --- | --- | --- | --- |
| Alanine | 68.57 ± 0.72 | ***4.05E-10*** | ***7.02E-12*** | 21.31 ± 2.24 | ***0.00587*** | ***2.08E-06*** |
| Arginine | -32.48 ± 3.58 | ***8.21E-08*** | ***0.000472*** | 6.77 ± 1.03 | 0.331 | ***0.00521*** |
| Cysteine | 30.20 ± 0.89 | ***2.23E-14*** | 0.127 | 14.77 ± 4.25 | ***2.76E-05*** | ***0.00556*** |
| Cystine | -103.50 ± 0.69 | ***1.28E-08*** | ***1.51E-09*** | BD | N/A | N/A |
| Glutamate | 49.18 ± 0.67 | ***3.06E-08*** | ***6.48E-15*** | 216.81 ± 19.47 | 0.0894 | ***0.00255*** |
| Glutamine | 4.70 ± 0.43 | ***2.11E-08*** | ***1.08E-11*** | BD | N/A | N/A |
| Glycine | 138.68 ± 5.93 | ***3.01E-04*** | ***6.12E-08*** | 318.53 ± 29.00 | 0.222 | ***0.0369*** |
| Histidine | -11.10 ± 1.53 | ***1.89E-07*** | ***0.00494*** | 19.09 ± 2.09 | ***0.00126*** | ***0.00176*** |
| Isoleucine | -263.57 ± 5.01 | ***5.02E-11*** | 0.510 | 21.39 ± 2.79 | 0.353 | ***5.53E-04*** |
| Leucine | -281.58 ± 4.69 | ***1.05E-11*** | ***0.00131*** | 48.06 ± 2.58 | ***1.53E-05*** | ***2.00E-05*** |
| Lysine | -21.57 ± 7.20 | ***6.95E-06*** | ***0.0152*** | 9.52 ± 0.94 | 0.794 | ***3.91E-04*** |
| Methionine | -29.15 ± 1.28 | ***5.46E-10*** | ***1.18E-05*** | 23.60 ± 3.04 | ***9.95E-05*** | ***7.67E-06*** |
| Phenylalanine | -28.32 ± 3.83 | ***2.45E-07*** | ***0.0149*** | 6.77 ± 1.36 | ***0.00284*** | ***0.00223*** |
| Proline | 34.72 ± 2.02 | 0.392 | ***1.48E-05*** | 28.72 ± 4.15 | ***7.51E-04*** | 0.138 |
| Serine | -346.38 ± 0.33 | ***5.34E-18*** | 0.0827 | 13.87 ± 0.92 | ***6.58E-06*** | ***0.0188*** |
| Threonine | -65.17 ± 6.50 | ***6.06E-08*** | ***0.000863*** | 132.61 ± 11.02 | ***0.00598*** | ***5.95E-05*** |
| Tyrosine | -31.10 ± 3.22 | ***4.42E-08*** | ***0.0215*** | 44.34 ± 4.03 | ***4.36E-04*** | ***5.35E-04*** |
| Valine | -137.70 ± 5.57 | ***6.28E-10*** | 0.524 | 30.45 ± 3.76 | 0.724 | 0.204 |

Media concentration: (mean ± SE) was calculated as the net gain, signifying the amount of amino acid present in the media after blank media subtraction. Lysate Concentration: (mean ± SE) was calculated as the amount of amino acid present normalized to total protein concentration of the lysate. Student’s t-Test comparing NetG1 KO CAF with normal PSC or CON CAF. Values in bold represent statistical significance (p<0.05). BD: below detection. N/A: not applicable.
