## Supplementary material for "Netrin G1 promotes pancreatic tumorigenesis through cancer associated fibroblast driven nutritional support and immunosuppression": Sup Table 10

**Supplementary Table 7.** Concentration of metabolites (Net Gain) in the conditioned media of various fibroblast lines

| **Metabolite** | **Normal**  **PSC** | **CON**  **CAF** | **NetG1 KO CAF** | **NetG1**  **KD 1 CAF** | **NetG1**  **KD 2 CAF** | **VGlut1 KD 1 CAF** | **VGlut1**  **KD 2 CAF** |
| --- | --- | --- | --- | --- | --- | --- | --- |
| Alanine | 28.35 ± 2.437  ***N/A***  ***###*** | 36.55 ± 0.57  *********  ***N/A*** | 7.813 ± 0.06473  **********  ***####*** | 13.36 ± 0.2513  **********  ***####*** | 11.6 ± 0.4475  **********  ***####*** | 13.3 ± 0.3105  **********  ***####*** | 10.13 ± 0.2023  **********  ***####*** |
| Alpha Ketoglutarate | 0.8457 ±  0.06361  ***N/A***  ***####*** | 5.094 ± 0.08205  **********  ***N/A*** | 1.022 ± 0.002391  ***NS***  ***####*** | 1.592 ± 0.02504  **********  ***####*** | 1.094 ± 0.06425  *******  ***####*** | 1.486 ±  0.02299  **********  ***####*** | 1.235 ± 0.02687  *********  ***####*** |
| Asparagine | 1.704 ±  0.122  ***N/A***  ***####*** | 2.911 ±  0.05682  **********  ***N/A*** | 1.004 ±  0.006507  **********  ***####*** | 0.8863 ±  0.04785  **********  ***####*** | 1.008 ±  0.02226  **********  ***####*** | 0.8297 ± 0.03761  **********  ***####*** | 0.512 ±  0.04849  **********  ***####*** |
| Cystine | 197.8 ± 9.045  ***N/A***  ***####*** | 17.74 ± 0.3624  **********  ***N/A*** | 3.923 ± 0.06942  **********  ***NS*** | 11.89 ± 2.064  **********  ***NS*** | 6.891 ± 0.182  **********  ***NS*** | 6.692 ± 0.2800  **********  ***NS*** | 10.11 ± 0.2522  **********  ***NS*** |
| Glutamate | 62.92 ± 7.366  ***N/A***  ***NS*** | 74.76 ± 4.256  ***NS***  ***N/A*** | 17.02 ± 0.1554  **********  ***####*** | 27.63 ± 0.3697  **********  ***####*** | 29.82 ±  0.9573  **********  ***####*** | 27.98 ± 0.9761  **********  ***####*** | 28.53 ± 0.8144  **********  ***####*** |
| Glutamine | 5.505 ± 0.569  ***N/A***  ***####*** | 12.66 ± 0.1578  ****  ***N/A*** | 6.635 ± 0.04415  ***NS***  ***####*** | 7.7 ±  0.3491  *********  ***####*** | 5.464 ±  0.2617  ***NS***  ***####*** | 7.901 ±  0.2921  *********  ***####*** | 3.795 ± 0.06163  ********  ***####*** |
| Glycine | 96.72 ± 22.65  ***N/A***  ***##*** | 49.41 ± 3.173  ********  ***N/A*** | 13.46 ± 0.1306  **********  ***#*** | 18.71 ± 0.7797  *********  ***NS*** | 20.3 ± 2.093  *********  ***NS*** | 17.05 ± 1.639  **********  ***NS*** | 33.66 ± 0.6189  *********  ***NS*** |
| Isoleucine | 88.66 ± 68.68  ***N/A***  ***###*** | -126 ± 5.726  *********  ***N/A*** | -16.98 ± 0.4396  ***NS***  ***#*** | -30.32 ± 0.7639  *******  ***NS*** | -20.87 ± 1.983  *******  ***NS*** | -29.11 ± 2.851  *******  ***NS*** | -29.95 ± 1.813  *******  ***NS*** |
| Lactate | 3509 ± 251.1  ***N/A***  ***####*** | 5187 ± 93.17  **********  ***N/A*** | 927.1 ± 9.604  **********  ***####*** | 2071 ± 36.45  **********  ***####*** | 1579 ± 64.38  **********  ***####*** | 1485 ± 21.41  **********  ***####*** | 1749 ±  41.8  **********  ***####*** |
| Leucine | 43.94 ± 41.18  ***N/A***  ***###*** | -89.04 ± 4.347  *********  ***N/A*** | -10.67 ± 0.1996  ***NS***  ***#*** | -20.88 ± 0.8872  *******  ***#*** | -12.4 ± 1.54  ***NS***  ***#*** | -20.2 ± 1.843  ***NS***  ***#*** | -22.06 ± 1.075  *******  ***#*** |
| Methionine | 22.51 ± 11.4  ***N/A***  ***##*** | -0.8776 ± 1.348  ********  ***N/A*** | 1.287 ± 0.08895  *******  ***NS*** | 1.624 ± 0.3146  *******  ***NS*** | 2.465 ± 0.5353  *******  ***NS*** | 1.985 ± 0.6482  *******  ***NS*** | 1.127 ± 0.2662  *******  ***NS*** |
| Phenylalanine | 51.53 ± 23.46  ***N/A***  ***##*** | 2.349 ± 2.793  ********  ***N/A*** | 2.6 ± 0.1689  ********  ***NS*** | 2.335 ± 0.3539  ********  ***NS*** | 4.182 ± 1.196  ********  ***NS*** | 3.707 ± 1.326  ********  ***NS*** | 3.23 ± 0.9012  ********  ***NS*** |
| Proline | 11.46 ± 0.9729  ***N/A***  ***####*** | 34.56 ± 0.5586  **********  ***N/A*** | 6.316 ± 0.1315  **********  ***####*** | 8.74 ± 0.2204  ********  ***####*** | 11.1 ± 0.4035  ***NS***  ***####*** | 7.876 ± 0.212  **********  ***####*** | 5.234 ± 0.0368  **********  ***####*** |
| Serine | -46.47 ± 17.51  ***N/A***  ***####*** | -107 ± 0.4106  **********  ***N/A*** | -12.63 ± 0.2324  *******  ***####*** | -24.02 ± 0.3876  ***NS***  ***####*** | -19.99 ± 0.3561  ***NS***  ***####*** | -23.81 ± 0.8496  ***NS***  ***####*** | -36.89 ± 0.5659  ***NS***  ***####*** |
| Threonine | 93.33 46.52  ***N/A***  ***#*** | -0.6362 ± 4.047  *******  ***N/A*** | 3.275 ± 0.7523  *******  ***#*** | 4.309 ± 0.7000  *******  ***#*** | 7.112 ± 1.485  *******  ***#*** | 5.644 ± 2.329  *******  ***#*** | 5.105 ± 1.767  *******  ***#*** |
| Tyrosine | 15.8 ± 22.14  ***N/A***  ***NS*** | -3.832 ± 2.567  ***NS***  ***N/A*** | 0.7498 ± 0.1584  ***NS***  ***NS*** | -1.2 ± 1.195  ***NS***  ***NS*** | 1.71 ± 1.078  ***NS***  ***NS*** | 0.831 ± 0.8365  ***NS***  ***NS*** | -0.1625 ± 0.7787  ***NS***  ***NS*** |
| Valine | 66.87 ± 42.92  ***N/A***  ***##*** | -41.11 ± 5.187  ********  ***N/A*** | -3.147 ± 0.3228  ***NS***  ***NS*** | -6.109 ± 0.9047  ***NS***  ***NS*** | -2.677 ± 1.658  ***NS***  ***NS*** | -5.893 ± 2.000  ***NS***  ***NS*** | -5.618 ± 1.298  ***NS***  ***NS*** |

Concentration (µM) of metabolites (net gain, subtracted from unconditioned media) is displayed as mean ± standard error. One-way ANOVA was performed to determine statistically significant differences in metabolite concentrations between cell lines. ***** Compared to Normal PSCs. **#** Compared to Control (CON) CAFs.
