## Supplementary material for "Netrin G1 promotes pancreatic tumorigenesis through cancer associated fibroblast driven nutritional support and immunosuppression": Key Resources Table

| **Antibodies** |  |  |
| --- | --- | --- |
| **Reagent or Resource** | **Source** | **Identifier** |
| Netrin G1 Antibody (D-2) (1:100) | Santa Cruz | RRID:AB_10707668 |
| Netrin G1 Antibody (N2C1)  (1:100) | Genetex | RRID:AB_10625511 |
| Glutaminase antibody [EP7212] (1:100,000) | Abcam | #ab156876 |
| Glutamine Synthetase antibody (1:50,000) | Abcam | #ab73593 |
| Histone H3 (D1H2) XP® antibody (1:10,000) | Cell Signaling Technology | #4499 |
| EpCam (MOC-31) antibody (1:200) | Novus Biologicals | #NBP2-22388 |
| pan-cytokeratin (AE1/AE3) antibodies (1:200) | DAKO | #M3515 |
| CD-70 antibody (1:200) | Biolegend | RRID:AB_2561429 |
| Vimentin antibody (1:200) | Abcam | #ab92547 |
| GAPDH antibody (1:10,000) | Abcam | RRID:AB_307275 |
| VGlut1 antibody (1:100) | Thermo Fisher Scientific | RRID:AB_2533843 |
| Fibronectin antibody (1:200) | Sigma-Aldrich | RRID:AB_476961 |
| αSMA antibody (1:300) | Sigma-Aldrich | RRID: AB_476701 |
| SNAKA51 Antibody (45 µg/mL) | M Humphries | N/A |
| p-FAK [Y397] Antibody (1:200) | Thermo Fisher Scientific | RRID:AB_2533701 |
| Ki67 Antibody (1:500) | Abcam | RRID:AB_443209 |
| Granzyme B Antibody (1:50) | ebioscence | RRID:AB_1659718 |
| IFNγ secretion assay detection kit-PE | Miltenyi Biotec | #130-054-20 |
| NKp80-APC | Biolegend | RRID:AB_2044041 |
| CD56-BUV395 | BD Horizon | RRID:AB_2687886 |
| CD69-Pac Blue | Biolegend | RRID:AB_493667 |
| CD3-Cy7APC | BD Pharmigen | RRID:AB_1645475 |
| donkey anti-rabbit secondary antibody- Cy5 conjugation (1:200) | Jackson ImmunoResearch | RRID:AB_2340607 |
| donkey anti-mouse secondary antibody- TRITC conjugation (1:200) | Jackson ImmunoResearch | RRID:AB_2340766 |
| Human IL-15 Antibody (10 µg/mL) | R&D Systems | RRID:AB_2124577 |
| Anti-Mouse IgG (whole molecule)–Peroxidase antibody (1:5,000) | Sigma-Aldrich | RRID:AB_258431 |
| Goat anti-Rabbit IgG (H+L) Secondary Antibody, HRP (1:10,000) | Thermo Fisher Scientific | RRID:AB_2533967 |
| p-p38 (Thr180/Tyr182) (1:250) | Cell Signaling Technology | RRID: AB_2139682 |
| p38 (1:2000) | Cell Signaling Technology | RRID: AB_10999090 |
| p-AKT (Ser473) (1:1000) | Cell Signaling Technology | #4060 |
| Pan AKT (1:2500) | Cell Signaling Technology | RRID: AB_915783 |
| p-4E-BP1 (Thr37/46) (1:1000) | Cell Signaling Technology | RRID: AB_560835 |
| 4E-BP1 (1:2000) | Cell Signaling Technology | RRID: AB_2097841 |
| FRA1 (1:2000) | Cell Signaling Technology | RRID: AB_10557418 |
| FRA2 (1:2000) | Cell Signaling Technology | RRID: AB_2722526 |
| FosB | Cell Signaling Technology | RRID: AB_2106903 |
| cFOS | Cell Signaling Technology | RRID: AB_2247211 |
| c-Jun | Cell Signaling Technology | RRID: AB_2130165 |
| JunB | Cell Signaling Technology | RRID: AB_2130002 |
| p-mTOR (Ser2448) | Cell Signaling Technology | #5536 |
| p-FOXO1 (Ser256) | Cell Signaling Technology | RRID: AB_2800035 |
| FOXO1 | Cell Signaling Technology | RRID: AB_2106495 |
| p-FOXO3 (Ser294) | Cell Signaling Technology | #5538 |
| FOXO3 | Cell Signaling Technology | RRID: AB_836876 |
| p-GSK3β (Ser9) | Cell Signaling Technology | #9323 |
| Mouse IgG1 | Bio X Cell | #BE-0083 |
| **Bacterial Strains** |  |  |
| **Reagent or Resource** | **Source** | **Identifier** |
| One Shot Stbl3 Chemically Competent E. coli | Thermo Fisher Scientific | # C737303 |
| **Chemicals** |  |  |
| **Reagent or Resource** | **Source** | **Identifier** |
| L-glutamate | Abcam | #ab120049 |
| MK-2206 | Cayman Chemical | #11593 |
| SB 202190 | Tocris Bioscience | #1264 |
| Sulfosalicylic acid | Sigma-Aldrich | #S-7422 |
| L-Ascorbic acid | Sigma-Aldrich | #A92902-100G |
| 25% Glutaraldehyde Solution in water | Sigma-Aldrich | #G6257-1L |
| Ethanolamine | Sigma-Aldrich | #E9508-1L |
| Polybrene | Santa Cruz Biotechnology | # sc-134220 |
| L-Methionine sulfoximine (MSO) | Sigma-Aldrich | #M5379-250MG |
| isotopically labeled amino acids | Cambridge Isotope Laboratories | #MSK-A2-1.2 |
| MOX reagent | Thermofisher | # TS-45950 |
| N-tertbutyldimethylsilyl-N-methyltrifluoroacetamide with 1% tert-butyldimethylchlorosilane | Sigma-Aldrich | # 375934 |
| **Cell Culture** |  |  |
| **Reagent or Resource** | **Source** | **Identifier** |
| FBS | Atlanta Biologicals | #S11150 |
| DMEM | Corning | #50-013-PB |
| L-Glutamine | Corning | #25-005CI |
| Penicillin/Streptomycin | Corning | #30-002-CI |
| DMEM low glucose | Corning | #10-014 |
| M3 Base F | INCELL | #M300F-500 |
| Alpha MEM | Corning | # 10-022-CV |
| IL-2 | Fox Chase Cancer Center | N/A |
| IL-12 | Fox Chase Cancer Center | N/A |
| **Cell Lines** |  |  |
| **Reagent or Resource** | **Source** | **Identifier** |
| hTERT-HPNE E6/E7/K-RasG12D (*KRAS*) | ATCC | RRID:CVCL_C469 |
| hTERT-HPNE (*hTERT*) | ATCC | RRID:CVCL_C466 |
| hTERT-HPNE E6/E7 (*E6/E7*) | ATCC | RRID:CVCL_C467 |
| KPC3 | This paper | N/A |
| KPC4B | This paper | N/A |
| Panc-1 | ATCC | RRID:CVCL_0480 |
| Cancer associated fibroblasts (patient derived) | This Paper | N/A |
| Normal PSCs (patient derived) | This Paper | N/A |
| NK-92 cells | ATCC | RRID:CVCL_2142 |
| 293T Cells | ATCC | RRID:CVCL_0063 |
| Phoenix-Amphotropic (φNX) | ATCC | RRID:CVCL_H716 |
| Murine pancreatic fibroblasts | This paper | N/A |
| **Organisms/Strains (Mice)** |  |  |
| **Reagent or Resource** | **Source** | **Identifier** |
| C57BL/6 | Fox Chase Cancer Center | N/A |
| KPC model: LSL-Kras(G12D/+) ; Trp53(flox/flox) ;Pdx-1-Cre | Kerry Campbell, Igor Astsaturov | Original Mice: PMID: 15894267 |
| KC Model: LSL-Kras(G12D/+); Pdx-1-Cre | Kerry Campbell | Original Mice: PMID: 14706336 |
| C.B17 scid | Taconic Bioscience | #CB17SC |
| Nod SCID Gamma (NSG) | Jackson Laboratories | #005557 |
| **Plasmids** |  |  |
| **Reagent or Resource** | **Source** | **Identifier** |
| Plv-Cmv-Puro | Fox Chase Cancer Center | N/A |
| LentiCRISPR v2 | Addgene  (Sanjana et al., 2014) | #52961 |
| psPAX2 | Addgene | #12260 |
| pCMV-VSV-G | Addgene | #8454 |
| pBABE-neo-hTERT | Addgene | #1774 |
| CRISPRi-Puro | Modified from Addgene;  (Thakore et al., 2015) | #71236 |
| **Critical Commercial Reagents** |  |  |
| **Reagent or Resource** | **Source** | **Identifier** |
| PureLink™ RNA Mini Kit | Thermo Fisher Scientific | #12183025 |
| SuperScript III First Strand Synthesis System | Thermo Fisher Scientific | #18080-051 |
| Phusion High-Fidelity PCR Master Mix | Thermo Fisher Scientific | # F-531L |
| 2x Laemmli buffer | BioRad | #1610737 |
| Sytox Blue | Thermo Fisher Scientific | #S11348 |
| Odyssey Blocking Buffer (PBS) | LI-COR | #927-40000 |
| DeadEnd™ Fluorometric TUNEL System | Promega | #G3250 |
| Immobilon Western Chemiluminescent HRP Substrate | EMD Millipore | #WBKLS0500 |
| Ultrafree-MC-GV centrifugal filters | Millipore | #UFC30GVNB |
| Dnase I | Thermo Fisher Scientific | #EN0525 |
| X-tremeGene9 | Sigma-Aldrich | #6365787001 |
| U-PLEX Biomarker Group 1 (hu) Assays, SECTOR (1 PL) | Meso Scale Discoveries | #K15067L-1 |
| DuoSet ELISAs (IL-15, IFN-β, TGF-β, IL-6, IL-8, GM-CSF, mTGF-β, mGM-CSF, mIL-6) | R&D Systems | #DY247-05, #DY814-05, #DY240-05, #DY206-05, #DY208-05, #DY215-05, #DY1679-05, #DY415-05, # DY406-05 |
| DuoSet ELISA Ancillary Reagent Kit 1 | R&D Systems | DY007 |
| DuoSet ELISA Ancillary Reagent Kit 2 | R&D Systems | DY008 |
| Sample Activation Kit 1 | R&D Systems | DY010 |
| Immobilon-P PVDF Membrane | EMD Millipore | IPVH00010 |
| SYBR Green | Invitrogen | S7563 |
| Collagenase/Hyaluronidase | StemCell Technologies | #07912 |
| Dispase | StemCell Technologies | #7446 |
| Magnevist | Bayer | 50419-188-58 |
| SiteClick Qdot labeling Kits (525-655) | Thermo Fisher Scientific | #S10449, S10450, S10451, S10469, S10452, S10453 |
| Truseq stranded mRNA library kit | Illumina, Inc. | # 20020595 |
| SuperscriptII reverse transcriptase | Thermo Fisher Scientific | #18064014 |
| SPRIselect beads | Beckman Coulter | # B23318 |
| HiSeq rapid SBS kit v2 (50 cycle) | Illumina, Inc. | # FC-402-4022 |
| Qubit 1x dsDNA HS assay kit | Thermo Fisher Scientific | #Q33216 |
| High sensitivity DNA kit | Agilent Technologies | # 5067-4626 |
| Glutamate Assay Kit | Abcam | # ab83389 |
| Glutamine ELISA Kit | MyBioSource | # MBS732543 |
| High Capacity cDNA Reverse Transcription Kit | Thermo Fisher Scientific | #4368814 |
| PowerSYBR Green PCR Master Mix | Thermo Fisher Scientific | #4367659 |
| MicroAmp Fast Optical 96 well Reaction Plate with Barcode (0.1 mL) | Thermo Fisher Scientific | #4346906 |
| **Oligonucleotides** |  |  |
| **Reagent or Resource** | **Source** | **Identifier** |
| GFP Xba FW: TAGGTATCTAGAATGGTGAGCAAGGGCGAGG | This paper | N/A |
| GFP Xho RV: TAGGTACTCGAGCTTGTACAGCTCGTCCATGCC | This paper | N/A |
| mCherry Xba FW: TAGGTATCTAGAatggtgagcaagggcgag | This paper | N/A |
| mCherry Xho RV: TAGGTACTCGAGttacttgtacagctcgtccatgc | This paper | N/A |
| NTNG1 qPCR FW: GGAAATGCAAGAAGAATTATCAGG | This paper | N/A |
| NTNG1 qPCR RV: GTTGTCGCAGACATTCGTACC | This paper | N/A |
| POLR2F qPCR internal control: 6fam-CCCCATCATCATTCGCCGTTACC-bqh1 | This paper | N/A |
| eGFP gRNA 1.1: CACCGCATGTGATCGCGCTTCTCGT | This paper | N/A |
| eGFP gRNA 1.2 : AAACACGAGAAGCGCGATCACATGC | This paper | N/A |
| NetrinG1 gRNA 1.1: CACCGGCGTCCAGACCAAATGATCC | This paper | N/A |
| NetrinG1 gRNA 1.2: AAACGGATCATTTGGTCTGGACGCC | This paper | N/A |
| NetrinG1 gRNA 2.1: CACCGTGATGCGAGTACCCCTGAGC | This paper | N/A |
| NetrinG1 gRNA 2.2: AAACGCTCAGGGGTACTCGCATCAC | This paper | N/A |
| NGL1 gRNA1.1: CACCGAACCTGCGTGAGGTTCCGGA | This paper | N/A |
| NGL1 gRNA1.2: AAACTCCGGAACCTCACGCAGGTTC | This paper | N/A |
| NGL1 gRNA 2.1: CACCGTGCCATCCGGAACCTCACGC | This paper | N/A |
| NGL1 gRNA 2.2: AAACGCGTGAGGTTCCGGATGGCAC | This paper | N/A |
| murine NGL1 gRNA 1.1: CACCGCGATTTCAATGGTTCGAATA | This paper | N/A |
| murine NGL1 gRNA 1.2: AAACTATTCGAACCATTGAAATCGC | This paper | N/A |
| murine NGL1 gRNA 2.1: CACCGAACCTTCGTGAAGTTCCGGA | This paper | N/A |
| murine NGL1 gRNA 2.2: AAACTCCGGAACTTCACGAAGGTTC | This paper | N/A |
| NetrinG1 CRISPRi gRNA 1.1: CACCGGCGCCCCGAGGTCGTGGAG | (Horlbeck et al., 2016 | N/A |
| NetrinG1 CRISPRi gRNA 1.2: *AAAC*CTCCACGACCTCGGGGCGCC*C* | (Horlbeck et al., 2016 | N/A |
| NetrinG1 CRISPRi gRNA 2.1: *CACC*GACAGCAACAGCGAGCGGGA | (Horlbeck et al., 2016 | N/A |
| NetrinG1 CRISPRi gRNA 2.2: *AAAC*TCCCGCTCGCTGTTGCTGTC*C* | (Horlbeck et al., 2016 | N/A |
| NetrinG1 CRISPRi gRNA 3.1: *CACC*GCGAGAGCCGGAAAGAGGAG | (Horlbeck et al., 2016) | N/A |
| NetrinG1 CRISPRi gRNA 3.2:  *AAAC*CTCCTCTTTCCGGCTCTCGC*C* | (Horlbeck et al., 2016 | N/A |
| VGlut1 CRISPRi gRNA 1.1: *CACCG*AGAGAGAGTGGAGCCGGGT | (Horlbeck et al., 2016 | N/A |
| VGlut1 CRISPRi gRNA 1.2: *AAAC*ACCCGGCTCCACTCTCTCTC*C* | (Horlbeck et al., 2016) | N/A |
| VGlut1 CRISPRi gRNA 2.1: *CACCG*GCTCCGCTCGGGGGGAAGG | (Horlbeck et al., 2016) | N/A |
| VGlut1 CRISPRi gRNA 2.2: *AAAC*CCTTCCCCCCGAGCGGAGCC*C* | (Horlbeck et al., 2016) | N/A |
| VGlut1 CRISPRi gRNA 3.1: *CACCG*AGAGACCCAGAAGTGGTGG | (Horlbeck et al., 2016) | N/A |
| VGlut1 CRISPRi gRNA 3.2: *AAAC*CCACCACTTCTGGGTCTCTC*C* | (Horlbeck et al., 2016) | N/A |
| GS CRISPRi 1.1: CACCGGCTCTGCAGAGTCGAGAGT | (Horlbeck et al., 2016) | N/A |
| GS CRISPRi 1.2: AAACACTCTCGACTCTGCAGAGCCC | (Horlbeck et al., 2016) | N/A |
| GS CRISPRi 2.1: CACCGGAGCGTGTGAGCAGTACTG | (Horlbeck et al., 2016) | N/A |
| GS CRISPRi 2.2: *AAAC*CAGTACTGCTCACACGCTCC*C* | (Horlbeck et al., 2016) | N/A |
| GS CRISPRi 3.1: CACCGGACGGGTCCAAGCCACCAG | (Horlbeck et al., 2016) | N/A |
| GS CRISPRi 3.2: AAACCTGGTGGCTTGGACCCGTCCC | (Horlbeck et al., 2016) | N/A |
| 4E-BP1 CRISPRI 1.1: CACCGCACAGGAGACCATGTCCGG | (Horlbeck et al., 2016) | N/A |
| 4E-BP1 CRISPRI 1.2: AAACCCGGACATGGTCTCCTGTGC | (Horlbeck et al., 2016) | N/A |
| 4E-BP1 CRISPRI 2.1: CACCGAGGGCAGCGAGAGGTTCGC | (Horlbeck et al., 2016) | N/A |
| 4E-BP1 CRISPRI 2.2: AAACGCGAACCTCTCGCTGCCCTC | (Horlbeck et al., 2016) | N/A |
| 4E-BP1 CRISPRI 3.1: CACCGACCGGGTGTCCAGGCTCAA | (Horlbeck et al., 2016) | N/A |
| 4E-BP1 CRISPRI 3.2: AAACTTGAGCCTGGACACCCGGTC | (Horlbeck et al., 2016) | N/A |
| FRA1 CRISPRi 1.1: CACCGAAGTCTCGGAACATGCCCG | (Horlbeck et al., 2016) | N/A |
| FRA1 CRISPRi 1.2: AAACCGGGCATGTTCCGAGACTTC | (Horlbeck et al., 2016) | N/A |
| FRA1 CRISPRi 2.1: CACCGGGCGGCTCTGCGGGGTACA | (Horlbeck et al., 2016) | N/A |
| FRA1 CRISPRi 2.2: AAACTGTACCCCGCAGAGCCGCCC | (Horlbeck et al., 2016) | N/A |
| FRA1 CRISPRi 3.1: CACCGGGCATGTTCCGAGACTTCG | (Horlbeck et al., 2016) | N/A |
| FRA1 CRISPRi 3.2: AAACCGAAGTCTCGGAACATGCCC | (Horlbeck et al., 2016) | N/A |
| VGlut1 qPCR FW: CTGGGGCTACATTGTCACTCA | Primer Bank | N/A |
| VGlut1 qPCR RV: GCAAAGCCGAAAACTCTGTTG | Primer Bank | N/A |
| GS qPCR FW: AAGAGTTGCCTGAGTGGAATTTC | Primer Bank | N/A |
| GS qPCR RV: AGCTTGTTAGGGTCCTTACGG | Primer Bank | N/A |
| IL6 qPCR FW: ACTCACCTCTTCAGAACGAATTG | Primer Bank | N/A |
| IL6 qPCR RV: CCATCTTTGGAAGGTTCAGGTTG | Primer Bank | N/A |
| IL8 qPCR FW: TTTTGCCAAGGAGTGCTAAAGA | Primer Bank | N/A |
| IL8 qPCR RV: AACCCTCTGCACCCAGTTTTC | Primer Bank | N/A |
| IL15 qPCR FW: CAGTGCAGGGCTTCCTAAAAC | Primer Bank | N/A |
| IL15 iso1 qPCR RV: TGGGGTGAACATCACTTTCCG | Primer Bank | N/A |
| TGFB1 qPCR FW: GGCCAGATCCTGTCCAAGC | Primer Bank | N/A |
| TGFB1 qPCR RV: GTGGGTTTCCACCATTAGCAC | Primer Bank | N/A |
| GM-CSF qPCR FW: TCCTGAACCTGAGTAGAGACAC | Primer Bank | N/A |
| GM-CSF qPCR RV: TGCTGCTTGTAGTGGCTGG | Primer Bank | N/A |
| 18s qPCR FW: GGCCCTGTAATTGGAATGAGTC | Primer Bank | N/A |
| 18s qPCR RV: CCAAGATCCAACTACGAGCTT | Primer Bank | N/A |
| **Instruments** |  |  |
| **Reagent or Resource** | **Source** | **Identifier** |
| Agilent Technologies Gene Chip | Agilent | http://www.genomics.agilent.com |
| Agilent Technologies Microarray Instrument | Agilent | http://www.genomics.agilent.com |
| Confocal spinning disk Ultraview | Perkin-Elmer | http://www.perkinelmer.com/ |
| NanoDrop 1000 | Thermo Fisher Scientific | http://www.thermofisherscientific.com |
| FlourChem E Imaging System | Protein Simple | https://www.proteinsimple.com/ |
| Biochrom 30 amino acid analyzer | Biochrom | http://www.biochrom.co.uk/ |
| Nikon Eclipse TE2000U | Nikon | http://www.nikon.com/ |
| MSD SECTOR Imager 2400 | Meso Scale Discoveries | https://www.mesoscale.com |
| Spark® Multimode Microplate | Tecan | https://www.tecan.com |
| CoolSNAP HQ CCD Camera | Photometrics | https://www.photometrics.com/ |
| Transblot SD Semi Dry Transfer System | BioRad | www.bio-rad.com |
| Bruker DRX 300 | Bruker | www.bruker.com |
| DB-35MS column | Agilent Technologies | www.agilent.com |
| Agilent 7890A gas chromatograph | Agilent Technologies | www.agilent.com |
| Agilent 5997B mass spectrometer | Agilent Technologies | www.agilent.com |
| HiSeq2500 System | Illumina, Inc. | www.illumina.com |
| Agilent 2100 bioanalyzer | Agilent Technologies | www.agilent.com |
| Qubit 3.0 flurometer | Thermo Fisher Scientific | http://www.thermofisherscientific.com |
| StepOnePlus Real Time-PCR System | Applied Biosystems/Thermo Fisher Scienfific | https://www.thermofisher.com/order/catalog/product/4376600#/4376600 |
| **Software/Algorithms** |  |  |
| **Reagent or Resource** | **Source** | **Identifier** |
| Agilent Feature Extraction 9.5.1.1 Software | Agilent | http://www.genomics.agilent.com/en/Microarray-Scanner-Processing-Hardware/Feature-Extraction-Software/?cid=AG-PT-144&tabId=AG-PR-1050 (RRID:SCR_014963) |
| ImageJ | NIH | https://imagej.nih.gov/ij/ (RRID:SCR_003070) |
| Metamorph 7.8.1.0 Image Anaylsis Software | Molecular Devices | http://www.moleculardevices.com/Products/Software/Meta-Imaging-Series/MetaMorph.html (RRID:SCR_002368) |
| Volocity 3D Image Analysis Software | Perkin Elmer | http://www.perkinelmer.com/pages/020/cellularimaging/products/volocity.xhtml (RRID:SCR_002668) |
| SMIA-CUKIE | Franco-Barraza et al., 2017 | https://github.com/cukie/SMIA (RRID:SCR_014795) |
| FlowJo | FlowJo LLC | https://www.flowjo.com/solutions/flowjo (RRID:SCR_008520) |
| ParaVision | Bruker | http://www.bruker.com/service/support-upgrades/software-downloads/mri.html (RRID:SCR_001964) |
| OrientationJ plugin for ImageJ | Rezakhaniha et al, 2012 | http://bigwww.epfl.ch/demo/orientation/ (RRID:SCR_014796) |
| MSD Workbench 4.0 software | Meso Scale Discoveries | https://www.mesoscale.com |
| Graph Pad Prism 7.0 software | GraphPad Software | https://www.graphpad.com/scientific-software/prism/ (RRID:SCR_000306) |
| ParaVision | Bruker | https://www.bruker.com (RRID:SCR_001964) |
| Adobe Photoshop CS6 13.0.1 | Adobe | http://www.adobe.com/fr/products/photoshop.html (RRID: SCR_014199) |
| Nuance 3.0.1 | Caliper Life Sciences/Perkin Elmer | https://www.moleculardevices.com/Products/Software/Meta-Imaging-Series/MetaMorph.html (RRID: SCR_002368) |
| Illumina base space | Illumina, Inc. | (https://basespace.illumina.com |
